# The structure of a 2-MDa chloroplast RNA polymerase reveals unexpected evolutionary complexity

**DOI:** 10.64898/2026.06.23.732312

**Authors:** Sandra K. Schuller, Robert Collison, Anuj Kumar, Cheuk-Ling Wun, Carla Brillada, Josef Hoff, Polina N. Foteva, Pamela Vetrano, Julia Kober, Sophie De Vries, Iker Irisarri, Lianyong Wang, Clinton Gabel, Stefan Bohn, Ana Ilieva, Jan De Vries, Jan M. Schuller, Silvia Ramundo

## Abstract

Transcription in chloroplasts depends on the Plastid-Encoded RNA polymerase (PEP), a bacterial-derived enzyme whose catalytic core remains encoded by the highly reduced genome inherited from the cyanobacterial ancestor. In land plants, PEP has roughly doubled in size, expanding into a ∼1 MDa multisubunit machinery through the acquisition of numerous nuclear-encoded subunits. Based on phylogenetic analyses, this added complexity has been widely attributed to the demands of plant terrestrialization. Contrary to this view, we show that in the unicellular green alga *Chlamydomonas reinhardtii*, PEP assembles into an even larger ∼2 MDa complex containing twelve previously uncharacterized nuclear-encoded subunits (PEPS1–12), representing an RNA polymerase architecture of unprecedented size. A cryo-EM structure at 2.7 Å resolution reveals that several of these subunits occupy positions analogous to those in land plant PEP, and that metabolic enzyme folds have been repurposed as structural scaffolds stabilizing the highly expanded plastid-encoded core. Despite this, most of the newly identified PEPS subunits lack detectable sequence or structural similarity to their land plant counterparts. These findings demonstrate that PEP complexity is not a hallmark of land plant evolution and may instead reflect, at least in part, the evolutionary entrenchment of additional subunits around an expanded plastid-encoded core. More broadly, they suggest that essential organellar machines can acquire substantial structural complexity that leaves little trace in sequence-based analyses, a pattern consistent with constructive neutral evolution.

## Introduction

The evolution of chloroplasts from free-living cyanobacteria was a turning point in Earth’s history, enabling the emergence of oxygenic photosynthesis in eukaryotes and reshaping global biogeochemical cycles (*1*, *2*). During this transition, the ancestral cyanobacterial genome—originally containing several thousand genes—was extensively reduced, leaving modern chloroplasts with only about 100 genes (*3–7*). In sharp contrast, the plastid-encoded RNA polymerase (PEP)— the major enzyme responsible for chloroplast gene transcription—evolved into a highly complex molecular machine (*8–11*). In land plants, where PEP has been most extensively studied, this enzyme consists of four core subunits (α, β, β′, β**′′)** encoded by the chloroplast genome and homologous to the eubacterial RNA polymerase subunits α, β, β′. Features such as the split of β′ into β′ (rpoC1) and β′′ (rpoC2) as well as a large insertion in β″ (SI3), underscore its cyanobacterial origin (*12*, *13*). Surrounding this conserved core are at least fourteen different nucleus-encoded PEP-Accessory Proteins (PAPs), which expand the size of the complex to approximately 1MDa. Initially identified through biochemical and genetic studies in several plant species, these proteins colocalize with chloroplast DNA within nucleoid structures (*14–19*), and their loss severely disrupts chloroplast transcription, producing albino or pale-green phenotypes similar to core subunit mutants (*20–24*). Recent structural studies have confirmed that PAPs are integral components of the land plant PEP complex (*8–11*). Moreover, they revealed that PAP12 shares a striking structural similarity with the bacterial ω core subunit, despite significant sequence divergence that had previously led to the assumption that ω was lost from the PEP complex (*25*). Nevertheless, the evolutionary origins of most PAPs and their specific functions are still largely enigmatic. In particular, sequence-based searches fail to identify clear PAP homologs in Chlorophyta (*9*, *26*). Even when potential PAP homologs are found in these species, they typically share less than 30% sequence similarity in interface residues, suggesting they are unlikely to interact with the PEP core subunits in the same way as land plant PAPs (*9*). This phylogenetic gap has fuelled the view that PAPs represent a land plant innovation, perhaps facilitating terrestrial adaptation or enabling tissue-specific plastid differentiation (*9*, *27*, *28*). However, this assumption relies on the untested premise that Chlorophytes harbour a much simpler PEP complex. Here we report the biochemical and structural characterization of PEP from the green alga *Chlamydomonas reinhardtii*, which diverged from land plants ∼1 billion years ago (*29*, *30*). The unexpectedly elaborate PEP assembly in this unicellular photosynthetic eukaryote is compatible with a model in which constructive neutral evolution contributed to a ratchet-like increase in architectural complexity of this essential organellar machine.

## Purification of the CrPEP complex

To enable affinity purification of the Chlamydomonas PEP complex (hereafter CrPEP) in its native state, we used a strain in which a FLAG affinity tag was introduced into the endogenous chloroplast *rpoA* gene (encoding the PEP α subunit) by homologous recombination (Fig. S1) (*31*). Using this engineered strain, we successfully isolated the CrPEP complex through a single-step affinity purification. Following size filtration and sample concentration using a 100 kDa cutoff centrifugal filter, the sample yielded sufficient quality and quantity for biochemical and structural studies (Fig. 1a). Unexpectedly, mass photometry showed that CrPEP forms a complex with an estimated molecular mass of ∼2 MDa. (Fig. 1c). Further gel electrophoresis, followed by Coomassie staining, revealed the presence of several additional proteins alongside the expected cyanobacterial-like core subunits, which account for much of the increased size of the complex (Fig. 1b). Finally, an *in vitro* RNA elongation assay using *E. coli* RNA polymerase as a positive control confirmed that purified CrPEP is transcriptionally active and capable of elongating RNA in the presence of nucleotides (Fig. 1d).

**Figure 1.**
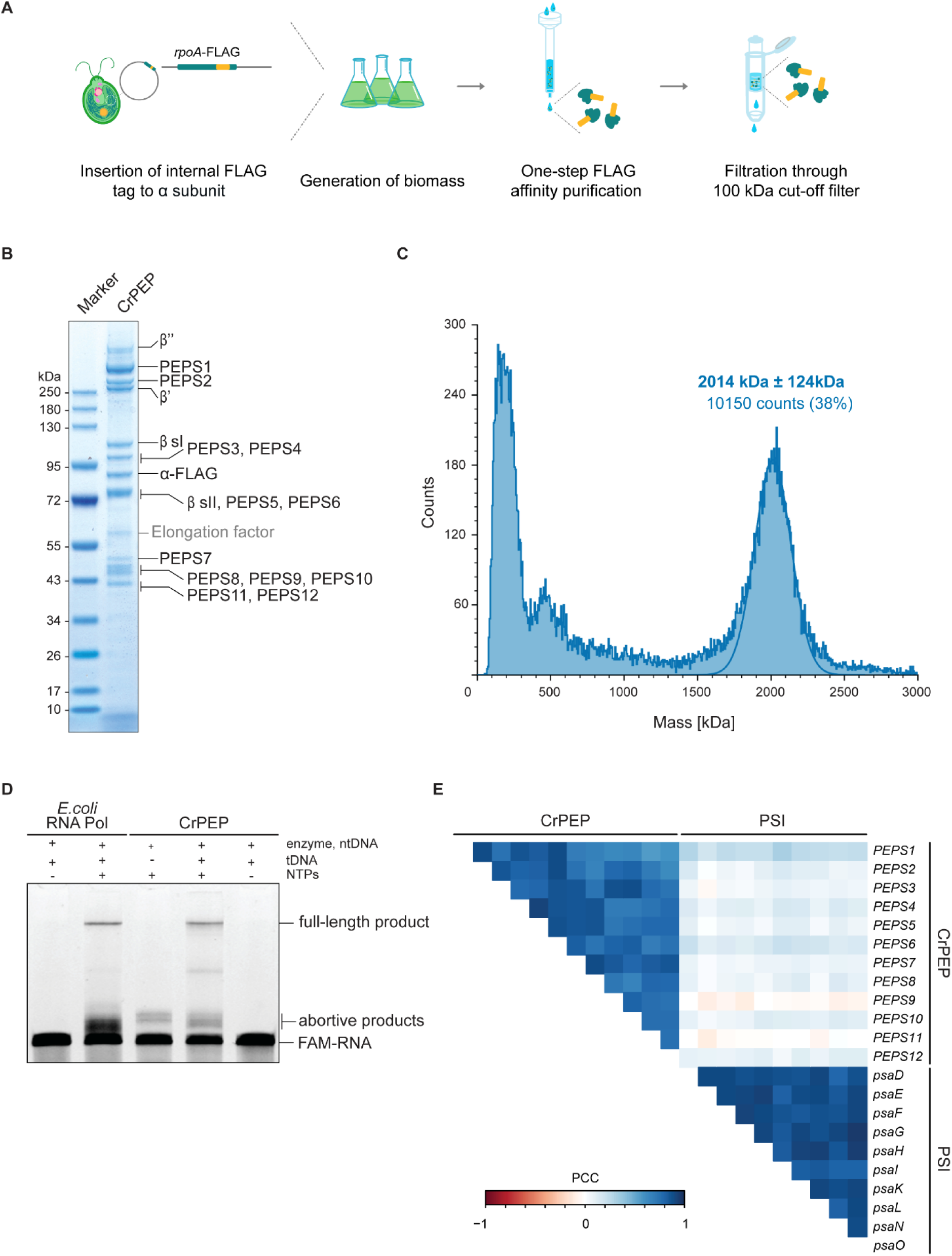
Purification and functional characterization of CrPEP. (**A**) Schematic representation of the strategy used for purification of native CrPEP. (**B**) SDS-PAGE analysis of purified CrPEP. Individual bands were excised and analyzed by LC-MS for the identification of PEP subunits. (**C**) Mass photometry analysis of CrPEP. The solid line shows the best-fit Gaussian distribution. Indicated above are the best-fit molecular mass (kDa), and the width of the peak. (**D**) *In vitro* RNA extension assays. *E. coli* RNA Polymerase (RNA Pol) was included as a positive control. FAM-RNA oligomer and extension products indicated. ntDNA = non-template DNA. tDNA = template DNA. (**E**) Coexpression analysis of nuclear-encoded CrPEP components. Correlation matrix showing Pearson Correlation Coefficients (PCC) for PEPS1-12. Nuclear-encoded photosystem I (PSI) subunits are provided as a reference.

To identify the components of CrPEP, we excised visible bands from the Coomassie-stained gel and subjected them to mass spectrometry analysis (Data S1). Six of the identified subunits correspond to the chloroplast-encoded bacterial-like core of PEP, although their sizes were notably larger than those of their bacterial counterparts. These include two α subunits (encoded by *rpoA*), β sI (encoded by *rpoB1*) and β sII (encoded by *rpoB2*) subunits, representing a Chlamydomonas-specific genetic split of the bacterial β subunit (*rpoB*) (*32*), and β’ (encoded by *rpoC1*) and β’’ (encoded by *rpoC2*) subunits representing a cyanobacteria-inherited genetic split of the bacterial β’ subunit (*13*). Beyond this expanded plastid-encoded core, we identified twelve additional nuclear-encoded subunits with no previously known functions. We refer to these newly identified proteins as PEP Subunits (PEPS), designating them PEPS1 through PEPS12 based on their respective positions when separated by gel electrophoresis (Fig. 1b, 2c)

## Molecular characterization of the CrPEP complex

We reasoned that the newly identified PEPS, as bona fide components of the CrPEP, should localize within the nucleoids of Chlamydomonas, which can be easily visualized with DNA-binding fluorescent dyes such as 4ʹ,6-diamidino-2-phenylindole (DAPI) (*33*, *34*). To test this, we used Chlamydomonas strains expressing C-terminally fluorescently tagged versions of three of the newly identified PEPS proteins (PEPS3, PEPS9, and PEPS12) (*35*) and performed confocal microscopy after fixing the cells and staining them with DAPI. All three proteins were found to form punctate structures that colocalized with nucleoids inside the chloroplast (Fig. S2d). Intriguingly, PEPS12 and PEPS3 were also found inside the pyrenoid, a membrane-less compartment where the carbon-fixing enzyme Rubisco is located. To confirm the correct incorporation of tagged subunits into the CrPEP complex, we performed reciprocal co-immunoprecipitations of PEPS3, PEPS9, and PEPS12, followed by gel electrophoresis and Coomassie staining of the native eluates. Consistently, all CrPEP subunits were detected (Fig. S2a, b). Further, RNA elongation analyses also show that complexes purified from each of these tagged strains show transcriptional activity *in vitro* (Fig. S2c). Some reduction in activity was observed in the preparation from the PEPS12-Venus-3xFLAG strain; while the basis for this is unclear, one possibility is that the presence of this large epitope at the C-terminus of PEPS12 may partially interfere with transcriptional activity.

As additional evidence for the involvement of the newly identified nuclear-encoded PEPS proteins in the CrPEP complex, we analyzed publicly available RNA-Seq datasets from 518 samples across 58 independent experiments (*36*) and calculated Pearson’s correlation coefficients (PCCs) as a measure of transcriptional correlation between genes. We found that the PEPS genes not only form a distinct coexpression cluster with strong positive PCC values, separate from other known clusters such as photosystem I genes (Fig. 1e), but also display coordinated expression with the plastid-encoded subunits of the PEP complex across the diurnal cycle, exhibiting a pattern distinct from other plastid genes (Fig. S3) (*37*). The observed positive correlations suggest coordinated regulation of PEPS genes with PEP subunits, supporting their functional integration within the CrPEP complex.

## Structure of the CrPEP complex

In order to understand how these additional subunits are arranged around and interact with the conserved bacterial core, we subjected purified CrPEP to single-particle cryo-electron microscopy (cryo-EM). Although the presence of a flexible region of the complex posed significant challenges, these were mitigated by focused refinements during processing, resulting in a 3D reconstruction of CrPEP at a global resolution of 2.7 Å (Fig. 2a, Fig. S4, S5, Table S1, Data S2). The high resolution of the map allowed us to assign corresponding EM densities with the proteins most prominently visible on the Coomassie-stained gel, first using first automated sequence tracing (ModelAngelo), and then by manual corrections and re-building of the identified protein chains (Fig. 2b). In this way, we were able to build the majority of the complex, as the cumulative molecular mass of the resolved structure closely matches the ∼ 2 MDa complex previously determined by mass photometry (Table S2).

**Figure 2:**
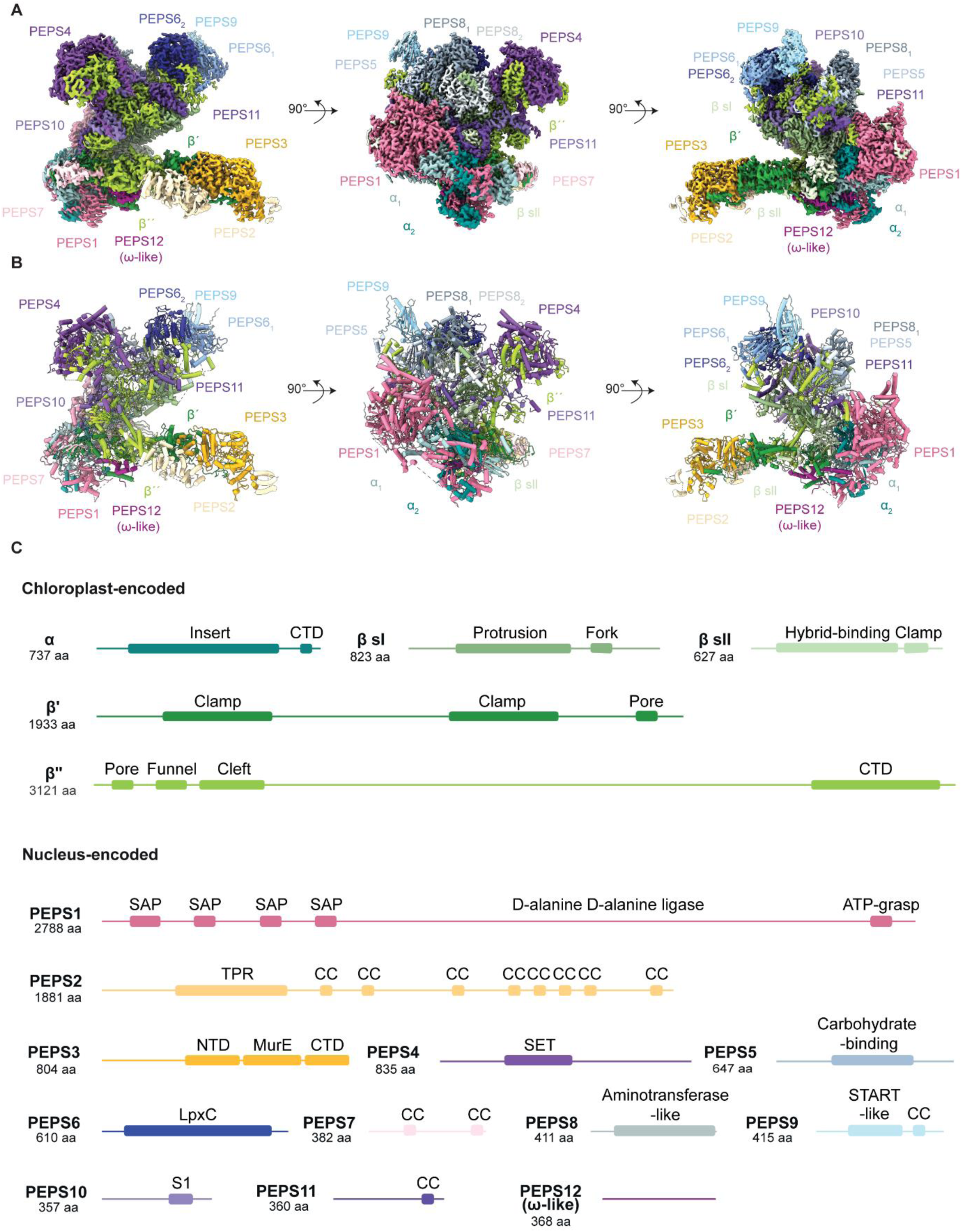
Single-particle cryo-EM structure of CrPEP. **(A)** Segmented cryo-EM reconstruction of CrPEP. Three different views are shown, proteins are coloured as in (C). **(B)** Cartoon representation of the structure, shown in the same orientations and colours. Helices are rendered as solid cylinders.(**C**) Schematic representation of chloroplast-encoded and nuclear-encoded CrPEP subunits with conserved functional domains annotated. NTD = N-terminal domain, CTD = C-terminal domain, CC= Coiled-Coil.

Consistent with observations in other RNA polymerases, the CrPEP core adopts the canonical “crab-claw” architecture: the split β and β′ subunits form the upper and lower pincers of the active-site cleft, while the α-subunit dimer stabilizes the assembly at the rear. In addition to this conserved core, CrPEP contains three peripheral modules that are largely contributed by PEPS subunits: a crown-like apical module, a posterior “backpack” module, and a projecting “tongue” module. Multiple PEPS proteins (PEPS4, PEPS5, PEPS6, PEPS8, PEPS9, PEPS10, and PEPS11) associate with the expanded Sequence Insertion 3 domain (hereafter referred to as SI3) located in the plastid-encoded β’’ subunit (residues 448–3028), forming the crown-like module along the upper jaw of the polymerase. At the rear of the complex, the ∼280 kDa PEPS1 together with the smaller PEPS7 forms the backpack module that overlays the α-subunit dimer. Finally, PEPS2 and PEPS3 reside in the lower jaw and extend outward to form the tongue-like projection together with a large part of the β′ and β’’ subunit extension (Fig. 2b)

## The crown-like region – stabilization of SI3 extension by enzymatic folds

As in the cyanobacterial RNA polymerase, the CrSI3 in the β" subunit is located between the two helices of the trigger loop, a highly conserved region that undergoes an essential conformational change with the addition of each nucleotide in the nascent RNA chain (Fig. 3a) (*38*). This domain retains a structurally similar organization to that of the cyanobacterial complex, which is characterized by four distinct segments: the SI3 tail, the SI3 fin, the SI3 body, and the SI3 head (Fig. 3b). However, the fin and head domains of CrSI3 are massively expanded, providing binding sites for several of the newly identified PEPS proteins (Fig. 3c). These PEPSs often exhibit folds reminiscent of metabolic enzymes, as identified via Foldseek and DALI searches. For instance, PEPS4, contains a SET-binding/Rubisco-LSMT substrate binding domain I (*39*) and interacts with the CrSI3 fin domain through a pincer-like extension comprised of a beta-sheet hairpin on one side and a helical bundle on the other. Meanwhile, the expanded head domain of CrSI3 provides an additional binding interface for further PEPS subunits containing enzymatic domains. These include PEPS5, which adopts a glucose-6-phosphate epimerase fold (*40*); PEPS6, a homodimer harboring a UDP-3-O-acyl-N-acetylglucosamine deacetylase domain—(characteristic of LpxC deacetylases involved in lipid A biosynthesis in Gram-negative bacteria) (*41*); and PEPS8, another homodimer positioned centrally within the upper jaw region and structurally similar to branched-chain amino acid transaminases (*42*). PEPS6 also serves as an anchor for PEPS9, a START-domain–containing protein typically associated with lipid-related enzymes such as polyketide cyclases and COQ10 (*43*). Despite their enzyme-like folds, some of these PEPS subunits lack conservation of canonical active-site features or display structural occlusion of regions corresponding to putative substrate-binding pockets. (Fig. S6). For example, in PEPS5, essential catalytic residues are missing. In PEPS4 and PEPS9, substitutions introduce steric clashes that would obstruct substrate binding. In PEPS8, the putative substrate entry channel is blocked by neighbouring partners in the complex. For PEPS6, which resembles a bacterial cell wall–biosynthetic enzyme (*44*), the predicted substrates are not present in the chloroplast.

**Figure 3:**
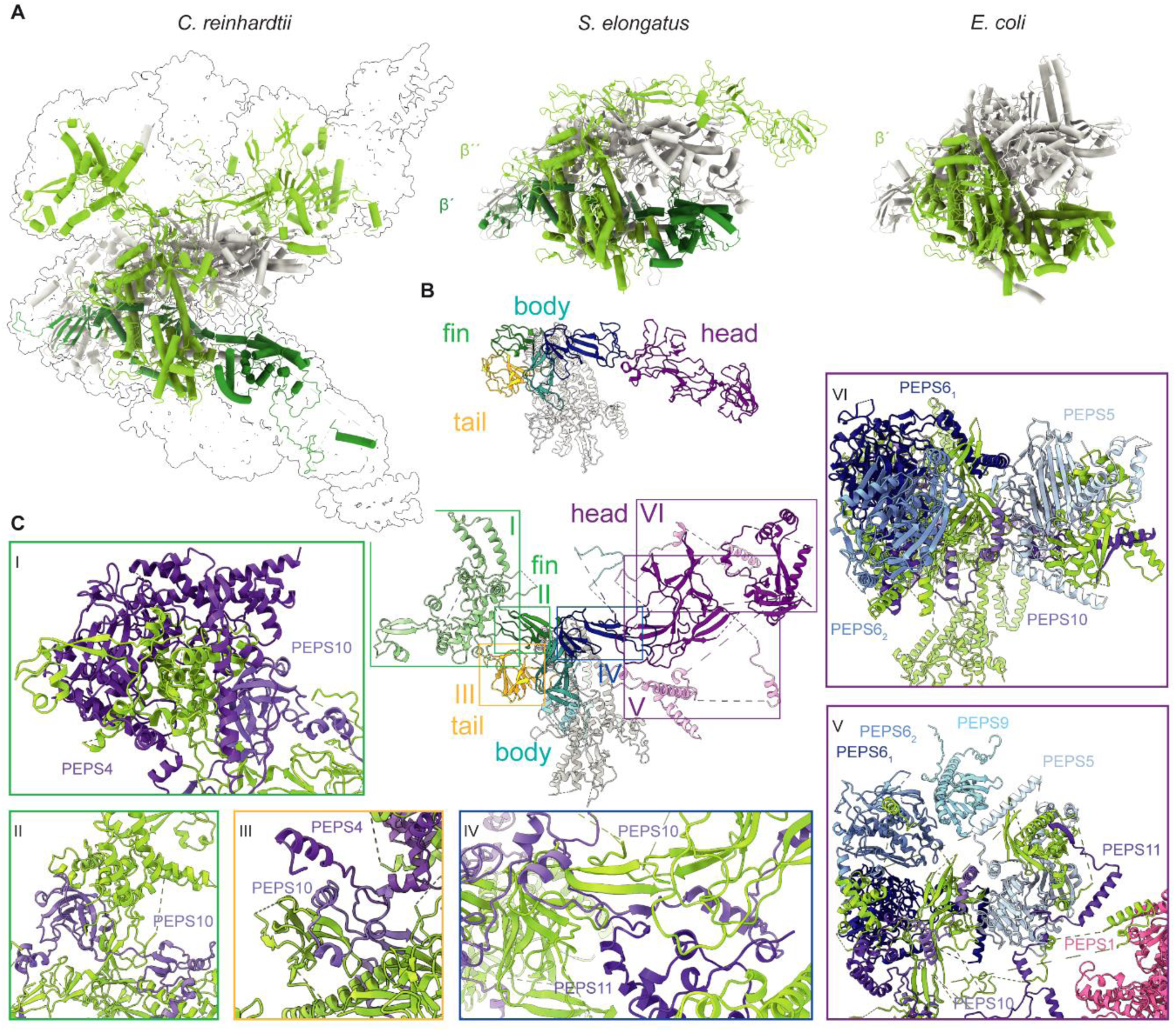
The plastid-encoded core of CrPEP is structurally related to that of the *S. elongatus* RNA polymerase. **(A)** Comparison of conserved β′and β′′ subunits in CrPEP with corresponding subunits in *S. elongatus* (PDB: 8syi) and *E. coli* RNA polymerases (PDB: 4yln). (**B**) Structural organisation of *S. elongatus* β′′ SI3 domain, including SI3 tail (yellow), SI3 fin (green), SI3 body (turquois and blue), and SI3 head (violet) with comparison to the corresponding regions in the CrSI3 domain. (**C**) Close-up views showing the intermolecular interactions between the PEPS proteins and the protrusions formed by the fin, the tail, the body, and the head of CrSI3 (I) PEPS4 and PEPS10 binding to the fin region. (II) PEPS10 binding to the fin region. (III) PEPS4 and PEPS10 binding to the tail region. (IV) PEPS10 and PEPS11 binding to the upper body region. (V) Multiple PEPS proteins (PEPS1, PEPS5, both PEPS6 monomers, PEPS9, PEPS10) interacting with the lower head region. (VI) PEPS5, both PEPS6 monomers, and PEPS10 binding to the upper head region.

We also identified two elongated, largely unstructured subunits, PEPS10 and PEPS11, which make extensive contacts with the CrPEP upper jaw and might therefore support its structural integrity (Fig. 3c). PEPS10, which contains an S1-domain motif, spans multiple domains of the CrSI3. Its N-terminal, β-sheet–rich region interacts with the CrSI3 tail, while its S1 domain wraps around the conserved section of the CrSI3 fin, forming a cuff-like structure, before extending further toward the midpoint of the CrSI3 head arch. PEPS11 follows a more elaborate trajectory: contacting the CrSI3 body via its N-terminal region, arching downward near the fin, and ascending again to contribute an additional β-strand to the CrSI3 head. In doing so, this protein forms multiple interactions with the PEP core subunits and various other PEPS, hence is likely to contribute to the overall stability of the complex.

## The tongue

The lower jaw of the CrPEP complex accommodates the binding sites for three additional PEP subunits, PEPS2, PEPS3 and PEPS7, as well as a substantial portion of the conserved core protein β’ (Fig. 4a). Interestingly, β’ occupies two distinct regions in the complex: one centrally located and conserved in the available land plant PEP structures, and another in the lower jaw, forming a Chlamydomonas-specific N-terminal extension.

**Figure 4:**
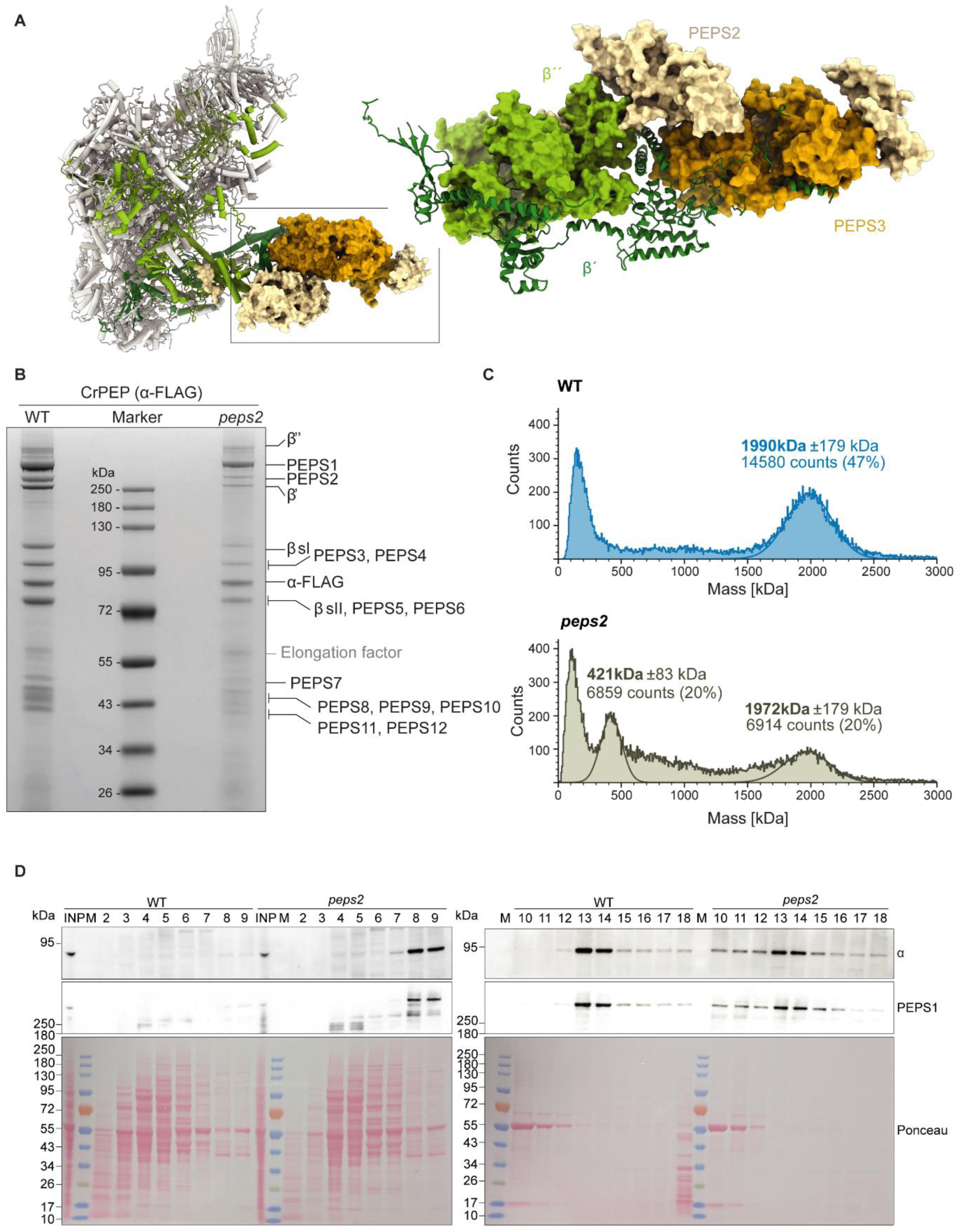
PEPS2 contributes to CrPEP complex assembly and stability. (**A**) Cartoon representation of the structure of CrPEP with PEPS2 and PEPS3 shown as surface representation. The colors of β′, β′′, PEPS2 and PEPS3 are the same as in Fig. 2c, whereas the remaining PEP subunits are depicted in grey. Inset shows a close-up view of the interaction of β′ with PEPS3 and PEPS2. The lower part of β′′ is shown as surface representation. (**B**) SDS-PAGE analysis of CrPEP purified from wild-type and *peps2* mutant cells. (**C**) Mass photometry analysis of CrPEP eluates from wild type and *peps2* mutant backgrounds. (**D**) Immunoblots using PEPS1 and α-subunit antibodies following sucrose gradient (15-45%) fractionation of whole-cell lysates from wild type and *peps2* mutant. Fractions collected from top (Fraction 1) to bottom (Fraction 18). INP = Input.

In the conserved central region, the C-terminal end of β′ forms an extended β-sheet with PEPS7, while the linker connecting these regions is stabilized by a helical bundle that inserts between the central core and the lower jaw. At the distal tip of the lower jaw, β′ makes direct and extensive contacts with PEPS3— the best conserved PEPS subunit conserved at both structural and sequence level between *Chlamydomonas* and land plant PEP complexes (Fig. 4a and 6b) (*9*). PEPS3 adopts a fold similar to MurE, a UDP-N-acetylmuramoyl-L-alanyl-D-glutamate–2,6-diaminopimelate ligase involved in bacterial cell wall biosynthesis (*45*). However, PEPS3 is unlikely to retain enzymatic activity once assembled into the complex, as its substrate-binding site is occupied by the N-terminal region of β′, preventing substrate access. Instead, PEPS3 appears to function primarily as a structural element within CrPEP, forming an extensive interface with β′ that spans both the protein surface and the occluded pocket region. This mode of engagement may be particularly relevant in green algal lineages where β′ is encoded as two separate polypeptides (β′ and β′′), potentially increasing the requirement for inter-subunit stabilization. In addition, the placement of PEPS3 at the lower jaw may contribute to maintenance of an open-jaw conformation, a feature observed across the available PEP structures from land plants (*8–11*), and one that may relate to plastid-specific properties of transcription. Additional stabilization of the lower jaw is provided by PEPS2, a large tetratricopeptide repeat (TPR)–containing subunit. Pronounced conformational heterogeneity in this region limited local resolution, precluding complete modeling of PEPS2. Nevertheless, the resolved densities indicate that PEPS2 is organized into two distinct segments: one segment anchors laterally to β′, β′′, and PEPS3 through a spiral fold typical of TPR domains (*46*), while the other interacts predominantly with the front face of PEPS3, without making additional contacts with other PEPS subunits. This spatial isolation, combined with its modular TPR structure, suggests that PEPS2 may serve a specialized, potentially regulatory role in complex assembly or transcriptional activity.

**Figure 5:**
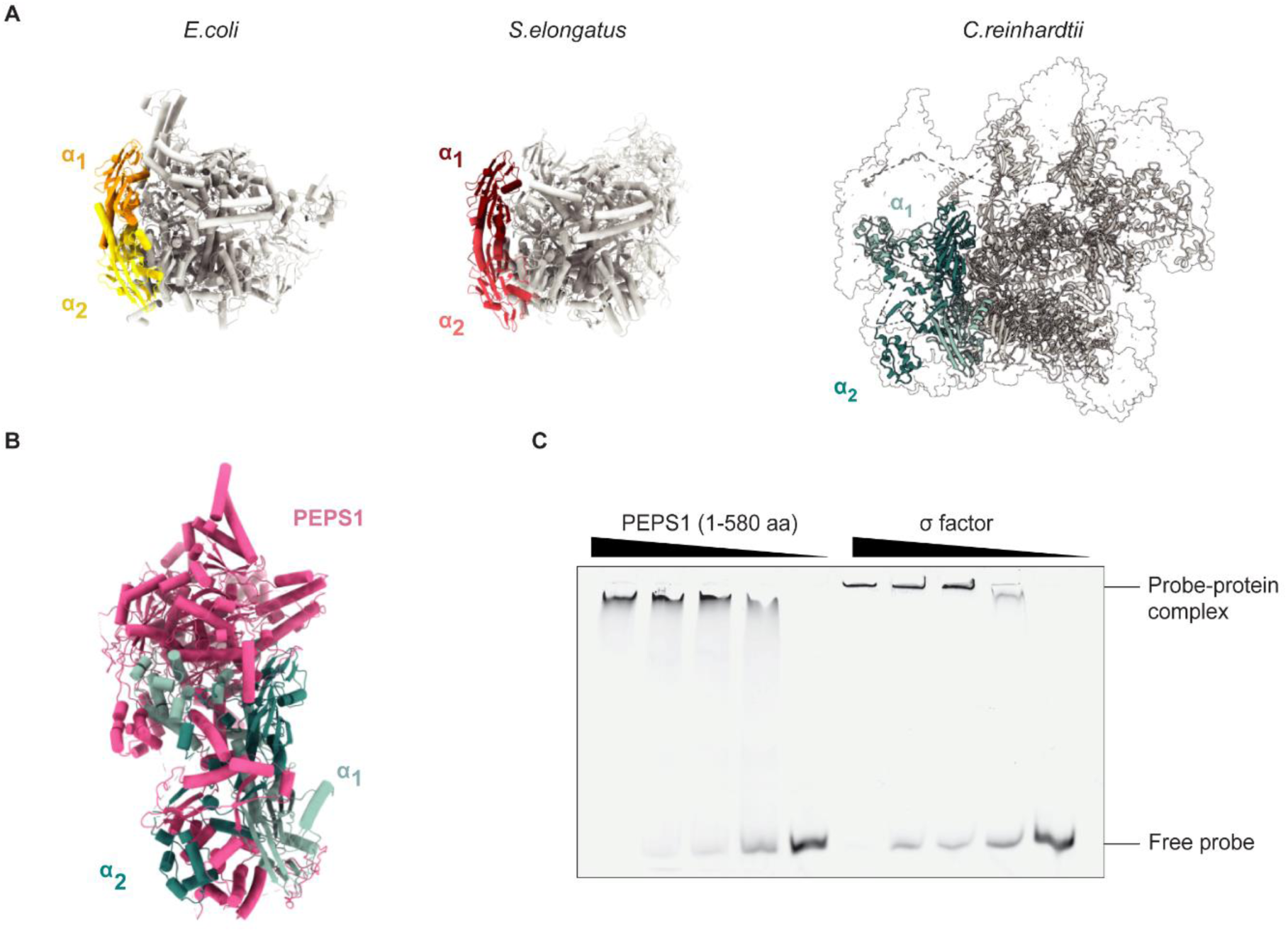
Interactions of PEPS1 with the α-subunits in the backpack region of CrPEP. (**A**) Cartoon representation of α_1_ and α_2_ subunits of the *E. coli* (PDB: 4yln) and *S. elongatus* (PDB: 8syi) RNA polymerases in comparison with those of CrPEP. The α subunits are shown in two different shades of yellow, red, and turquoise, respectively, while the other subunits of the complexes are shown in grey. (**B**) Cartoon model illustrating the stabilization of CrPEP α_1_ and α_2_ through their interaction with PEPS1. (**C**) Electrophoretic mobility shift assay (EMSA) using two-fold serial dilutions of a truncated form of PEPS1 containing the four SAP domains shown in Fig. S10 (residues 1–580; 2.5–0 µM) and the Cr σ factor (10-0 µM) with a 147 bp DNA probe corresponding to the *C. reinhardtii* plastid 16S rRNA promoter region.

**Figure 6:**
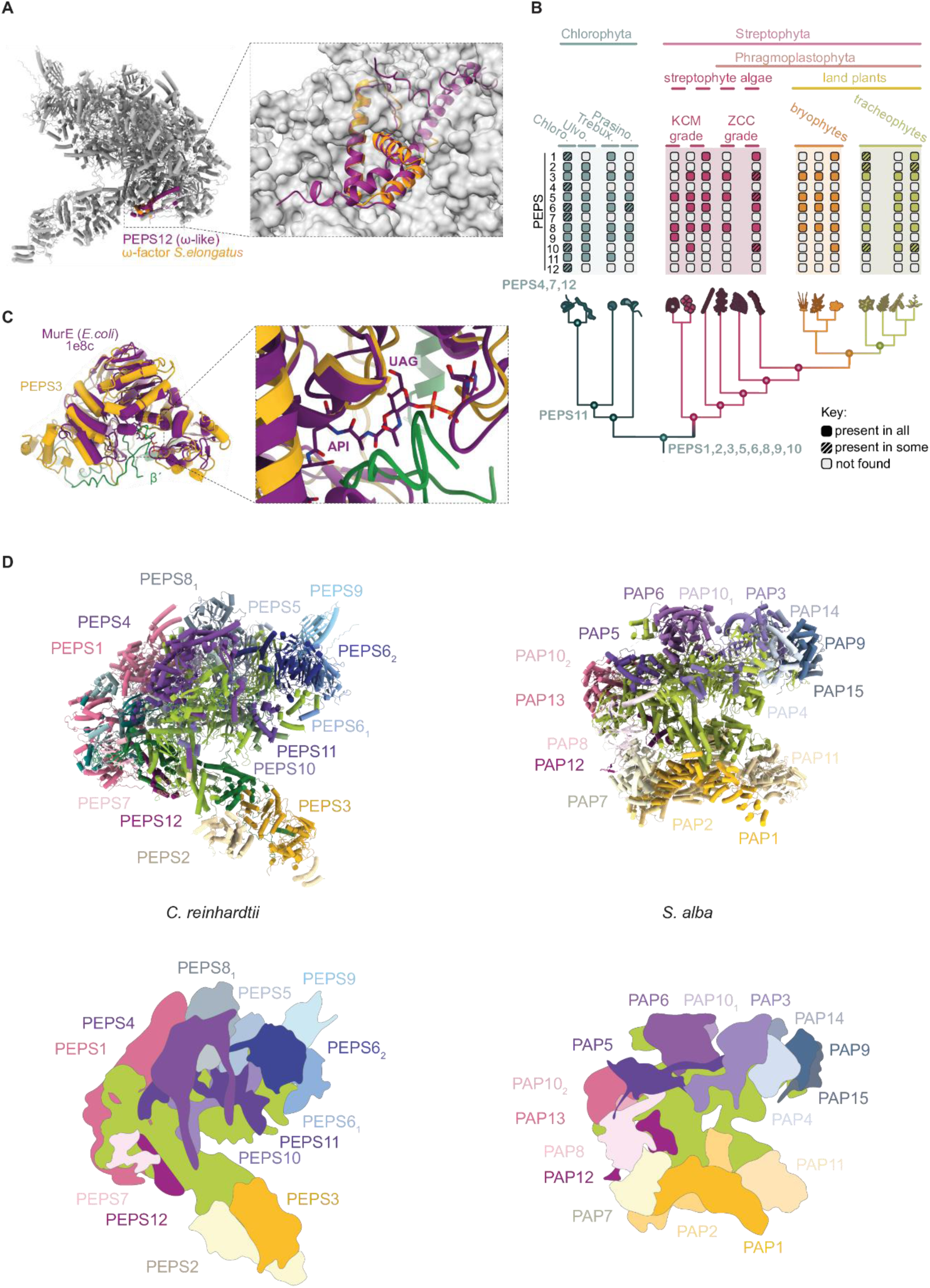
Comparison of CrPEP to land plant PEP and cyanobacterial RNAP complexes. (**A**) Superposition of PEPS12 (violet) with the ω factor from *S. elongatus* (PDB: 8urw, orange) in the context of the CrPEP shown as grey cartoon representation (left), and a close-up view with the remaining CrPEP subunits shown as surface representation (right). (**B**) Evolutionary tree showing the phylogenetic distribution of PEPS proteins across Viridiplantae. (**C**) Superposition of PEPS3 (orange) and the β′ loop of CrPEP (green) with *E. coli* MurE (violet, PDB: 1e8c). The close-up on the right shows that the β′ loop occupies the substrate-binding pocket of MurE. MurE substrates annotated as UAG (UDP-N-acetylmuramoyl-L-alanine-D-glutamate), and API (2,6-diaminopimelate). (**D**) Comparison of the chloroplast RNA polymerase structures from *C. reinhardtii* and *S. alba* (PDB: 8r6s), shown as cartoon representations and schematic drawings.

To explore this hypothesis, we searched a genome-wide insertional mutant library in *C. reinhardtii* (*47*) and identified a mutant harbouring an insertion within an intron of the nuclear PEPS2 gene, resulting in an approximately 80% reduction in transcript abundance (Fig. S7a).

In order to investigate the effect of this mutation on CrPEP, a FLAG affinity tag was introduced into the endogenous *rpoA* gene of this mutant via chloroplast transformation as before (Fig. S1). CrPEP purifications from this background showed an approximately eight-fold reduction in *in vitro* transcriptional activity compared to wild type, accompanied by changes in subunit stoichiometry, particularly affecting components near PEPS2 (Fig. 4b, c, Fig. S7, Fig. S8, Data S3, Data S4). Mass photometry and sucrose gradient fractionation identified a ∼500 kDa subcomplex containing the α subunit and PEPS1 (Fig. 4d., Fig. S7, Fig. S8, Fig. S9, Data S5). Because α-subunit dimerization is the first step in bacterial RNA polymerase assembly (*48*), the accumulation of this intermediate in the absence of PEPS2 suggests that PEPS subunits are incorporated together with the core during assembly and that, in the absence of PEPS2, complex assembly stalls at an intermediate stage rather than forming the fully assembled PEP. Interestingly, despite the pronounced defect in *in vitro* enzymatic activity, steady-state levels of chloroplast rRNAs and mRNAs, including those encoding photosynthetic and ribosomal proteins, remained largely unchanged, and no growth defects were observed in the *peps2* mutant (Fig. S7d,e). One possibility is that residual CrPEP activity in the *peps2* knockdown is sufficient to sustain near–wild-type transcript levels, for example if CrPEP is present in excess under the tested conditions. Alternatively, transcript abundance may be maintained by buffering mechanisms operating downstream of transcription, such as increased RNA stability (*49*).

## The backpack region

At the rear of the CrPEP complex, PEPS1 associates with the conserved homodimeric α-subunit core. As in all eubacterial RNA polymerases, the α-subunit comprises two independently folded domains: the N-terminal domain (NTD) and the C-terminal domain (CTD), connected by a flexible linker (*50*). However, in CrPEP, this linker is significantly extended and features α-helical insertions (Fig. 5a). PEPS1, a 280 kDa protein containing four SAP domains at its N-terminus, wraps around the α-dimer in a configuration reminiscent of a backpack (Fig. 5b, Fig. S10). While the SAP domains are not resolved in the structure, presumably due to their conformational flexibility, the remaining portion of PEPS1 makes extensive contact with both the α-CTD and its extended linker via a complementary electrostatic surface. These contacts likely contribute to the stabilization of the CrPEP α-homodimer, but they also appear to restrict the mobility of the CTD’s helical bundle, a region known in bacterial RNA polymerases to mediate recognition of upstream promoter elements. Given that SAP domains are well-characterized nucleic acid-binding modules, typically recognizing DNA in a non-sequence-specific manner (*51–53*), we hypothesize that the flexible SAP domains of PEPS1 may functionally substitute for the constrained α-CTD in promoter recognition (*54*). Consistent with this, a recombinant PEPS1 fragment containing all four SAP domains exhibited DNA-binding activity in electrophoretic mobility shift assays using a probe corresponding to the 16S rRNA promoter region, with an affinity comparable to that of *C. reinhardtii* σ (Fig. 5c, Fig. S11), demonstrating that these domains alone are sufficient to engage DNA and may contribute to CrPEP promoter recruitment.

## Discussion

We have shown that *C. reinhardtii*, a unicellular green alga that diverged from land plants over a billion years ago (*29*), possesses a PEP complex that is roughly twice the size of its land-plant counterparts, making it the largest RNA polymerase structurally characterized to date. While most of the newly identified subunits in CrPEP were likely present in the last common ancestor of Chloroplastida, as supported by their broad distribution across streptophytes in our phylogenetic analyses (Fig. 6b), their evolutionary relationship to land-plant PAPs cannot generally be inferred from sequence conservation alone.

PEPS3, which contains a MurE catalytic domain, is the only newly identified subunit with clear sequence-level conservation to a land-plant PAP (PAP11/MurE) (Table S3), suggesting it may represent one of the earliest recruited proteins in the evolution of this complex (Fig. 6b–c). PEPS2 appears orthologous to PAP1/pTAC3 (Data S6), although this relationship is far less evident at the sequence level; its structural position within the complex nonetheless supports this assignment and points to substantial divergence over evolutionary time (Fig. 6d). Particularly intriguing is the case of PEPS12, which displays structural similarity to the bacterial ω subunit and to land-plant PAP12, yet lacks detectable sequence conservation. This observation points either to extensive sequence divergence or to the independent recruitment of a non-homologous protein likely fulfilling a similar structural role within the complex (Fig. 6a–b).

The remaining newly identified subunits have no detectable counterparts among known land-plant PAPs, and several appear to have been secondarily lost in a lineage-dependent manner (Fig. 6b). Notably, even when a protein is highly conserved at the sequence level, its incorporation into the complex is not guaranteed, as exemplified by PEPS8, whose homologs have not been detected in any land-plant PEP structure to date. These observations argue that additional constraints—beyond sequence conservation—determine which accessory proteins are integrated into PEP across photosynthetic eukaryotes. Consistent with this idea, several non-homologous PEPS occupy positions around the polymerase core that correspond to sites populated by PAPs in land plants. This is particularly evident in the upper jaw, where the β′′-SI3 arch is stabilized by cooperative binding of peripheral factors in both systems, and the backpack region, where dedicated subunits engage the α-CTDs despite sharing no sequence homology with their plant counterparts.

The presence of a highly elaborated PEP complex in a unicellular alga— despite the fact that chloroplast gene expression is largely regulated post-transcriptionally and transcript levels exhibit limited responsiveness to environmental cues (*55*)—raises a fundamental question: why did such complexity arise? A compelling explanation is offered by the theory of constructive neutral evolution (CNE), whereby complexity emerges not through adaptive selection for novel regulatory functions, but through the evolutionary entrenchment of initially non-adaptive interactions (*56–58*). In this framework, fortuitous, low-affinity associations between the polymerase core and pre-existing cellular proteins arise neutrally and persist. Subsequent mutations in the core enzyme—such as insertions, expansions, or conformational destabilization—are then tolerated only in the presence of these interacting partners, converting dispensable associations into structural necessities. Once this dependency is established, accessory subunits become irreversibly integrated, locking the system into a more elaborate, but not necessarily more regulated state.

In the population-genetic context of organellar genomes—marked by high copy number and reduced effective population size—this model provides a unifying explanation for several otherwise paradoxical features of PEP. Many accessory subunits in both Chlamydomonas and land plants retain recognizable enzyme-like folds yet display loss of catalytic residues or occlusion of active sites upon incorporation into the complex, consistent with structural co-option rather than de novo evolution for transcription. Moreover, distinct proteins can fulfill equivalent architectural roles in different lineages, explaining why non-homologous factors stabilize analogous regions of the polymerase core.

Resolving whether the PEPS subunits in Chlamydomonas and the PAP subunits in land plants act solely as structural stabilizers or have acquired additional regulatory functions is a critical step toward testing the constructive neutral evolution hypothesis for the emergence of this essential protein complex. Chlamydomonas, with its unique blend of dual genetic manipulability of nuclear and plastid genomes and biochemical accessibility (*30*, *59*, *60*), provides an ideal platform for this effort—enabling not only mechanistic dissection but also synthetic reconstruction of ancestral architectures and the design of engineered complexes, thereby linking evolutionary biology with applied synthetic biology.

## Resource Availability

### Lead contact

- Requests for further information and resources should be directed to and will be fulfilled by the lead contact, Silvia Ramundo

### Materials availability

- All unique/stable reagents generated in this study are available from the lead contact with a completed material transfer agreement

### Data and code availability

- Cryo-EM reconstructions of the CrPEP complex have been deposited in the Electron Microscopy Data Bank and RCSB data bank and are publicly available under PDB accession number 28RF, and EMDB accession numbers: 56766, 56920, 56921 for the consensus map, tongue region, and core region, respectively.
- The mass spectrometry proteomics data have been deposited to the ProteomeXchange Consortium via the PRIDE partner repository with the dataset identifier PXD073490.
- Raw confocal microscopy images for PEPS3, PEPS9, and PEPS12-Venus-3xFLAG tagged lines are available through Figshare (https://figshare.com/s/a761003830830efa0663).
- This paper analyzes existing, publicly available data, accessible at https://doi.org/10.1093/plcell/koab042 for general coexpression analysis (Fig. 1e) and https://doi.org/10.1073/pnas.1815238116 for diurnal coexpression analysis (Fig S3)
- All other data is available in the manuscript or associated supplementary materials.
- Any additional information required to reanalyze the data reported in this paper is available from the lead contact upon request.

## Acknowledgments

We thank the Vienna Biocenter community for stimulating scientific discussions, the Stanford Mass Spectrometry Facility and the Vienna Biocenter Facilities (Biooptics, Mass Spectrometry, Protein Technologies, and Biophysics) for providing essential technical support and expertise, and Dorotea Fracchiolla, PhD (Art&Science) for professional assistance in finalizing the figures. We thank the Cryo-EM facilities at Philipps-University Marburg and Helmholtz Munich for data collection. We thank Martin Jonikas for providing early access to PEPS3, PEPS9, and PEPS12 tagged lines. We thank Renato Arnese, Alejandro Burga, Tim Clausen, Liam Dolan, Matthias Kopf-Vorländer, Steven McKnight, Magnus Nordborg, and Peter Walter for providing constructive feedback on this manuscript.

S.R. further acknowledges Martin Jonikas and Peter Walter for funding the initial mass spectrometry experiment that led to the discovery of the CrPEP complex.

## Funding

Austrian Academy of Sciences (SR)

Austrian Science Fund AST1678424, F7916-B (SR)

Federation of European Biochemical Societies (Excellence Award) (SR)

LOEWE RobuCop (JMS)

LOEWE Tree-M (JMS)

Deutsche Forschungsgemeinschaft (DFG, German Research Foundation) under Germany’s Excellence Strategy – EXC 3048 – Project number 533620160: Microbes-for-Climate (M4C) Cluster of Excellence, Synmikro, Marburg, Germany (JMS)

Deutsche Forschungsgemeinschaft (DFG, German Research Foundation) Project number 515101361 (JdV,SdV)

MAdLand (DFG priority programme 2237) 527846412 (JMS)

MAdLand (DFG priority programme 2237) 440231723 (JdV)

## Author contributions

S.R. conceived the project, supervised all research activities except structural and phylogenetic analyses, developed the PEP purification protocol, purified and characterized the initial complex, identified its subunits in collaboration with the Stanford Mass Spectrometry Facility, provided resources, and coordinated collaborations. J.M.S. supervised structural work and provided resources. J.D.V. coordinated phylogenetic analyses and provided resources. R.C. carried out coexpression analyses. R.C., S.R., J.H., and L.W. analyzed mass spectrometry data. C.B. and C.L.W. generated chloroplast- and nuclear-tagged strains, respectively. R.C., C.B., J.H., and P.F. purified PEP complexes for biochemical analyses. P.F. developed the transcriptional elongation assay. J.K. generated and purified recombinant Cr σ factor. R.C., C.B., P.V., and C.-L.W. characterized the *peps2* mutant. R.C. designed and purified PEPS1Trunc protein and performed EMSAs. R.C. and A.K. ensured public data deposition. A.K. and A.I. purified the PEP complex for cryo-EM analysis, optimized crosslinking and grid preparation protocols. A.K. collected and processed cryo-EM data and performed structural validation. S.K.S. built the atomic model, performed structural interpretation and analysis, assisted with grid preparation and contributed to respective figure preparation. S.D.V. and I.I. conducted and interpreted phylogenetic analyses and prepared the respective figure. C.G. contributed to structural analysis. S.B. collected cryo-EM data. S.R., J.M.S., R.C., A.K., and S.K.S. wrote the original manuscript draft and assembled the original figures with contributions from all authors. S.R. and R.C. coordinated the final editing of the manuscript and figures. All authors reviewed and approved the final manuscript.

## Supplementary Information

**Data S1. (dataS1_AlphaFLAGIP_BandIdentification_MSResults.xslx)**

Mass Spectrometry results for band identification from α-FLAG IP CrPEP purification

**Data S2. (dataS2_Validation_Report_PEP.pdf)**

cryo-EM validation report for CrPEP

**Data S3. (dataS3_CellLysate_*peps2*vsWT_MSResults.xslx)**

Mass Spectrometry results comparing abundance of CrPEP subunits in whole-cell lysate from *peps2* mutant and wild-type backgrounds

**Data S4. (dataS4_AlphaFLAGIP_*peps2*vsWT_Eluate_MSResults.xslx)**

Mass Spectrometry results comparing abundance of CrPEP subunits in α-FLAG IP eluate from *peps2* mutant and wild-type backgrounds

**Data S5. (dataS5_SucroseGradient_*peps2*vsWT_MSResults.xslx)**

Mass Spectrometry results comparing abundance of CrPEP subunits in selected sucrose gradient fractions from whole-cell lysate derived from *peps2* mutant and wild-type backgrounds

**Data S6. (dataS6_PEPSTrees.zip)**

Maximum likelihood phylogenies of the PEPS proteins and their homologs across the green lineage

## Supplementary Figures

**Figure S1.**
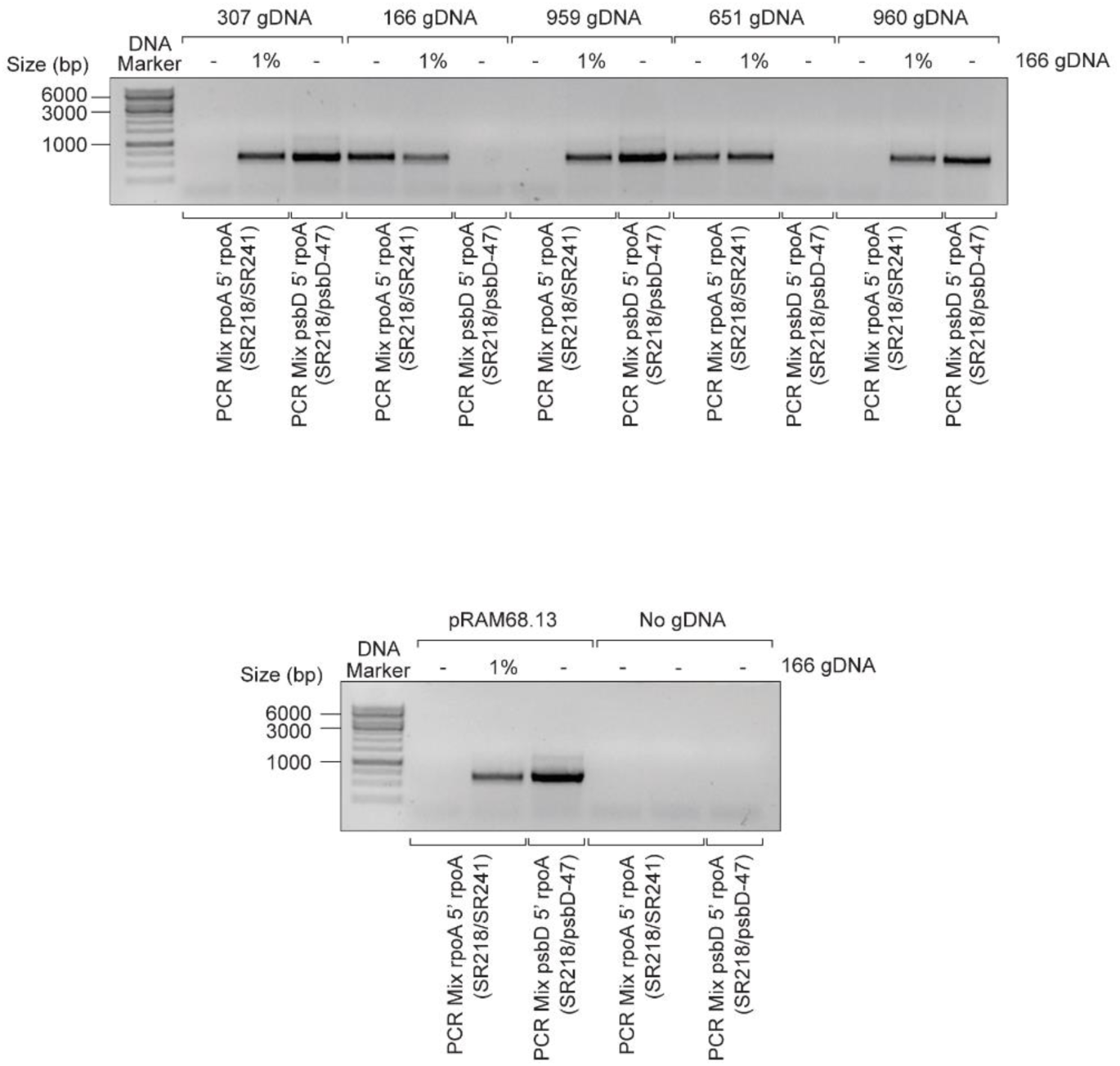
PCR analysis used to verify the homoplasmicity of chloroplast transformant strains. Genomic DNA (gDNA) from chloroplast transformant strains (307, 959, 960), the wild-type (166) and *peps2* mutant (651) background strains, and a plasmid (pRAM68.13) containing the *psbD5*’*rpoA* chimeric gene were used as templates. Primer mixtures specific for *psbD5’rpoA* (psbD-47/SR218) and *rpoA*5’*rpoA* (SR218/SR241) were used for amplification. 1% of wild-type DNA was added as a control for heteroplasmicity corresponding to less than one gene copy per chloroplast. 5’ = 5’ UTR, marker = GeneRuler 1kb DNA Ladder (ThermoScientific).

**Figure S2.**
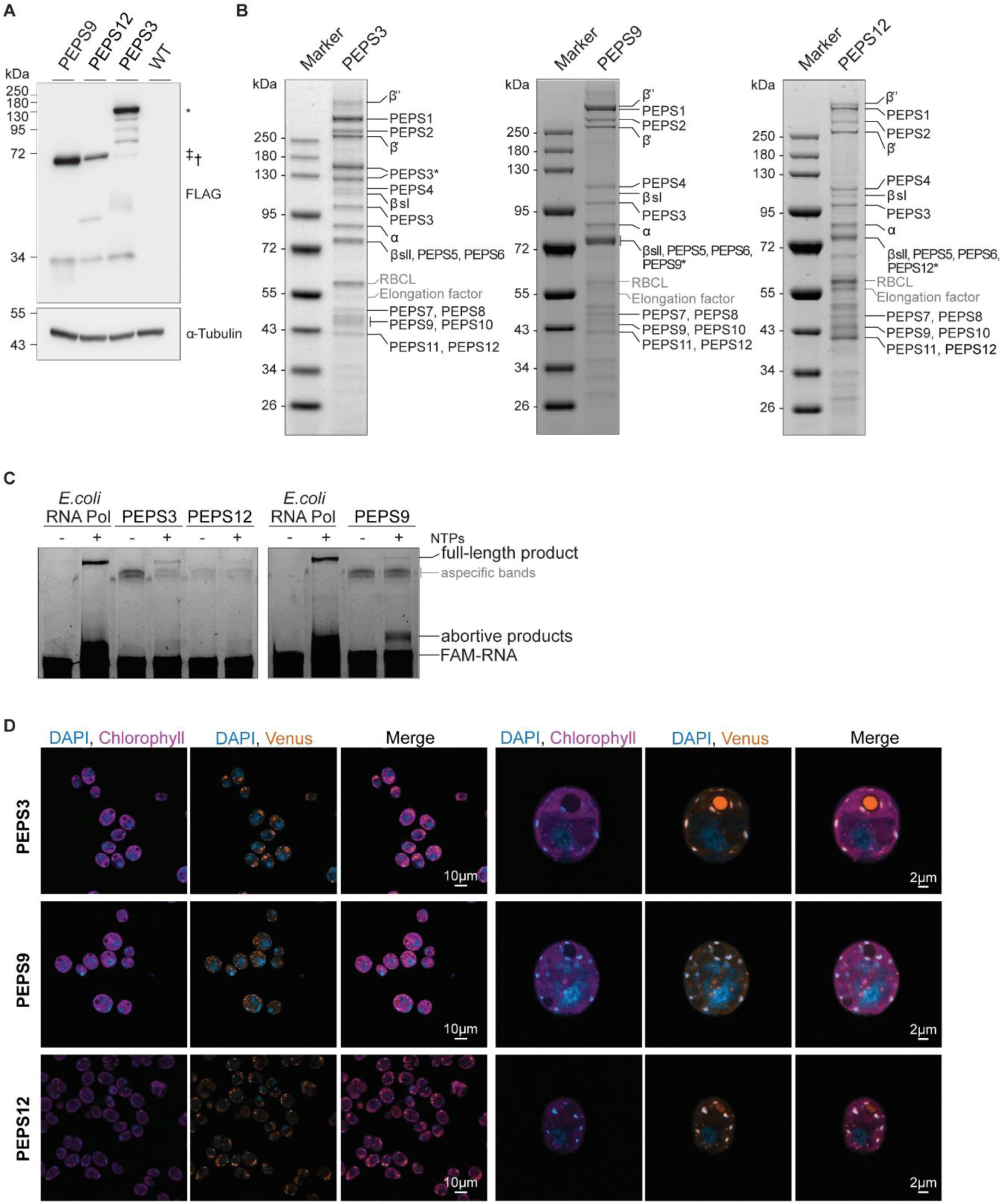
CrPEP purified from strains expressing differing tagged PEPS proteins showing similar composition and activity. (**A**) Immunoblot analysis of whole cell lysates from strains expressing Venus-3xFLAG-tagged PEPS3, PEPS9, and PEPS12. Symbols denote tagged full-length proteins (*, PEPS3; †, PEPS9; ‡, PEPS12); smaller bands represent degradation products. (**B**) SDS-PAGE analysis of CrPEP purified from strains expressing PEPS-Venus-3xFLAG-tagged proteins. Asterisks indicate position of tagged protein. (**C**) *In vitro* RNA extension assays using CrPEP purified from strains expressing PEPS-Venus-3xFLAG-tagged proteins compared to *E. coli* RNAP control. FAM-RNA oligomer and extension products are indicated. (**D**) Broad-field and close-up confocal microscopy images of *C. reinhardtii* cells expressing Venus-3xFLAG-tagged PEPS proteins (PEPS3, PEPS9, and PEPS12), showing subcellular localization of CrPEP. Nuclei and chloroplast nucleoids were stained with DAPI.

**Figure S3.**
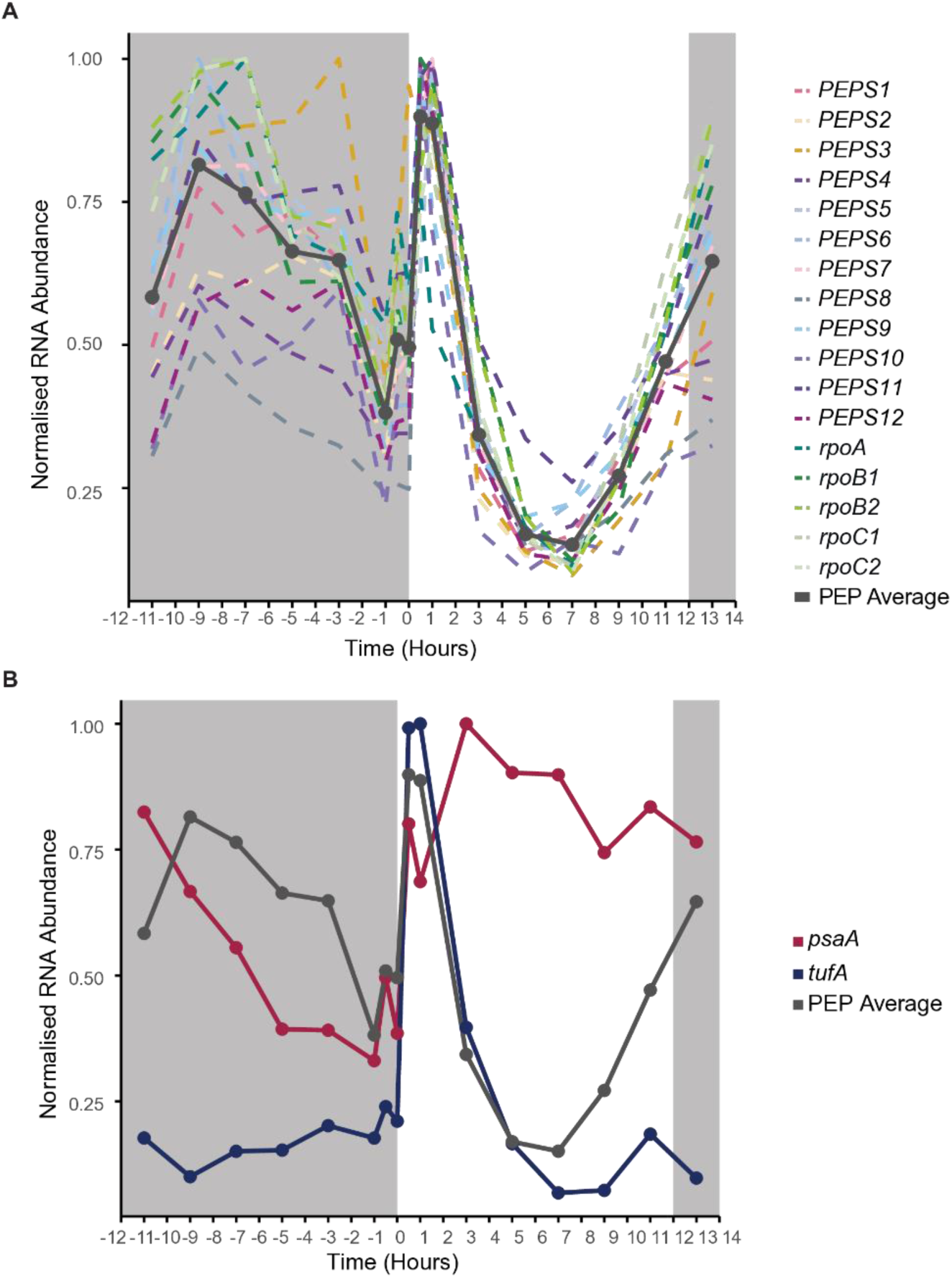
Diurnal expression patterns of Chlamydomonas genes encoding components of the CrPEP complex. (**A**) Coexpression profiles of plastid and nuclear-encoded CrPEP components over a diurnal cycle in Chlamydomonas. Shaded areas represent the dark period. (**B**) Average diurnal coexpression profile of CrPEP components (grey) compared to representative plastid genes: *psaA* (red) and *tufA* (blue).

**Figure S4.**
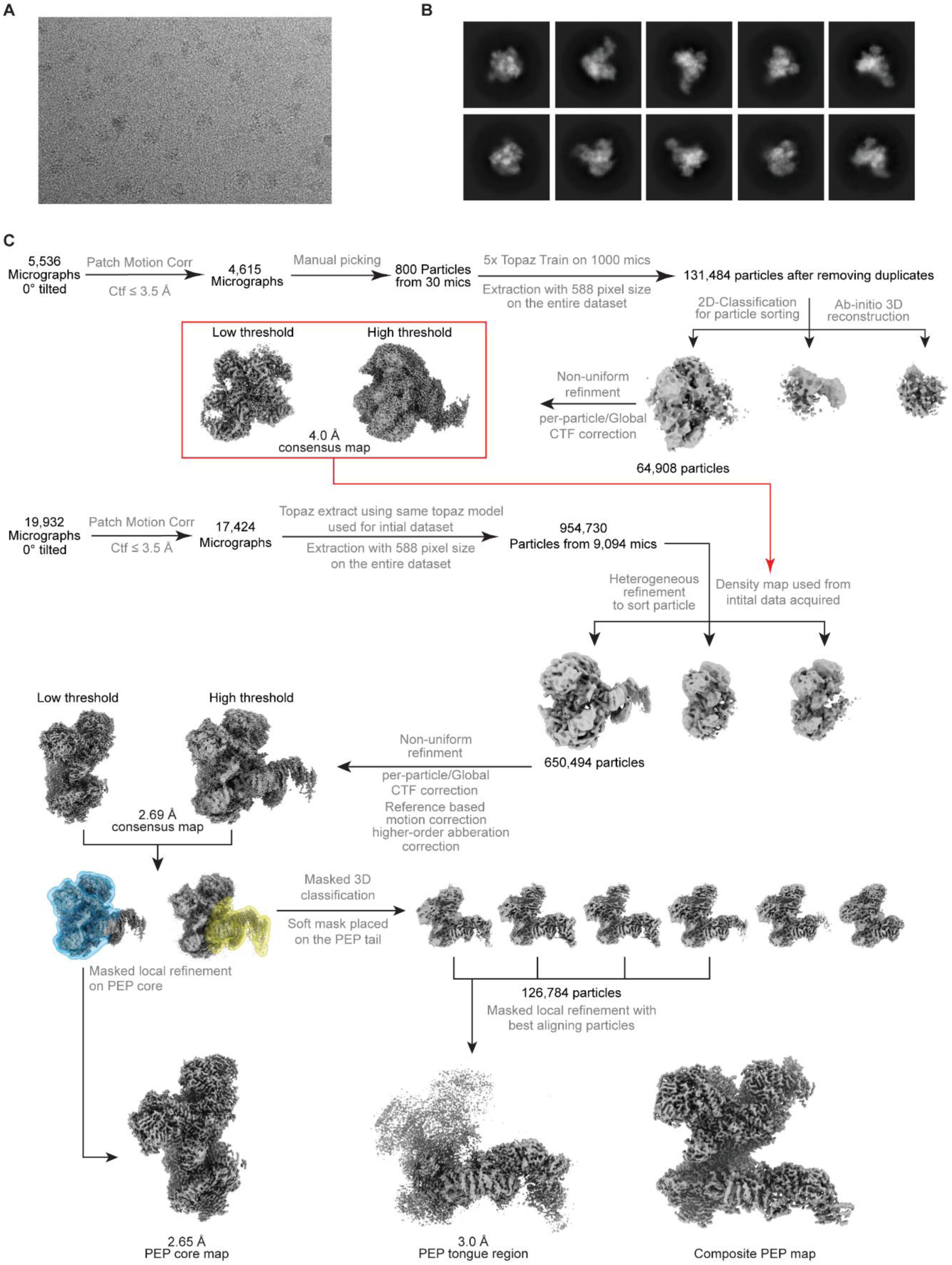
Cryo-EM data processing workflow for determining the structure of the CrPEP complex. **A**) Representative micrograph (total micrographs = 17,424) of the CrPEP vitrified after mild crosslinking (see Methods). (**B**) Reference-free 2D class averages from the datasets of CrPEP illustrating a range of particle orientations. (**C**) Workflow for processing and determining the structure of the CrPEP complex and its peripheral components (see Methods).

**Figure S5.**
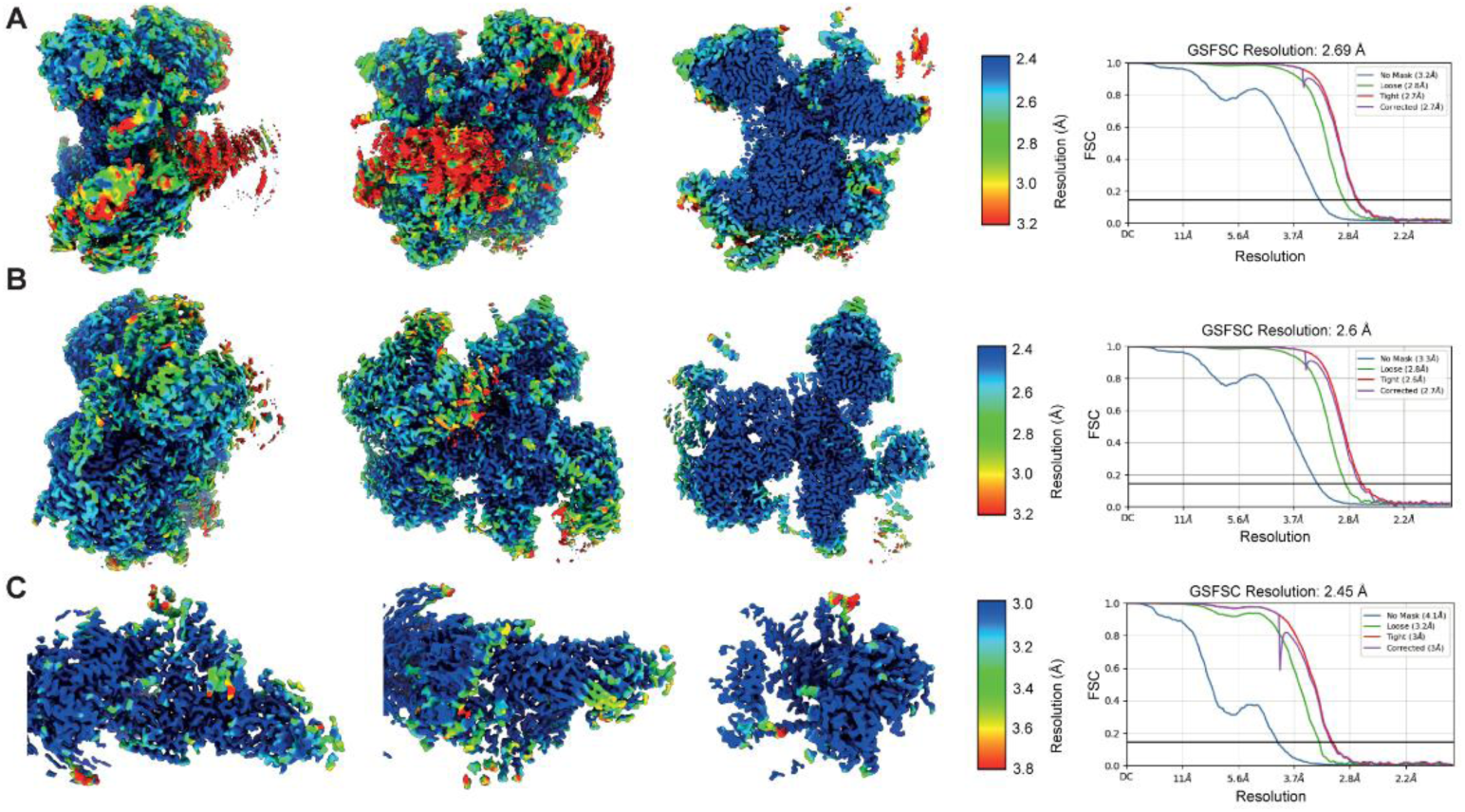
Cryo-EM map resolution estimation and quality. Cryo-EM maps coloured by the resolution estimated using gold-standard Fourier shell correlation (GSFSC). Two orthogonal views along with a cut open view of the of the center part (**A**) for the consensus map, (**B**) for the core map obtained after local refinement, and (**C**) for the tail region obtained after local refinement.

**Figure S6.**
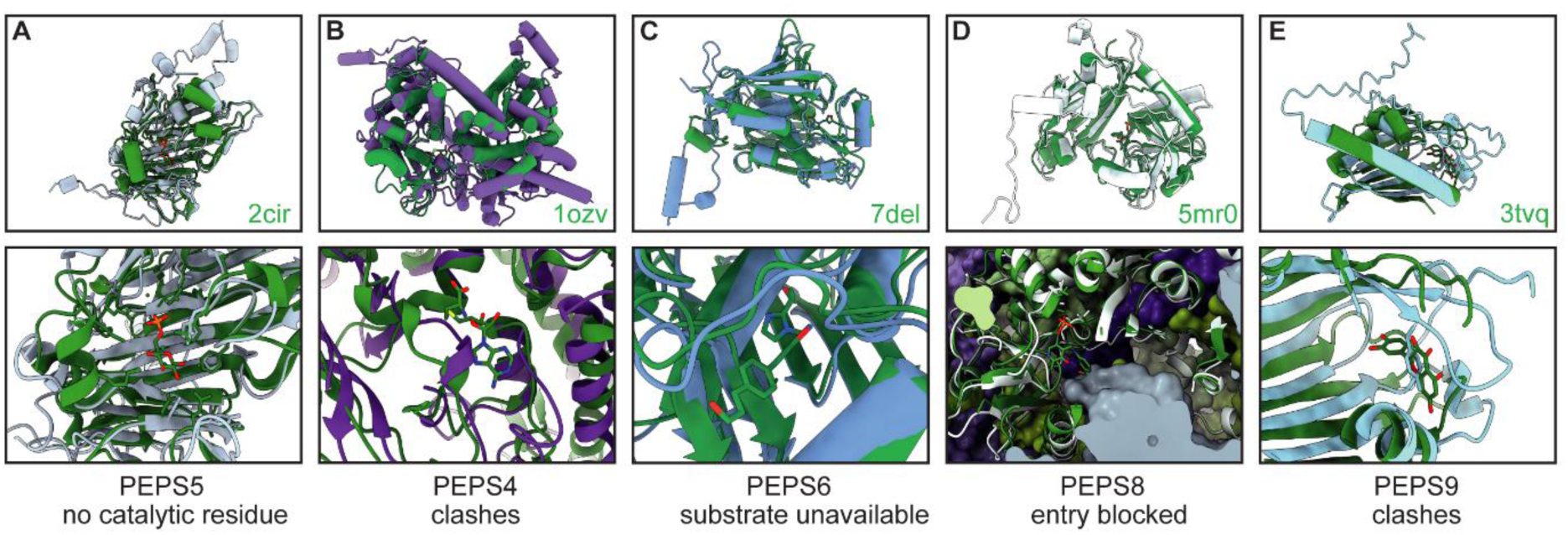
Superposition of selected PEPS with their respective enzymatic homologues displaying an overview and zoom in of the enzymatic active site. (**A**) Superposition of PEPS5 with Glucose-6-epimerase (pdb:2cir). (**B**) Superposition of PEPS4 with the SET domain of LSMT (pdb:1ozv). (**C**) Superposition of PEPS6 with UDP-3-O-acyl-N-acetylglucosamine deacetylase (pdb:7del) (**D**) Superposition of PEPS8 with branched-chain amino acid transaminase (pdb:5mr0) (**E**) Superposition of PEPS9 with transferase TCM Aro/Cyc (pdb:3tvq)

**Figure S7.**
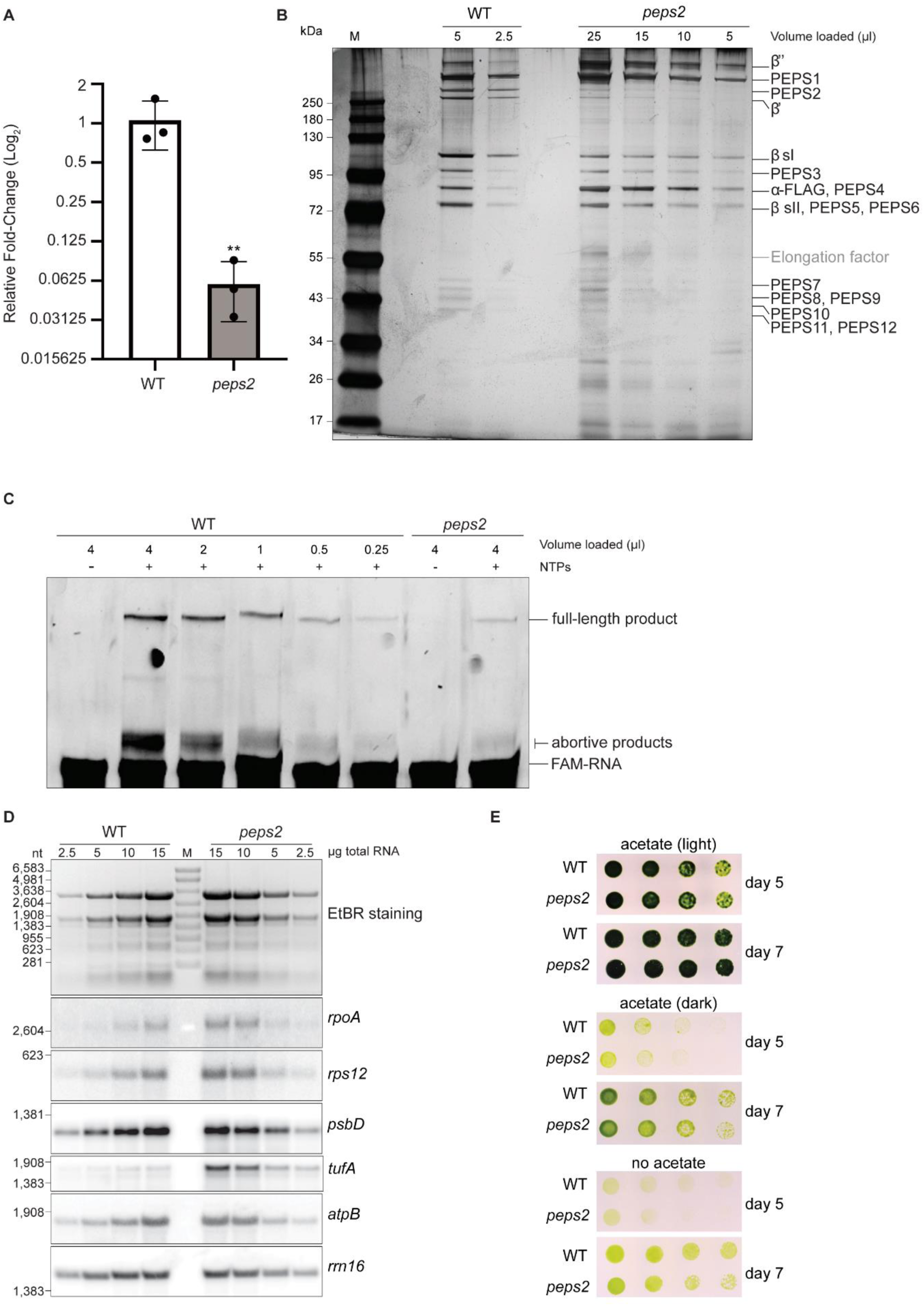
Biochemical and phenotypic analysis showing reduced CrPEP activity in the *peps2* mutant with no significant effects on chloroplast steady-state transcript levels or growth. (**A**) RT-qPCR analysis of *PEPS2* transcript levels in the *peps2* mutant background (**B**) SDS-PAGE analysis of serial dilutions of CrPEP purified from the wild-type and *peps2* mutant backgrounds, to assess loading in (C). (**C**) *In vitro* RNA extension assay comparing activity of CrPEP purified from the *peps2* mutant background to the wild-type background. FAM-RNA oligomer and extension products are indicated. (**D**) Northern Blot analysis of total RNA (2.5-15µg) extracted from wild-type and *peps2* mutant strains, using radioactive probes for chloroplast transcripts. (**E**) Phenotypic growth assay comparing growth of the *peps2* mutant and the wild type under heterotrophic and autotrophic conditions. Strains were spotted onto acetate-containing (TAP) or acetate-free minimal (HSM) agar plates in a three-fold serial dilution and imaged after either 5 or 7 days in either low-light (∼50 µmol photons m^−2^ s^−1^) or in the dark.

**Figure S8.**
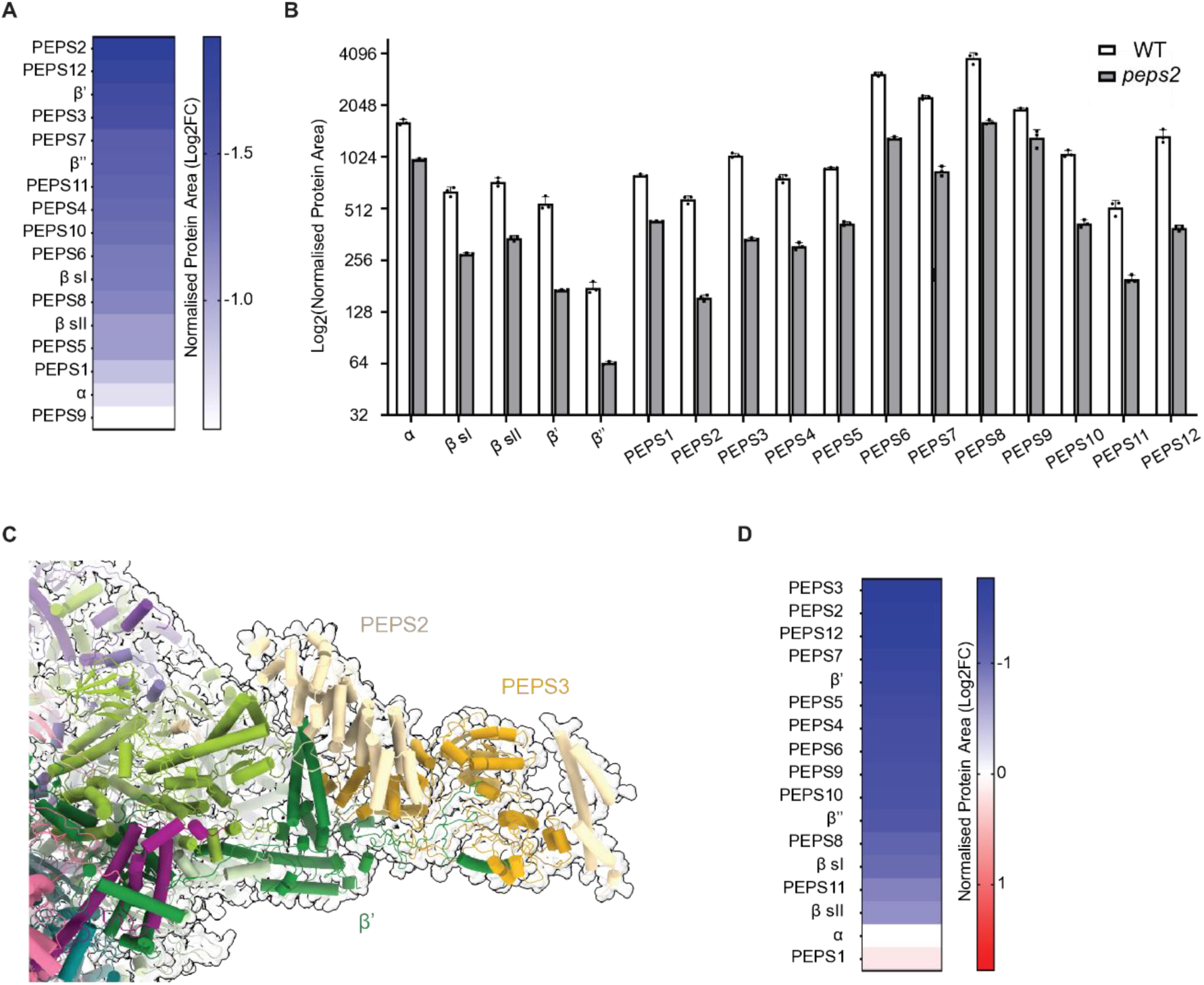
Mass spectrometry analysis of whole-cell lysate and eluate following α-FLAG immunoprecipitation from *peps2* mutant and wild-type strains. (**A**) Heatmap showing relative abundance of CrPEP components in whole-cell lysate from the *peps2* mutant compared to wild type. Protein area normalized via iBAQ, mean of three biological replicates. (**B**) Relative abundance of individual CrPEP subunits in whole-cell lysate from *peps2* and wild-type strains. Protein area normalized via iBAQ, mean of three biological replicates, log₂ scale. (**C**) Zoom-in on lower jaw of CrPEP complex, showing relative positions of PEPS2, PEPS3 and β’ subunits. (**D**) Heatmap showing relative enrichment of proteins co-immunoprecipitated with α-FLAG in the *peps2* mutant compared to the wild type. Protein area normalized to bait protein (α-FLAG).

**Figure S9.**
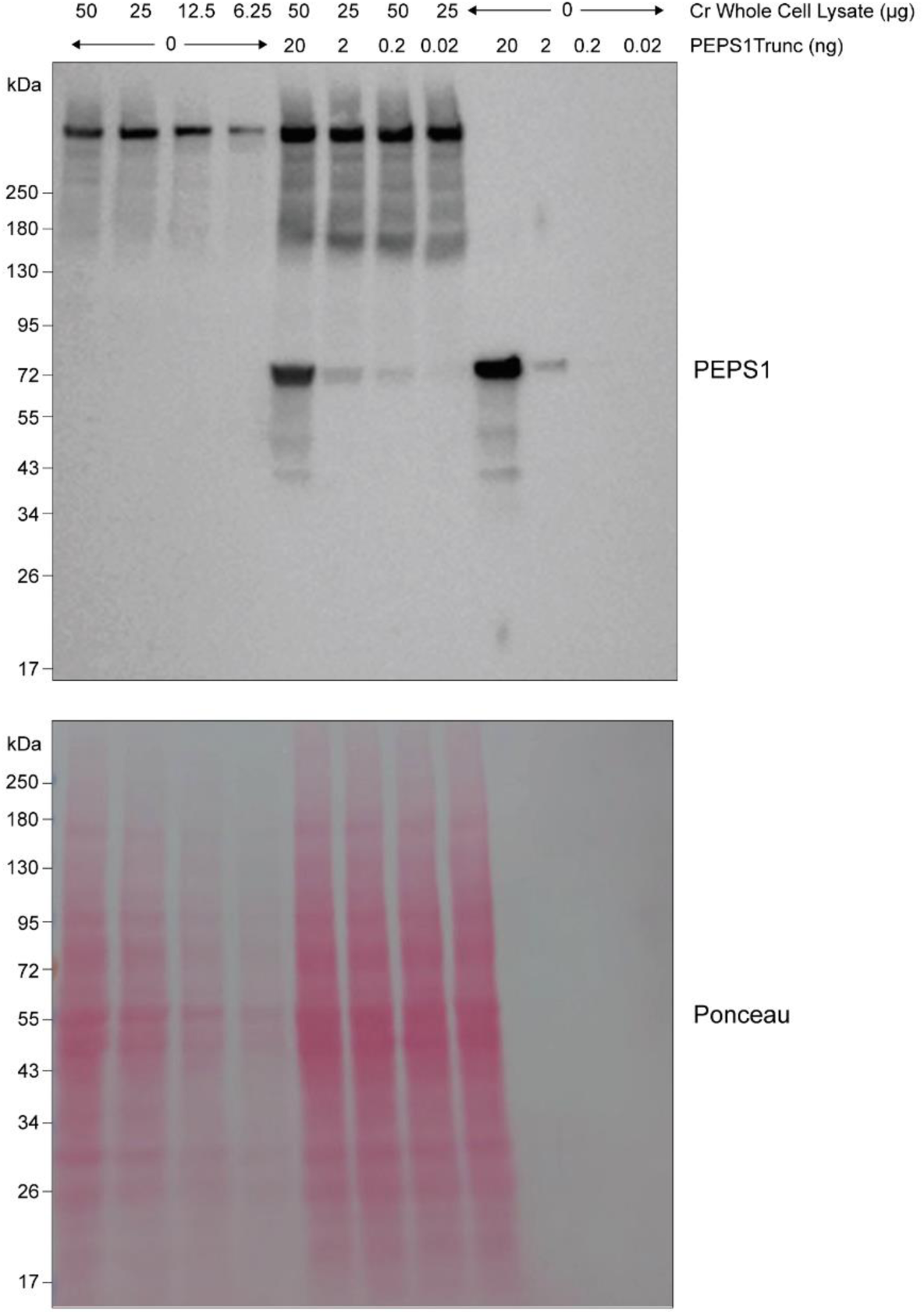
Immunoblot analysis verifying the specificity of the polyclonal PEPS1 antibody generated in this study. The antibody was tested against endogenous PEPS1 from *C. reinhardtii* wild-type lysate (280 kDa) and purified truncated PEPS1 protein (PEPS1Trunc; residues 1-580aa; 54.9 kDa), related to Figure 4

**Figure S10.**
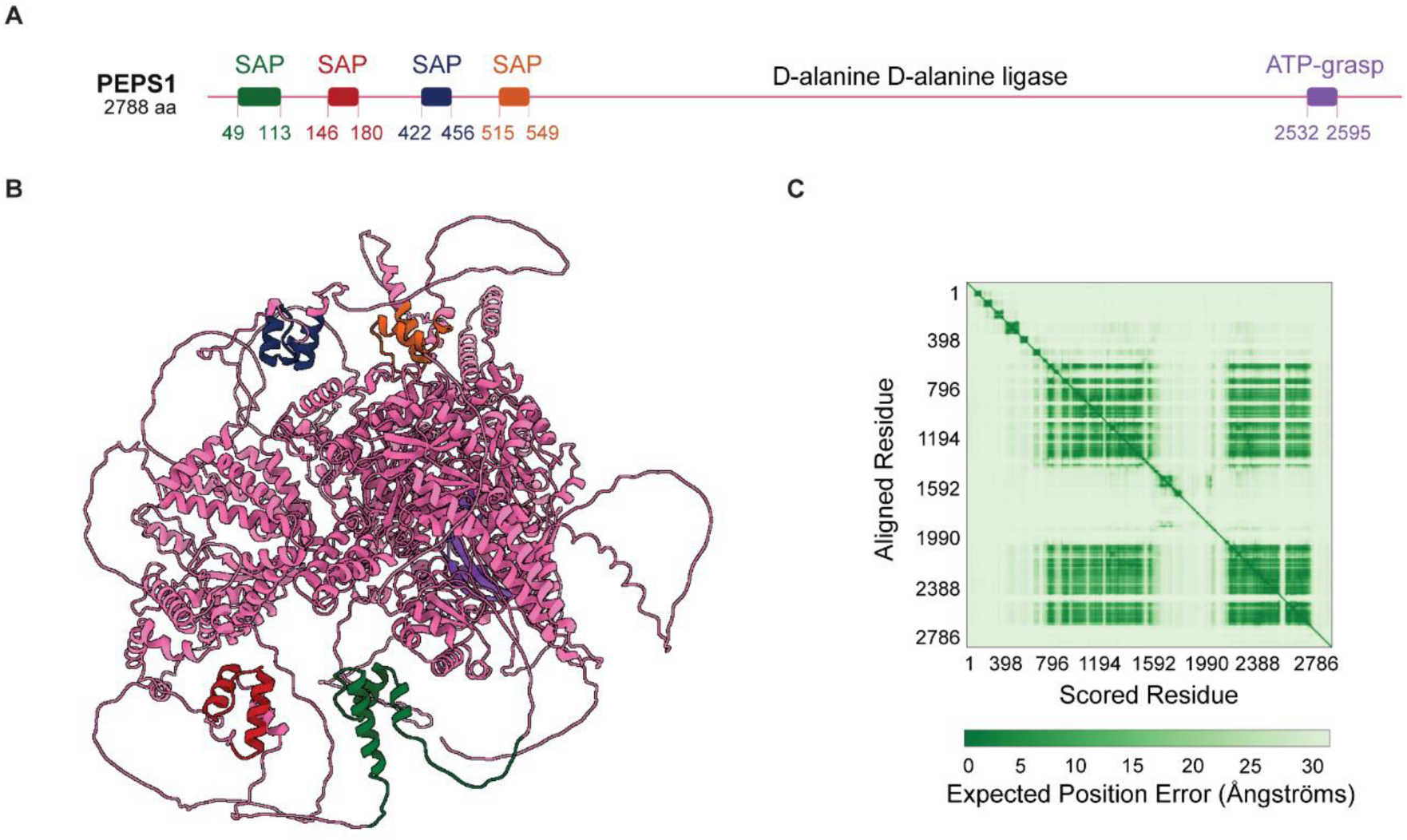
Positioning and predicted structure of the four N-terminal SAP domains in PEPS1. (**A**) Schematic representation of the full-length PEPS1 protein, showing the positions of the four N-terminal SAP domains. (**B**) AlphaFold3 prediction of the full-length PEPS1 protein with SAP domains colour-coded to match schematic in (A). Overall pTM score = 0.55 (**C**) Predicted Aligned Error (PAE) plot for the AlphaFold3 prediction of PEPS1 in (B).

**Figure S11.**
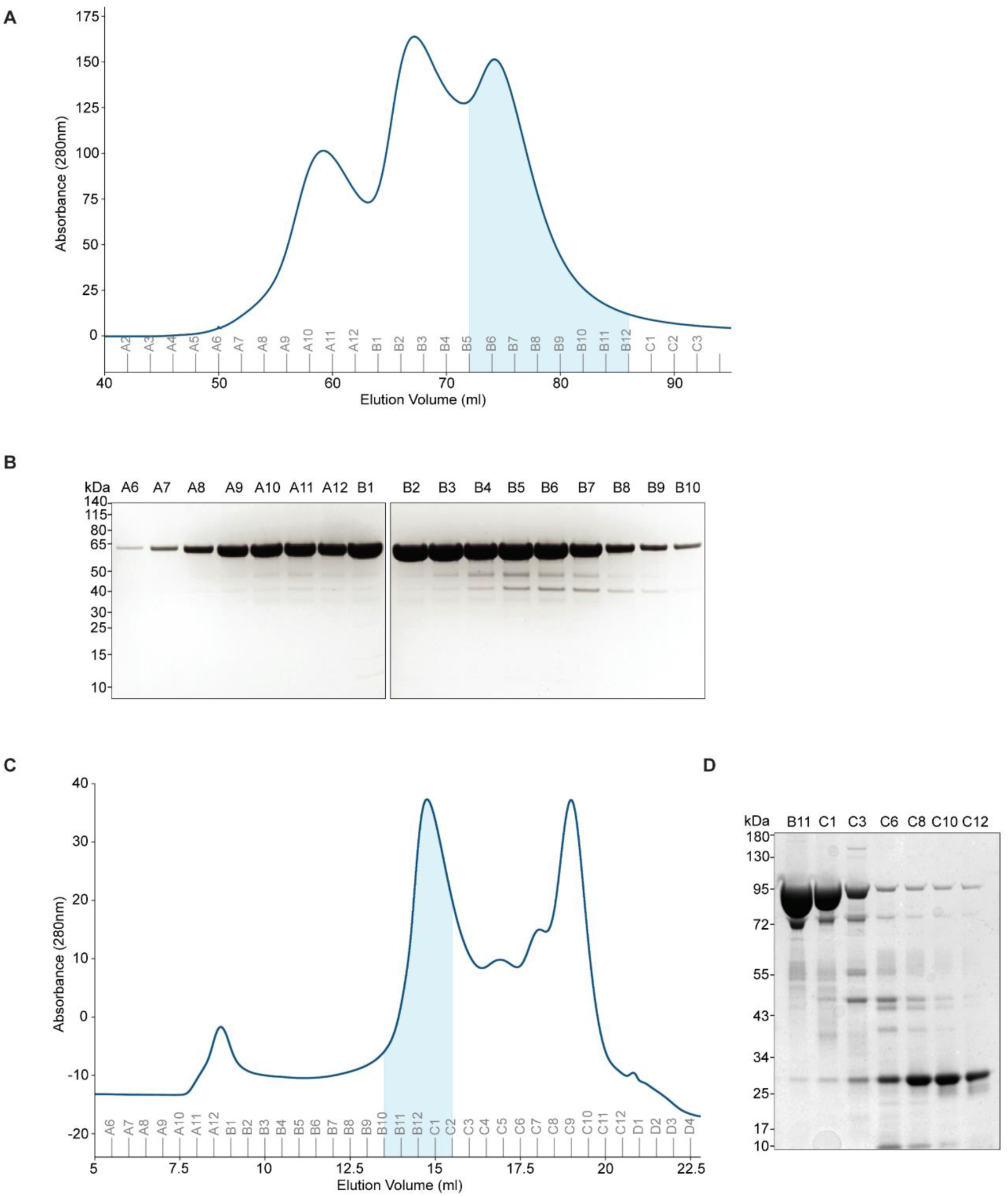
Purification of recombinant PEPS1Trunc and Cr σ factor used in EMSA. (**A**) Size-exclusion chromatogram at 280 nm following elution of the PEPS1Trunc protein (54.9 kDa) in 2 ml fractions after separation on a Superdex 200 16/60 column (Cytiva). The shaded region corresponds to the fractions pooled and concentrated for downstream use in EMSA. (**B**) SDS-PAGE analysis of peak fractions identified in size-exclusion chromatography (SEC) for PEPS1Trunc. Fractions B5-B12 (likely representing the lower molecular weight monomer) were pooled and concentrated for use in EMSA. (**C**) Size-exclusion chromatogram at 280 nm following elution of the Cr σ factor protein (84.1 kDa) in 500 µl fractions after separation on a Superose 6 Increase 10/300 column (Cytiva). The shaded region corresponds to the fractions pooled and concentrated for downstream use in EMSA. (**D**) SDS-PAGE analysis of peak fractions identified in SEC for Cr σ factor. Fractions B10-C2 were pooled and concentrated for downstream use in EMSA.

**Table S1.**
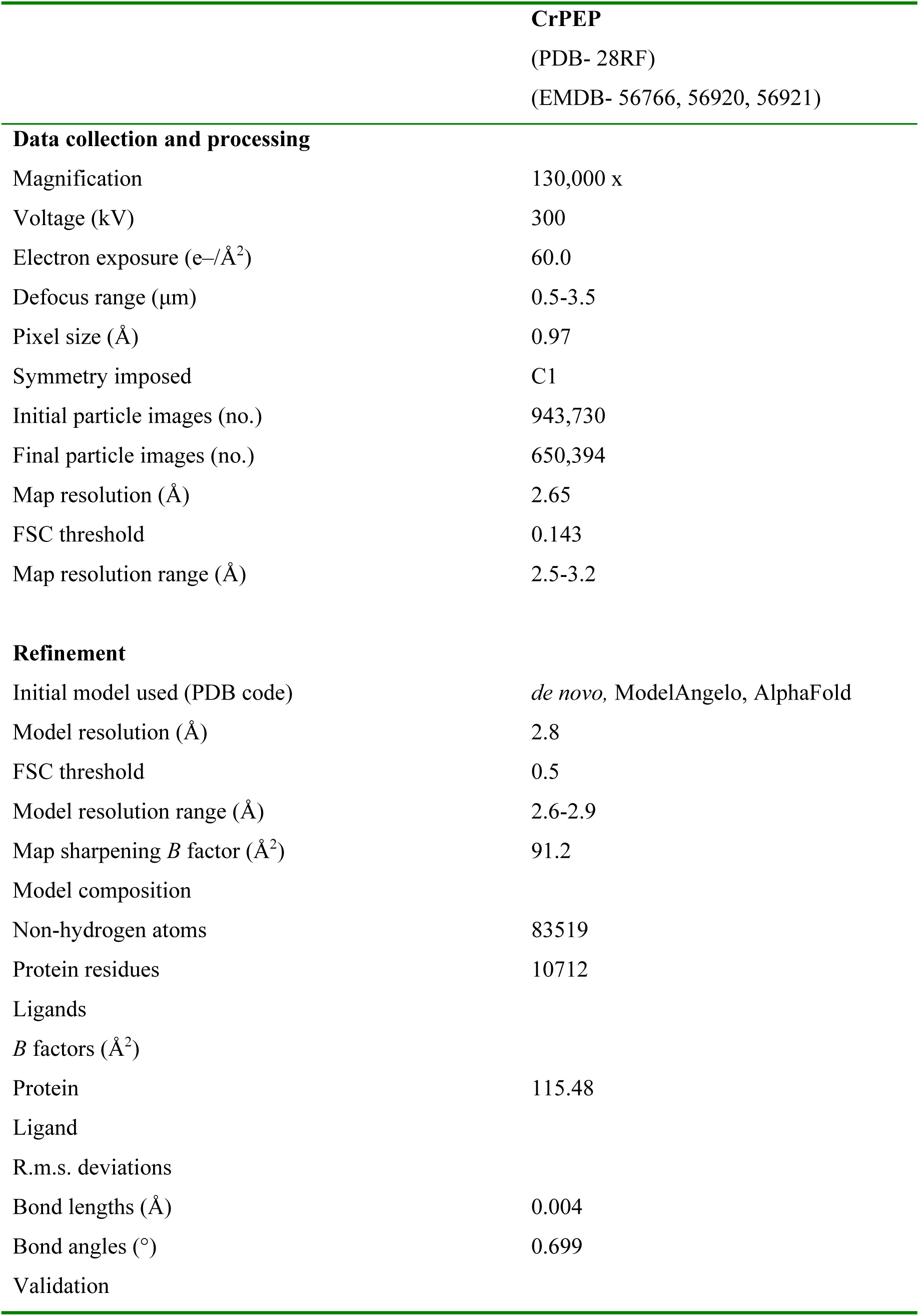
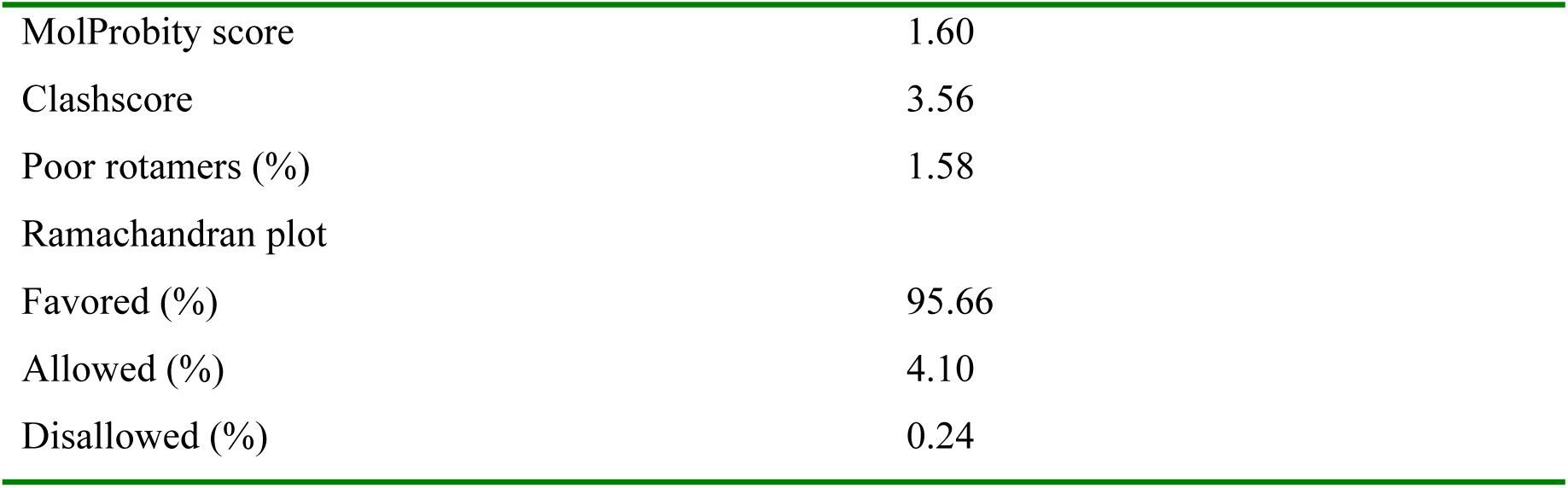
Cryo-EM data collection, refinement and validation statistics.

|  | **CrPEP**<br>(PDB- 28RF)<br>(EMDB- 56766, 56920, 56921) |
| --- | --- |
| **Data collection and processing** |  |
| Magnification | 130,000 x |
| Voltage (kV) | 300 |
| Electron exposure (e-/Å<sup>2</sup>) | 60.0 |
| Defocus range (μm) | 0.5-3.5 |
| Pixel size (Å) | 0.97 |
| Symmetry imposed | C1 |
| Initial particle images (no.) | 943,730 |
| Final particle images (no.) | 650,394 |
| Map resolution (Å) | 2.65 |
| FSC threshold | 0.143 |
| Map resolution range (Å) | 2.5-3.2 |
| **Refinement** |  |
| Initial model used (PDB code) | *de novo*, ModelAngelo, AlphaFold |
| Model resolution (Å) | 2.8 |
| FSC threshold | 0.5 |
| Model resolution range (Å) | 2.6-2.9 |
| Map sharpening *B* factor (Å<sup>2</sup>) | 91.2 |
| Model composition |  |
| Non-hydrogen atoms | 83519 |
| Protein residues | 10712 |
| Ligands |  |
| *B* factors (Å<sup>2</sup>) |  |
| Protein | 115.48 |
| Ligand |  |
| R.m.s. deviations |  |
| Bond lengths (Å) | 0.004 |
| Bond angles (°) | 0.699 |
| Validation |  |
| MolProbity score | 1.60 |
| Clashscore | 3.56 |
| Poor rotamers (%) | 1.58 |
| Ramachandran plot |  |
| Favored (%) | 95.66 |
| Allowed (%) | 4.10 |
| Disallowed (%) | 0.24 |

**Table S2.** Subunit composition of CrPEP.

| Subunit | Alternative Name | Gene Accession Number (Phytozome) | UniProt ID | Chain ID in CrPEP | Stoichiometric Ratio in CrPEP |
| --- | --- | --- | --- | --- | --- |
| $\alpha$ | RpoA | CreCp.g802298 | A0A218N9C3 | A, Z | 2 |
| $\beta$ sI | RpoB1 | CreCp.g802306 | A8HT91 | B | 1 |
| $\beta$ sII | RpoB2 | CreCp.g802305 | Q8HTL7 | C | 1 |
| $\beta'$ | RpoC1 | CreCp.g802335 | A0A218N8D6 | E | 1 |
| $\beta''$ | RpoC2 | CreCp.g802311 | Q7PCJ6 | D | 1 |
| PEPS1 |  | Cre15.g635650 | A0A2K3CWF2 | K | 1 |
| PEPS2 |  | Cre12.g497350 | A0A2K3D3G3 | M | 1 |
| PEPS3 | MurE | Cre12.g519900 | A0A2K3D404 | N | 1 |
| PEPS4 |  | Cre09.g388500 | A0A2K3DDP1 | G | 1 |
| PEPS5 |  | Cre05.g234651 | A0A2K3DSM7 | F | 1 |
| PEPS6 |  | Cre16.g653350 | A0A2K3CSZ9 | H, I | 2 |
| PEPS7 |  | Cre16.g649350 | A0A2K3CST8 | T | 1 |
| PEPS8 |  | Cre13.g576400 | A8HT91 | X, Y | 2 |
| PEPS9 |  | Cre09.g400627 | A0A2K3DEZ2 | Q | 1 |
| PEPS10 |  | Cre07.g348500 | A0A2K3DL30 | R | 1 |
| PEPS11 |  | Cre01.g024150 | A8HQ14 | S | 1 |
| PEPS12 |  | Cre05.g234050 | A8JG54 | J | 1 |

**Table S3.** Phylogenetic distribution of Chlamydomonas PEPS proteins and corresponding *A. thaliana* orthologs/homologs.

| Protein | Protein Accession Number (Phytozome) | Category | Distribution | Orthology/ Homology in *Arabidopsis* | Ortholog/Homolog Trivial Name |
| --- | --- | --- | --- | --- | --- |
| PEPS1 | Cre15.g635650.t1.2 | Novel | Chloroplastida | Ortholog to AT3G08840.7 | D-alanine-D-alanine ligase family |
| PEPS2 | Cre12.g497350.t1.1 | PAP | Chloroplastida | Ortholog to AT3G04260.1 | PAP1/pTAC3 |
| PEPS3 | Cre12.g519900.t1.2 | PAP | Chloroplastida | Ortholog to AT1G63680.2 | PAP11//MurE |
| PEPS4 | Cre09.g388500.t1.2 | Novel | Chlorophytes | - | - |
| PEPS5 | Cre05.g234651.t1.1 | Novel | Chloroplastida | Homolog to AT5G66530.3 | Galactose mutarotase-like superfamily protein |
| PEPS6 | Cre16.g653350.t1.2 | Novel | Chloroplastida | Ortholog to AT1G24793.1, AT1G25054.1, AT1G25145.1, AT1G25210.2 | LPXC1, LPXC3, LPXC4, LPXC5 |
| PEPS7 | Cre16.g649350.t1.1 | Novel | Chlamydomonas/ Chlorophytes | - | - |
| PEPS8 | Cre13.g576400.t1.2 | Novel | Chloroplastida | Homolog to AT5G57850.1, AT3G05190.2, AT5G27410.2 | DAT1, D-aminoacid aminotransferase-like PLP-dependent enzymes superfamily, D-aminoacid aminotransferase-like PLP-dependent enzymes superfamily protein |
| PEPS9 | Cre09.g400627.t1.1 | Novel | Chloroplastida | No hit in At | - |
| PEPS10 | Cre07.g348500.t1.1 | Novel | Chloroplastida | No hit in At | - |
| PEPS11 | Cre01.g024150.t1.2 | Novel | Chlorophytes | - | - |
| PEPS12 | Cre05.g234050.t1.2 | Novel | Chlamydomonas/ Chlorophytes | - | - |

### Materials and Methods

#### Strains and Growth Conditions

All *C. reinhardtii* strains used in this study were maintained on Tris-Acetate-Phosphate (TAP) solid media containing 1.6% (w/v) agar (USP grade, ThermoFisher) and Hutner’s trace elements (*60*) at 22°C in low light (∼50 µmol photons m^−2^ s^−1^). For chloroplast transformant strains the medium was supplemented with 100 µg ml^-1^ spectinomycin. For the PEPS3, PEPS9, and PEPS12-Venus-3xFLAG strains and the *peps2* mutant, the medium was supplemented with 10 µg ml^-1^ paromomycin. Generally, during liquid growth no antibiotic was supplemented, and cultures were grown at 22°C with gentle mixing in low light (∼50 µmol photons m^−2^ s^−1^). When strains were grown on solid medium lacking acetate, High-Salt Medium (HSM) (*61*) supplemented with 1.6% (w/v) agar was used in place of TAP. All strains used in this study are listed in Table S4.

**Table S4:** Chlamydomonas strains used in this study.

| **Internal Strain Reference** | **Strain Name** | **Description** | **Reference** |
| --- | --- | --- | --- |
| 166 | CC-4533 | Background strain for strains 651,959, and 960 | Available at the Chlamydomonas Resource Center |
| 307 | *rpoA*-FLAG | Wild type transformed with *rpoA*-FLAG transgene | (31) |
| 651 | *peps2-1* | Insertional mutant of Cre12.g497350 | Available at the Chlamydomonas Resource Center Clip ID: LMJ.RY0402.219638 |
| 959 | CC-4533:*rpoA*-FLAG | CC-4533 transformed with *rpoA*-FLAG transgene | Generated in this study |
| 960 | *peps2*:*rpoA*-FLAG | *peps2* mutant transformed with *rpoA*-FLAG transgene | Generated in this study |
| 456 | CSI_LW07H6 (PEPS3-Venus-3xFLAG) | CC-4533 expressing Cre12.g519900-Venus-3xFLAG transgene | Available at the Chlamydomonas Resource Center |
| 457 | CSI_LW05F7 (PEPS9-Venus-3xFLAG) | CC-4533 expressing Cre09.g400627-Venus-3xFLAG transgene | Available at the Chlamydomonas Resource Center |
| 647 | CSI_LW03B3 (PEPS12-Venus-3xFLAG) | CC-4533 expressing Cre05.g234050-Venus-3xFLAG-transgene | Available at the Chlamydomonas Resource Center |

#### Chloroplast Transformation

Chloroplast biolistic transformations for the generation of strains 959 and 960 were performed as previously described in (*31*). In brief, cells to be transformed were grown to a density of 2-8× 10^6^ cells ml^−1^ and plated onto TAP solid medium supplemented with 100 µg ml^-1^ spectinomycin. Plates were then bombarded with 550-nm-diameter gold particles (Seashell Technology S550d) coated with 1 µg of pRAM68.13 plasmid (Table S5). After 1 week of growth at 25°C in constant light, singles colonies were picked, replated a minimum of four times on TAP medium supplemented with 100 µg/ml spectinomycin, and screened for transgene expression via immunoblot. Homoplasmic status of the selected transformant strain was confirmed via PCR analysis.

**Table S5:** Plasmids used in this study.

| Plasmid | Used For | Reference |
| --- | --- | --- |
| pRAM68.13 | Biolistic transformation of *rpoA*-FLAG construct | (31) |
| pRAM366 | Insect-cell expression of Cr $\sigma$ factor (*RPOD1*) | Generated in this study |
| pRAM457 | *E. coli* expression of PEPS1Trunc | Generated in this study |

#### Purification of CrPEP

Four to eight-liter liquid cultures of the relevant FLAG-tagged strain were grown in TAP medium to a density of 2-8× 10^6^ cells ml^−1^. Cells were collected at 4°C by centrifugation at 3000 x g for 10 min, and the pellet, kept at 4°C, quickly resuspended in an equal volume of 2X Lysis buffer (100 mM HEPES, 100 mM KOAc, 4 mM Mg(OAc)_2_ 4H_2_O, 2 mM CaCl_2_, 30% glycerol (v/v), pH adjusted to 6.8 with KOH, supplemented freshly with two tablets per 25 ml of EDTA-Free Protease Inhibitor Cocktail (Roche)). The resuspended cell pellet was then snap-frozen in liquid nitrogen and immediately used for immunoprecipitation of the CrPEP complex. The frozen cells were lysed using a Cryomill (Retsch) at a frequency of 30 oscillations per second for 3 cycles of 1 min each, with 3 min cooldown in liquid nitrogen between each cycle. The resultant cell powder was then thawed on ice for approx. 1 hour. The lysate was then resuspended in an equal volume of 2X Lysis buffer supplemented with 0.2% NP-40 Alternative (Sigma-Aldrich) and incubated on a rolling incubator at 4°C for 45 min. Solubilized material was clarified by centrifugation at 17,800 × g for 30 min at 4°C, and the supernatant transferred to a fresh tube. ANTI-FLAG M2 Affinity Gel agarose beads (Sigma-Aldrich) were added (100 µl per 30 ml supernatant) and left to incubate overnight at 4°C on a rolling incubator.

Where two strains are compared, the total protein content of the soluble fraction from each strain was quantified by BCA assay. Supernatants were then diluted in 2X Lysis buffer supplemented with NP-40 Alternative and protease inhibitor to ensure the beads were incubated in equal volumes and equivalent total protein concentration.

Following overnight incubation, the agarose beads were collected on a 20-ml gravity flow column (Bio-Rad) and subjected to five 6 min washes in 1 ml 1X Lysis buffer supplemented with 0.005% NP-40 Alternative. The beads were then resuspended in 150 µl of Elution Buffer (50 mM HEPES, 150 mM NaCl, pH 7.5), supplied with 300 ng µl^-1^ of FLAG peptide, and incubated for 30 mins. This elution step was repeated three times in total. Each eluate was analysed via SDS-PAGE, followed by Coomassie Blue staining (see Total Protein Staining). Elutions of similar concentration were pooled. Aliquots for mass photometry were saved at 4°C, with the remaining sample snap-frozen on liquid nitrogen for storage at −70°C.

#### Total Protein Staining

Following SDS–PAGE, gels were stained either with Coomassie or with silver stain, depending on the desired sensitivity.

For Coomassie staining, gels were briefly rinsed in deionized water and then incubated overnight in staining solution (6.4% ethanol, 3.75% phosphoric acid, 1% β-cyclodextrin, 0.05% Coomassie Brilliant Blue G; Sigma-Aldrich). Following staining, gels were destained for 1 min in 10% acetic acid and 25% methanol, then for >20 min in 25% methanol, before imaging on an iBright FL 1500 system (ThermoFisher).

For silver staining, the ProteoSilver Silver Stain Kit (Sigma-Aldrich) was used according to the manufacturer’s instructions for direct silver staining, with development times of <7 min being sufficient for visualization. Imaging was performed using the iBright FL 1500 system (ThermoFisher).

#### Mass photometry

To remove the FLAG peptide and prepare sample for mass photometry, CrPEP eluate was concentrated using a centrifugal concentrator (Amicon-Ultra 500 100 kDa MWCO, Merck) to approximately 1 mg ml^-1^ (∼500 nM).

Measurements were made using the Refeyn TwoMP mass photometer. The instrument was calibrated using selected masses of NativeMark Unstained Protein Standard (146, 480 and 1048 kDa, ThermoFisher). CrPEP Eluate was diluted at a 1:40 ratio for mass photometry analysis. Binding events were recorded for 60 s using the AcquireMP software v2024 R2, and analysis conducted using DiscoveryMP v2024 R2. Mass of CrPEP and any associated subcomplexes were estimated as the mode of the histogram distribution following fit of a Gaussian distribution.

#### Cryo-EM sample preparation

To obtain the cryo-EM structure of CrPEP, freshly purified protein was concentrated to 2 mg ml⁻¹. A 4 µl aliquot was deposited onto glow-discharged Quantifoil grids, blotted at force 4 for 3.5 s, and plunge-frozen in liquid ethane cooled by liquid nitrogen using a Vitrobot Mark III (Thermo Fisher) operating at 100% humidity and 4°C. Initial grid screening revealed dissociation of the complex due to its contact with the air–water interface during vitrification. To circumvent the issue and to further stabilize the CrPEP complex, mild chemical cross-linking was performed using BS3 (bis(sulfosuccinimidyl) suberate). The eluted CrPEP complex was adjusted to 2 mg ml⁻¹. Cross-linking was initiated by adding BS3 to a final concentration of 1 mM, followed by incubation for 30 min at room temperature. The reaction was quenched with 50 mM Tris, and residual cross-linker was removed by buffer exchange into the elution buffer (see above) using a concentrator. The protein complex was then reconcentrated to 2 mg ml^-1^ for grid vitrification. For vitrification, 3 µl of the cross-linked complex was mixed with 1 µl of 6 mM CHAPSO immediately before application to glow-discharged Quantifoil grids. Grids were blotted at force 6 for 6 s and plunge-frozen in liquid ethane using the Vitrobot Mark III at 100% humidity and 4°C.

#### Cryo-EM data acquisition and processing

To assess the assembly and orientation of the protein complex, an initial screening dataset was collected on a JEOL CryoArm200 microscope operating at 200 keV and equipped with a Gatan K3 Summit direct electron detector. Automated data acquisition was performed using SerialEM (*62*) at a magnification of 60,000×, with a total electron dose of 50 e⁻ Å⁻² fractionated into 50 frames at a calibrated pixel size of 0.85 Å. In total, 5,536 micrographs were recorded. The dataset was processed in CryoSPARC (*63*). After import, the movie frames were gain-normalized, corrected for beam-induced motion using Patch Motion correction (with dose-weighting) (*64*), and subjected to contrast transfer function (CTF) estimation. Micrographs with a CTF fit worse than 3.5 Å were discarded, resulting in 4,615 images for further analysis. Particles were initially picked manually and subsequently used to train several rounds of Topaz-based particle picking (*65*). Using the trained Topaz model, 131,484 particles were extracted for generation of three ab initio models. Of these, 64,908 particles contributed to a well-defined ab initio class, which was then refined using non-uniform refinement. This yielded a 3D reconstruction at 4 Å resolution. Although the resolution remained moderate, the screening dataset confirmed that the protein complex was intact and exhibited isotropic particle orientation.

To obtain a high-resolution structure of CrPEP, vitrified grids were transferred to a Thermo Fisher Titan Krios G4 transmission electron microscope operating at 300 keV and equipped with a Falcon 4i direct electron detector and a Selectris energy filter with a 10 eV slit width. Automated data acquisition was carried out using smart EPU software, with movie frames recorded at a magnification of 130,000×, corresponding to a calibrated pixel size of 0.97 Å. Images were collected in counting mode with a total electron dose of 60 e⁻ Å⁻² and saved in electron-event representation (EER) format. Due to the low abundance of particles on the grids, a total of 19,932 micrographs were collected. The dataset was imported into CryoSPARC, where EER movies were fractionated into 60 frames, followed by patch motion correction and patch CTF estimation. Micrographs with a CTF fit worse than 3.5 Å were rejected for further processing, leaving a total of 17,424 micrographs for downstream processing. For initial particle picking, we applied the Topaz model previously trained on the screening dataset to a small subset of the high-resolution micrographs. Extracted particles were subjected to 2D classification to identify well-aligned classes, which were then used for iterative retraining of the Topaz model until a robust picker was obtained for the full dataset. After particle extraction, heterogeneous refinement with three classes was performed using the screening dataset reconstruction as an initial reference. This yielded three models: one representing the full CrPEP assembly (core + tail) and two in which only the core region was resolved. Particles corresponding to the fully assembled complex were re-extracted with a box size of 588 pixels and refined using non-uniform (NU) refinement, resulting in a 3 Å reconstruction. To further improve the resolution, particles underwent reference-based motion correction (polishing) and higher-order aberration correction using beam-shift parameters. These steps improved the NU-refined map to 2.7 Å resolution. While the core region was well resolved, the extended tail displayed lower map quality. To enhance the tail density, a soft mask was applied to the tail region, followed by 3D masked classification into six classes. Four classes retained clear tail density and were selected for further processing. Particles from these classes were subjected to local refinement, yielding a 3 Å map with improved tail features. An analogous focused refinement strategy applied to the core region further improved its resolution to 2.65 Å.

The initial model of the CrPEP complex from *C. reinhardtii* was generated using ModelAngelo (*66*). Based on this preliminary reconstruction, all subunits contributing to the mega-assembly were identified by BLAST analysis of their sequences on the NCBI server. To enrich and refine the ModelAngelo output, individual AlphaFold2 (*67*) homology models were generated for each subunit and rigid-body fitted into the electron density using UCSF Chimera and Coot. Model rebuilding and polishing were performed in Coot (v.0.9.8.92), followed by real-space refinement in PHENIX (v1.21-5207). Structural illustrations were prepared using UCSF Chimera (v1.17.3), UCSF ChimeraX (v1.6.1), and PyMOL.

#### Confocal Microscopy

For confocal imaging, 4 ml cultures of the relevant strains were grown in TAP liquid medium for 5 days. 24 hours before imaging, cells were pelleted via centrifugation at 1500 x g for 15 min at RT and resuspended in an equal volume of High-Salt-Minimal (HSM) media (*61*). On the day of imaging, cells were pelleted again, and the cell pellet washed with 1X Phosphate-Buffered Saline (PBS). Cells were then fixed via incubation for 15 min in 3.7% PFA, followed by a further washing with 1X PBS. Prior to observation, cells were subjected to 15 min of staining with 0.5 µg ml^-1^ DAPI (ThermoScientific) in the dark. All imaging was conducted using a LSM980 Axio Observer Confocal Microscope (Zeiss) with a 63X oil lens, and a total of 30 images collected per strain. All raw images are available through Figshare: (https://figshare.com/s/a761003830830efa0663).

#### Homoplasmicity, RT-qPCR, and RNA Gel Blot Analysis

Chloroplast homoplasmicity of all *rpoA*-FLAG transformants was verified via PCR as shown in Fig. S1, following the method previously described in (*31*). All relevant primers are listed in Table S6. Genomic DNA was extracted using the CTAB method from each *rpoA*-FLAG transformant and the parental background strain. In addition, a plasmid carrying the *psbD5′rpoA* chimeric gene (pRAM68.13) was included as a positive control, since it contains the exact construct expected to integrate into the chloroplast genome. PCR amplification with primers specific for the chimeric *psbD5’rpoA* locus (psbD-47/SR218, Table S6) confirmed the presence of the *rpoA*-FLAG construct in transformants, with the plasmid serving as a reference for the expected amplicon. As a negative control, primers targeting the wild-type *rpoA*5’*rpoA* locus (SR218/SR241) were used to amplify only the endogenous genomic sequence. To ensure that any residual wild-type DNA in the transformants could be detected, 1% (w/w) genomic DNA from the parental background strain was mixed into the PCR reactions as a sensitivity control. This dilution demonstrated that even a 1:100 ratio of wild-type to transformant DNA was sufficient for reliable detection of the wild-type *rpoA* locus, thereby confirming that absence of a wild-type signal in the transformants reflected true homoplasmicity.

**Table S6:**
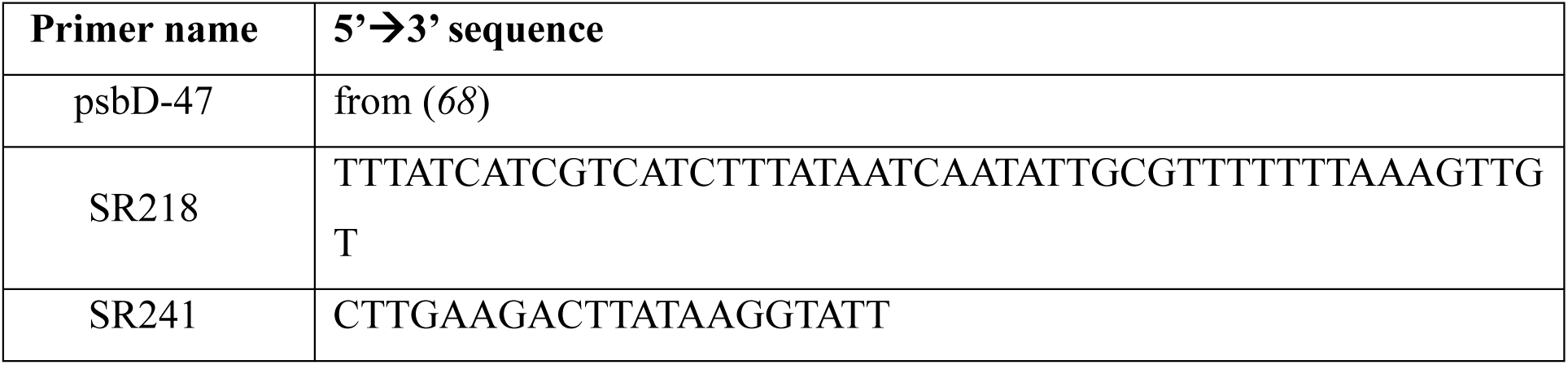
Primers used to assess homoplasmicity of *rpoA*-FLAG transformants.

#### RNA extraction

Total RNA was extracted with minimal adjustment to that previously reported in (*69*). In brief: 5-10 ml cell culture in early-mid exponential phase (1-5x10^6^ ml^-1^) was harvested by pelleting 5 min 3000 x g at RT. After decanting the media, 1 ml of TRI Reagent (Zymo Research) was added to the cell pellet, and complete lysis was achieved by vortexing for 5-10 min at 4°C. One-fifth volume of chloroform was added to the lysate, and tubes were shaken by hand for 1 min. Samples were then centrifuged for 7 min, 11000 x g at RT and the upper phase transferred to new nuclease-free tubes. After the addition of 1 volume of 100% ethanol and mixing by pipetting, samples were loaded in the columns of the Direct-zol RNA miniprep kit (ZymoResearch) and the manufacturer’s protocol was followed, using in-column DNA digestion. RNA concentration was quantified using a Nanodrop ND-100 Spectrophotometer (ThermoScientific).

#### cDNA Synthesis and RT-qPCR for *peps2* transcripts

One microgram of RNA was used for reverse transcription with the Maxima First Strand cDNA Synthesis Kit for RT-qPCR (ThermoScientific), according to the manafacturer’s protocol. After addition of the reverse transcriptase enzyme and the supplied 5X Reaction Mix (containing dNTPs, random hexamer primers, oligo(dT)_18_ and RNase inhibitor), samples were incubated in the following conditions: 10 min at 25°C, 30 min at 65°C (to accommodate the high GC content of the Chlamydomonas genome) and 5 min at 85°C to inactivate the enzyme. The resultant cDNA was diluted five-fold in nuclease-free water prior to use. RT-qPCR reactions were carried out using FastStart Essential DNA Green Master (Roche) according to the manufacturer’s protocol, with a final volume of 15 µl per sample. Each genotype was tested as three technical replicates, and a non-template control was used for each primer combination. *GBLP* was used as a reference transcript for normalization. Relevant primer sequences are listed in Table S7. The normalized fold change ± SD was calculated using the delta-delta Ct method (*70*). GraphPad Prism 8 was used to generate column charts and carry out statistical analyses.

**Table S7:**
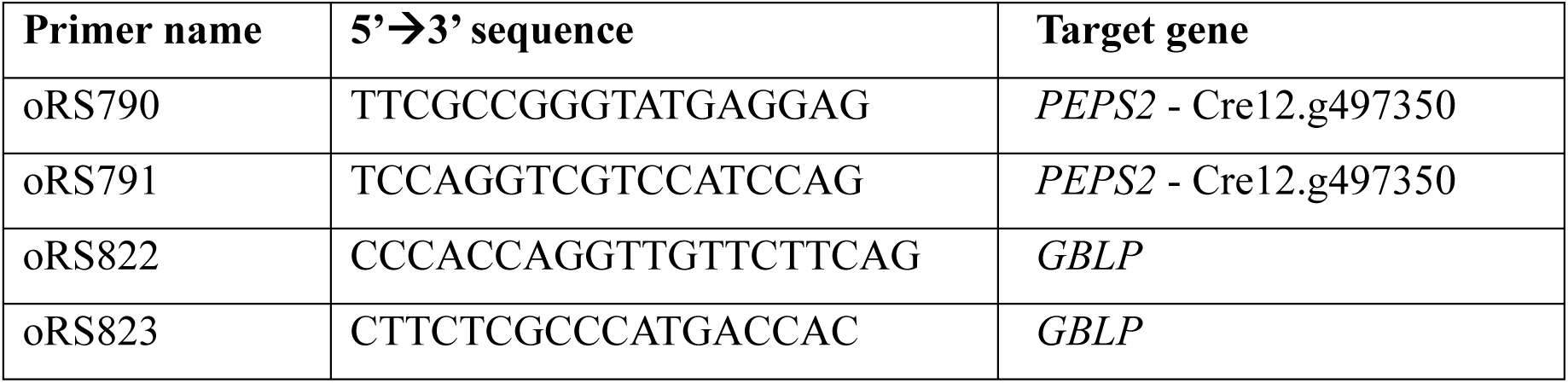
Primers used for RT-qPCR.

#### Northern Blot Analyses

Northern blotting was performed as described in (*31*). In brief, 2.5 to 15 μg of total RNA was separated on a 1.2% agarose/1.9% formaldehyde gel and transferred overnight onto a Hybond N+ nylon membrane (Amersham) in 10× SSC buffer (1.5 M NaCl, 0.15 M sodium citrate).The membrane was rinsed with MilliQ water, and the RNA was UV cross-linked using a Stratalinker cross-linking oven (Stratalinker) (set to autocross-linking 1200). Prehybridization (≥2 h) and hybridization (12 to 60 h) of the membrane with each [³²P]-labelled DNA probe was performed at 65°C in modified Church’s hybridization solution (0.5 M phosphate buffer, pH 7.2, 7% SDS (w/v), 10 mM EDTA, and 1% BSA). After hybridization, the membranes were washed at least twice at 65°C for 10 min each in washing buffer (40 mM phosphate buffer, pH 7.2, 1% SDS (w/v), and 1 mM EDTA). Signals were visualized by autoradiography or phosphorimaging. Each membrane was then subjected to mild stripping (1 h at 60°C in 0.5% SDS) and rehybridized with another probe for a transcript with a different size. Probes were labeled with [α-³²P]dATP by random priming (*71*), using PCR products corresponding to the full-length gene or gene fragments as templates. Primer combinations and sequences used to generate probes are listed in Tables S8 and S9.

**Table S8:** Primer combinations used to generate probes for Northern Blotting.

| Gene of interest | Primers used for PCR |
| --- | --- |
| *psbD* | SR109/SR110 |
| *rps12* | SR199/SR203 |
| *rpoA* | SR214/SR215 |
| *tufa* | SR261/SR262 |
| *atpB* | atpB For New/ atpB REV2 |
| *16s rRNA* | oRS1645/oRS1646 |

**Table S9:**
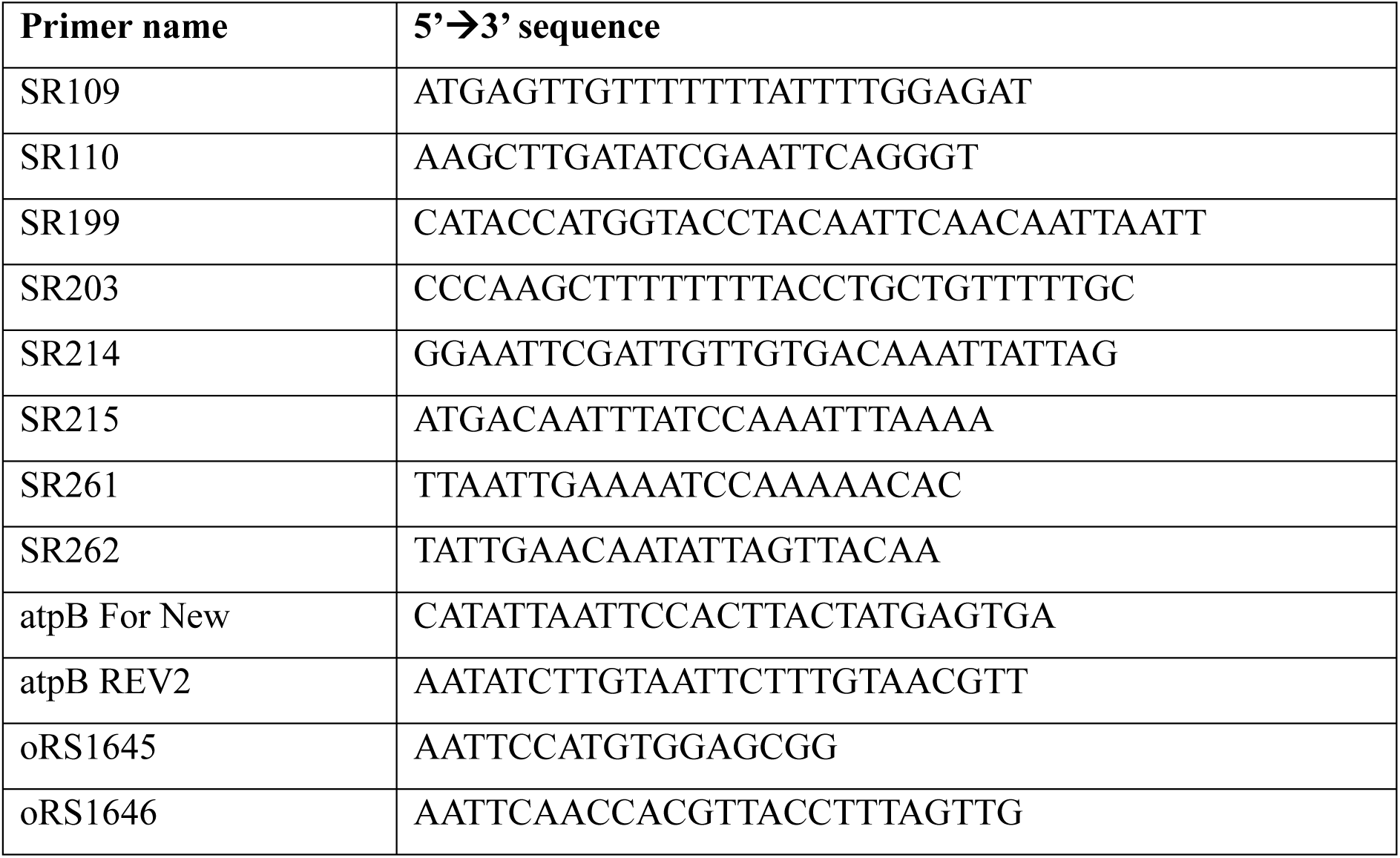
Primers used to generate probes for Northern Blotting.

#### Mass Spectrometry

##### STRAP

For the analysis of whole cell lysates from the wild type and *peps2* mutant (strains 959 and 960), the S-Trap protocol (Universal MS sample Prep Kit with micro columns, ProtiFi) was used according to the manufacturer’s protocol.

##### In-gel digest

For all other mass spectrometry analyses, proteins were separated by SDS–PAGE, visualized by Coomassie staining (see Total Protein Staining), and submitted for analysis as gel fragments. For the identification of CrPEP subunits following *rpoA*–FLAG immunoprecipitation, 15 discrete protein bands were excised and analyzed separately. For all other samples, the gel was run only briefly to allow protein entry and separation of any residual FLAG peptide used for elution, after which the entire lane was excised and processed for analysis.

Resultant Coomassie-stained gel fragments were cut into 2-3 mm pieces, transferred to 0.6 ml tubes and incubated with different solutions by shaking for 10 min at room temperature followed by removal of the supernatant as follows: Gel pieces were washed with 200 µl 100 mM ammonium bicarbonate (ABC), destained by 2 repeated rounds of shrinking in 200 µl 50% Acetonitrile (CAN) in 50mM ABC and reswelling in 200 µl 100 mM ABC. Gel pieces were shrunk with 100 µl ACN before being reduced with 100 µl 10mM Dithiothreitol (DTT) in 100 mM ABC by incubation at 56°C for 30 min. After another shrinking step in 100% ACN they were alkylated with 100 µl of 25 mM Iodoacetamide (IAA) in 100 mM ABC or 28 mM S-Methyl methanethiosulfonate (MMTS) by incubation at RT for 30 min in the dark.

Wash steps were repeated as described for destaining and gel pieces were shortly dried by vacuum centrifugation after the final shrinking step. Gel pieces were then soaked in 12.5 ng µl^-1^ Trypsin in ABC for 5 min at 4°C. Excess solution was removed and ABC was added to cover the pieces and samples were kept overnight at 37°C. The supernatant containing tryptic peptides was transferred to a fresh tube and gel pieces were extracted by addition of 20 µl 5% formic acid and sonication for 10 min in a cooled ultrasonic bath. This step was performed twice. All supernatants were unified.

Gel digests were desalted using an Oasis HLB 96-well µElution Plate (with 2 mg Sorbent per Well, Waters) according to the manufacturer’s description. A similar aliquot of each digest was analyzed by LC-MS/MS.

##### nanoLC-MS/MS Analysis

The nano HPLC system (Vanquish *Neo* UHPLC-System, Thermo Scientific) was coupled to an Orbitrap Exploris 480 mass spectrometer equipped with a FAIMS pro interface or to an Orbitrap Astral mass spectrometer, both equipped with a Nanospray Flex ion source (all parts Thermo Scientific). Peptides were loaded onto a trap column (PepMap Acclaim C18, 5 mm × 300 μm ID, 5 μm particle size, 100 Å pore size, Thermo Scientific) at a flow rate of 25 μl min^-1^ using 0.1% TFA as mobile phase.

After 5 min, the trap column was switched in line with the analytical column (PepMap Acclaim C18, 500 mm × 75 μm ID, 2 μm particles, 100 Å, Thermo Scientific operated at 30°C or Aurora Ultimate C18 25cm × 75 μm ID, 1.7 μm particles, 120 Å, with integrated emitter, Ionopticks operated at 50°C). Peptides were eluted using a flow rate of 230 nl min^-1,^ starting with the mobile phases 98% A (0.1% formic acid in water) and 2% B (80% acetonitrile, 0.1% formic acid) and linearly increasing to 35% B over the next 60 or 120 min respectively, followed by an increase to 95% B in 1.7 min, a 4-min hold at 95% B, and re-equilibration with 2% B for three column volumes (equilibration factor of 3.0).

The Orbitrap Exploris 480 mass spectrometer was operated in data-dependent mode, performing a full scan (m/z range 350-1200, resolution 60,000, AGC target 3,000,000) at 3 different compensation voltages (CV -45, -60, −75), followed each by MS/MS scans of the most abundant ions for a cycle time of 0.9 seconds per CV. MS/MS spectra were acquired using HCD collision energy of 30, isolation width of 1.2 m/z, orbitrap resolution of 30,000, AGC target 2,000,000, minimum intensity of 25,000 and maximum injection time of 100 ms. Precursor ions selected for fragmentation (include charge state 2-6) were excluded for 45 s. The monoisotopic precursor selection filter and exclude isotopes feature were enabled.

The Orbitrap Astral was operated in data-independent mode, performing a full scan in the Orbitrap every 0.6 s (m/z range 380–980 m/z; resolution 240,000; AGC target 1,000,000, maximum injection time 5 ms). MS/MS spectra were acquired in the Astral analyser by isolating 5 Da windows across 380–980 m/z (resulting in 119 scan events per cycle) and fragmenting precursor ions with HCD collision energy of 25% with a maximum injection time of 3 ms or until an AGC target of 30,000 was reached. Fragment ions ranging from 150–2000 m/z were acquired.

##### Data Processing

###### Exploris

For peptide identification, the RAW-files were loaded into Proteome Discoverer (version 3.2.0.450, Thermo Scientific). All MS/MS spectra were searched using MSAmanda version 3.2.22.93 (*72*). The peptide and fragment mass tolerance was set to ±10 ppm, the maximum number of missed cleavages was set to 2, using tryptic enzymatic specificity without proline restriction. The RAW-files were searched against the *Chlamydomonas reinhardtii* v6.1 Phytozome database (30,357 sequences; 24,276,980 residues, https://phytozome-next.jgi.doe.gov/) (*73*), supplemented with common contaminants using the following modifications: Carbamidomethylation or beta-methylthiolation of cysteine was set as fixed modification and oxidation of methionine, deamidation of asparagine and glutamine, glutamine to pyro-glutamate conversion at peptide N-terminal glutamine and acetylation on the protein N-terminus were set as variable modifications. The localization of the post-translational modification sites within the peptides was performed with the tool ptmRS, based on the tool phosphoRS (*74*).

The result was filtered to 1% FDR on PSM and protein level using the Percolator algorithm (*75*) integrated in Proteome Discoverer. Additionally, an Amanda score cut-off of at least 150 was applied. Proteins were filtered to be identified by a minimum of 2 PSMs in at least 1 sample.

Protein areas were computed in IMP-apQuant (*76*) by summing up unique and razor peptides. Resulting protein areas were normalized using iBAQ (*77*) and for comparisons between eluates, data were normalised on the amount of the bait protein (RpoA-FLAG). Match-between-runs (MBR) was applied for peptides with high confident peak area that were identified by MS/MS spectra in at least one run. Results were filtered for only the proteins of interest.

###### Astral

Astral DIA data were analyzed in Spectronaut 20.1. (Biognosys). Trypsin/P was specified as a proteolytic enzyme and up to 2 missed cleavages were allowed in the Pulsar directDIA+ search. Dynamic mass tolerance was applied for calibration and main search. The search was performed against the same Chlamydomonas and contaminant databases as described for the Exploris data. Beta-methylthiolation of cysteine was searched as fixed modification, whereas oxidation of methionine and acetylation at protein N-termini were defined as variable modifications. Peptides with a length between 7 and 52 amino acids were considered and results were filtered using Spectronaut default filtering criteria (Precursor Qvalue<0.01, Precursor PEP<0.2, Protein Qvalue <0.01 per Experiment and <0.05 per Run, Protein PEP<0.75). Quantification was performed as specified in Biognosys BGS Factory Default settings, grouping peptides by stripped sequence and performing protein inference using IDPicker. Cross-run normalization in Spectronaut was deactivated due to subsequent mode normalization.

Spectronaut results were exported using Pivot reports on the protein and peptide level and converted to Microsoft Excel files using our in-house software MS2Go (https://ms.imp.ac.at/?action=ms2go). For DIA data MS2Go utilizes the python library msReport (developed at the Max Perutz Labs Proteomics Facility) for data processing. Abundances were normalized by the mode of protein ratios in msReport and missing values were imputed with values obtained from a log-normal distribution with a mean of 100. To compensate for different protein lengths, protein quantification was then normalized using iBAQ (*77*). Statistical significance of differentially expressed proteins was determined using limma (*78*).

#### RNA extension assay

For the RNA extension assay, we adapted the protocol described in (*79*). Oligonucleotides as described in Table S10 were ordered from Integrated DNA Technologies at HPLC purity and diluted to stock concentrations of 20 µM in nuclease-free water. The RNA-tDNA hybrid was first prepared by mixing tDNA and 5’ FAM-labelled RNA to 2.8 µM final concentration in 1X

*E. coli* RNA Pol Buffer (New England BioLabs) and incubating for 5 min at 95°C, followed by gradual cooling to room temperature over 1 hour. 4 µl of the annealed scaffold was divided into individual PCR tubes, and to each tube 1 µl of RNase OUT™ Recombinant Ribonuclease Inhibitor (Invitrogen) was added to prevent RNA degradation. For reactions containing CrPEP, 3.5 μl of purified complex was added, unless otherwise stated. As a positive control, 1 μl of *E. coli* core RNA polymerase (New England BioLabs) was added to parallel reactions.

Reactions were then incubated for 10 min at 37°C prior to the addition of ntDNA to a final concentration of 750 nM, followed by a further 10 min incubation at 37°C. Each reaction was then adjusted to a final volume of 15 µl by adding 3 µl 5X *E. coli* RNA Pol buffer, 1.5 µl NTPs (5 mM stock), and nuclease-free water. The mixtures were then incubated for 30 min at 37°C, and transcription was terminated by addition of 4 µg of Proteinase K (ThermoFisher) followed by 20 min incubation at 30°C. Prior to loading, samples were denatured via incubation at 95°C for 5 min. Transcription products were separated via denaturing polyacrylamide gel electrophoresis using an 8 M urea 20% acrylamide/bisacrylamide gel (19:1 ratio) in 1X TBE buffer. Electrophoresis was performed with an initial pre-run at 260 V for 15 min, followed by 15 min at 250 V, and a final run at 300 V for 90 min. Products were visualized by detecting the 5’ FAM-labelled RNA using an Azure Sapphire biomolecular imager.

**Table S10:**
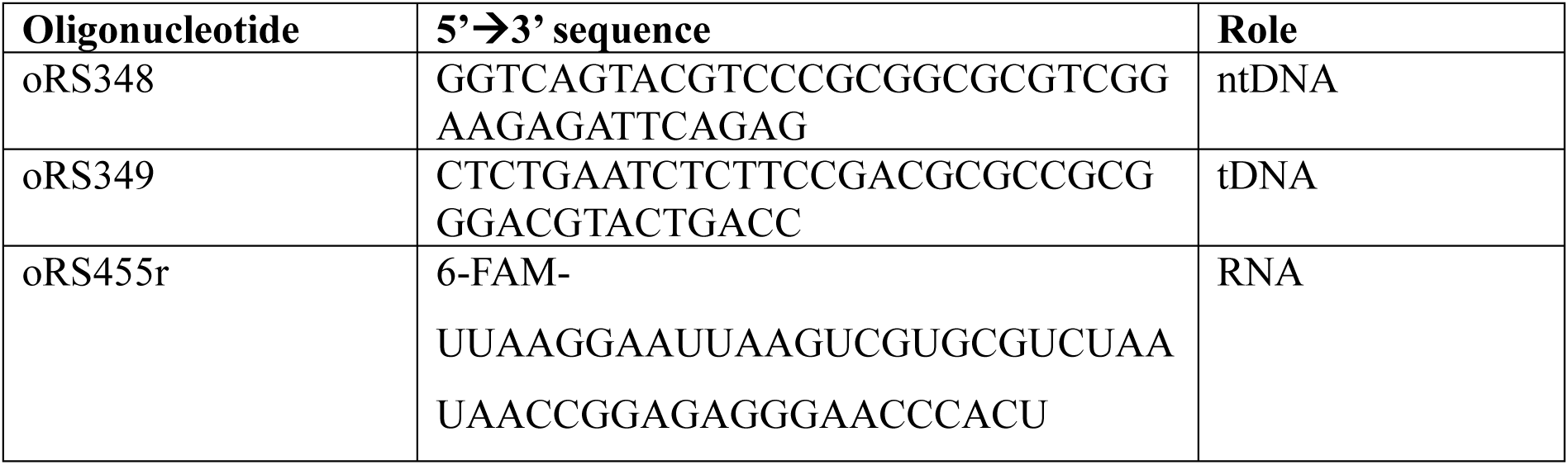
Oligonucleotide sequences used for the RNA extension assay.

#### Sucrose Gradient Analysis

For sucrose gradient analysis of cell lysates from the WT and *peps2* mutant (strains 959 and 960), cultures were grown in TAP medium to a density of 2–8 × 10⁶ cells ml⁻¹. Cells were harvested at 4 °C by centrifugation at 3000 × g for 10 min, and the pellet resuspended in an equal volume of 2× lysis buffer without glycerol (100 mM HEPES, 100 mM KOAc, 4 mM Mg(OAc)₂·4H₂O, 2 mM CaCl₂; pH adjusted to 6.8 with KOH; freshly supplemented with EDTA-free protease inhibitor cocktail (Roche), two tablets per 25 ml buffer).

Cell lysis was performed by freeze–thaw cycling: 2 ml of resuspended cells were transferred to an Eppendorf tube, snap-frozen in liquid nitrogen, thawed completely at room temperature, and this cycle was repeated five times. Lysates were clarified by centrifugation at maximum speed in a tabletop centrifuge for 30 min at 4 °C, and the supernatant collected as input.

Linear sucrose density gradients (15–45% sucrose in 1× lysis buffer plus protease inhibitor) were prepared in 11 × 60 mm tubes suitable for an SWTi-60 rotor (Beckman Instruments). A total of 200 µl clarified lysate was layered onto each gradient, which was centrifuged at 164,000 x g for 16 h at 4 °C. Following centrifugation, gradients were fractionated into 18 fractions. Proteins were recovered by NaDOC/TCA precipitation: fractions were adjusted to 1 ml with Milli-Q water, mixed with 10 µl of 10% sodium deoxycholate, vortexed, and precipitated with 50% trichloroacetic acid (TCA), followed by vortexing for 2 min and overnight incubation at –20°C. Protein pellets were collected by centrifugation at maximum speed for 20 min at 4°C, washed twice with 1 ml 100% acetone, and air-dried at room temperature for 2 h. The final protein pellets were resuspended in 50 µl 2× SDS lysis buffer (100 mM Tris, pH 8.0; 600 mM NaCl; 20 mM EDTA; 4% SDS), freshly supplemented with EDTA-free protease inhibitor cocktail (Roche). 10 µl of each fraction, along with 4 µl of clarified lysate input, were separated by SDS–PAGE on NuPAGE 4–12% Bis-Tris midi gels (Invitrogen). The distribution of CrPEP components across gradient fractions was analyzed by immunoblotting.

#### Immunoblot analyses

Proteins were transferred to nitrocellulose membranes (Amersham Protran 0.2 µm NC; Cytiva) by wet transfer at 70 V for 90 min at 4 °C. Membranes were blocked in PBS-T with 5% instant non-fat dry milk for 1h at room temperature and incubated overnight at 4°C with primary antibodies diluted in the same blocking buffer. The following antibodies were used in this study: for sucrose gradient analysis: polyclonal rabbit anti-RpoA (1:10,000) (*31*) and polyclonal rabbit anti-PEPS1 (raised against recombinant PEPS1Trunc protein, generated by Eurogentec; 1:20,000). For screening of *rpoA*-FLAG transformants and PEPS3/9/12-Venus-3xFLAG strains, monoclonal mouse anti-FLAG M2 (Sigma-Aldrich, 1:5000). As loading control: monoclonal mouse anti-α-Tubulin (Sigma-Aldrich, 1:10,000).

Membranes were washed three times in PBS-T with 5% milk (10 min each, room temperature), incubated with either HRP-conjugated anti-rabbit secondary antibody (Promega; 1:1,000) or HRP-conjugated anti-mouse secondary antibody (Promega, 1:10,000) for 1 h at room temperature, and washed again three times with PBS-T. Signal detection was carried out using luminol-based enhanced chemiluminescence (ECL). For PEPS1 detection, the SuperSignal West Dura Extended Duration Substrate kit (Thermo Fisher) was used, while RpoA, FLAG, and α-Tubulin required the more sensitive SuperSignal West Pico PLUS kit (ThermoFisher). Chemiluminescent signals were imaged using the iBright FL 1500 system (ThermoFisher).

#### Cloning and Purification of Recombinant Proteins

##### PEPS1Trunc

For PEPS1Trunc, an *E. coli* codon optimised sequence corresponding to residues 47-570 of the full-length PEPS1 protein (excluding the cTP as predicted by TargetP2.0 (*80*)) was synthesized by Twist Bioscience and cloned into the pET-28a(+) *E. coli* expression vector (pRAM457). These plasmids were then transformed into chemically competent BL21(DE3) *Escherichia coli* cells via heat shock. Transformed cells were grown in LB medium containing 50 µg ml^-1^ kanamycin at 37°C to an OD_600_ of 0.7-0.8, induced with 0.5 mM IPTG, and incubated overnight at 25°C. Cells were harvested by centrifugation at 4000 x g for 30 min at 4°C, and the pellet snap frozen and stored at −70°C until purification.

Cell pellet from 2 L of culture was resuspended in 100 ml of *E. coli* Lysis Buffer (1 M NaCl, 20 mM KCl, 60 mM Na_2_HPO_4_, 10 mM KH_2_PO_4_) supplemented with benzonase (2 µl per 1 ml of buffer), 1 mM MgCl2, and 4 tablets of EDTA-Free Protease Inhibitor Cocktail, and lysed by sonication using a 19 mm probe (VCX750 VibraCell Ultrasonic Processor, Sonics & Materials, Inc.) for a total of 5 min (1 s pulse on, 2 s pulse off) at 60% amplitude. To precipitate nucleic acids and ensure high protein purity, the cell lysate was incubated with 5% (w/v) polyethyleneimine (PEI) on ice for 20 min, then centrifuged at 40,000 x g for 30 min at 4°C. The supernatant was collected, and proteins were precipitated by slowly adding ammonium sulfate (8.7 g per 20 mL lysate) while stirring in the cold room for 30 min. The ammonium sulfate precipitate was collected by centrifugation at 40,000 x g for 30 min at 4°C and resuspended in 90 ml of IMAC A buffer (2X PBS, 20 mM imidazole) for further purification. Protein purification was carried out using a three-step approach. Clarified lysate was first loaded onto a 5 ml HisTrap FF column (Cytiva) and eluted using 25 ml step gradients of 50-500 mM imidazole. Fractions were analysed via SDS-PAGE, pooled based on purity, and the pooled eluate dialyzed overnight in QS Buffer (50 mM Tris 7.5, 50 mM NaCl) at 4°C with constant mixing. Following dialysis, the sample was loaded onto a 1 ml HiTrap Q FF column and eluted using a gradient from 50 to 500 mM NaCl. Peak fractions were concentrated using a Vivaspin 20 30 kDa MWCO concentrator (Sartorius) and further purified by size exclusion chromatography on a Superdex 200 16/60 column (Cytiva) equilibrated with SEC buffer (1× PBS, 5% glycerol). Purity of resultant fractions was assessed by SDS-PAGE. As the protein eluted at three different peaks, the lowest molecular weight peak (likely representing monomeric PEPS1Trunc), was pooled and concentrated to a final concentration of 2.64 mg ml^-1^ for use in downstream experiments. 25 µl aliquots of purified protein were prepared, snap-frozen in liquid nitrogen, and stored at –70°C to prevent repeated freeze–thaw cycles.

**Table S11:**
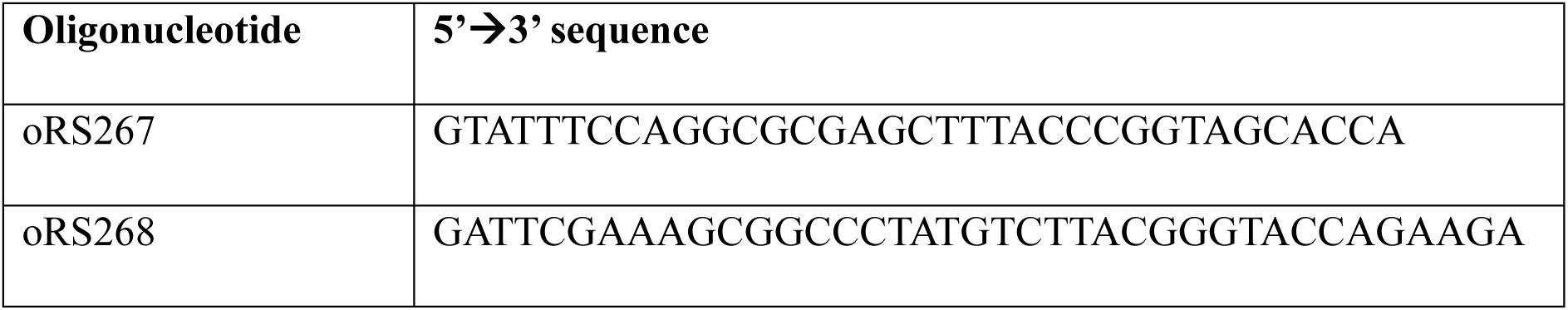
Primers used for insertion of *RPOD1* fragment into pFBD-HS destination plasmid.

##### Cr σ factor (*RPOD1*)

For expression of Cr σ factor (*RPOD1*; Cre03.g194950), an insect cell expression system was chosen to avoid potential toxicity in bacterial hosts. To generate the expression construct, a dsDNA fragment containing the full-length RPOD1 coding sequence was synthesized by Twist Bioscience. This fragment was amplified with primers oRS267 and oRS268 (Table S11) to introduce sequences overlapping with the destination plasmid. The expression vector used was the pFBD-HS plasmid (gift from the Leonard Lab), containing dual N-terminal DecaHis and Strep tags. Following digestion with BssHII and NotI, the vector and insert DNA fragments were assembled using In-Fusion reagents (Takara) according to the manufacturer’s instructions, yielding plasmid pRAM366.

The pRAM366 construct was then used to generate recombinant baculovirus using the FastBac system. Sf9 cells were infected with high-titer virus, harvested 3-4 days post-infection by centrifugation at 4000 x g for 30 min at 4°C, and stored at −70°C until purification.

Purification was carried out using a two-step approach. Two litre insect cell pellets were first resuspended in 100 ml IMAC Buffer A (50 mM HEPES 7.5, 500 mM NaCl, 1 mM TCEP, 20 mM imidazole) supplemented with benzonase (2 µl per 1 ml buffer), 2 mM MgCl, 0.2% (v/v) NP-40 Alternative and 4 EDTA-Free Protease Inhibitor Cocktail tablets with gentle mixing. Cells were cleared by centrifugation at 18000 x g for 30 min at 4°C, and the clarified lysate loaded onto a 5 ml HisTrap FF column (Cytiva) and eluted with a gradient from 50-500 mM imidazole. Elution fractions were analysed by SDS-PAGE, pooled based on purity, and dialysed overnight into SEC buffer (50 mM HEPES 7.5, 500 mM NaCl, 1 mM TCEP). The sample was then concentrated to 1 ml total volume (Vivaspin 20, 50 kDa MWCO, Sartorius), and loaded onto a Superose 6 Increase 10/300 column (Cytiva). Following size exclusion chromatography, eluent was collected in 48 sequential 500 μl fractions using a fraction collector. Elution fractions B11-C12 from SEC were analysed via SDS-PAGE, and the peak corresponding to the correct molecular weight product (fractions B10-C2), pooled and concentrated to 4 mg ml^-1^ for use in downstream experiments. 12 µl aliquots of purified protein were prepared, snap-frozen in liquid nitrogen, and stored at –70°C to prevent repeated freeze–thaw cycles.

##### *RPOD1* gene fragment sequence

GCTTTACCCGGTAGCACCATGAACTTAACCACACGTTGTTCCACAACGCCTCGTTCCGCGGTTGTTGCCCGCGCAGTAGCCGCCCCGACACGTCCCACCACAAAATCCGCGGTGCCAGAACTTTTAGACTCCCGTCCAGGAGAACGTAACCTTAATTTCATGGAGTACGCGCAAGCAACCCAAATGTTAGACCGCCTGAAGGGCCAAGCGTCGGATTTAGAGTTACTGTTAGATCAGCTGAACGCTTTGGAGGCGAGCCTGGATGAGAGTGTATTGGCCCCTCCTACAGTGGATGACCCAAAGGAACGTGCAGCGCGTCAGGCTCGCCGTGCGGCAAAACGTGCAGAGCGCCGCGCCCAGGCTACTTCGGCGACCGTCGCTGCTGCCGCCGGCCCGGCAATGTCGGCGGTTGTGTCTCACTCCACGCCCACCAAGGCGGCAGCAGCTCCAGCTACTTCCACCGCTAGCAGCTCATCATCAGACTCGGGACTTCTTGACCTTGTTTCATTTGTCGGGGGATTCGATACTCGTCCGATCCCGGCGACGACAAGTGCCCCACCGGCAGGGGCGAGCTCCAGTGATGTGCAACACCTGGAAGACCTGTTTAAATTAAGCGTAGGGGAACCAGATATTCCCCGTGCCTCAGCGTCGGCTGCCCCTGCTGTTTTACGCCCTCGTAAATTAACACCCAAGAAGCCCAGCGCTGCTCCGTCAGCAGCAGTCACAGCTGCACCCTCGCCCGCGCCCACATTACCCAGTACCCCATCTACTAGCGCGCGCATCGCCCCTGCTCCTGGGTCCCTTGCAGATGAATTGGAACGTTTGCTTGGTCCAACGACCTCTCGCGAGGCTGCCGAATCCGAGGATGAAGATTCCTTTGCGGGGCCGTCAGAGGATGATTTACTGGCATTGGAGCAAGAAGTATCACGTAAAAGCTCTCGCTTGCCTGTATTAGACGAAGAGGACGAAGAGGATGAACAGCAGCAATTAGAAGACAACGAAGAGGACGCAGTAGCCGGACCAGGCAGTTTGGAGGCGAGCGCGATGGCAACGCGCACGAGCAGCCAACTTTCCATTATGCAGACGGGGCCGTCGCTTCTTTCATTGGTGCCAGCTAGCGCGGCACCAGGTCGTTCCGCTAAAGCC CGTGCGAGCCGCCGCGCGGCGCGTAATGGGCACGCTTCGGGTCGCTTGGGTGGGGCCACTGCAAATGCGGCCGGGCGCGGTAAAGTTGGCAGCAAAGATGGCACAATGAATTTCTTGGGAAAAGTCGAATCTTTATCGACATTAGATGTCGAAAAGGAGCGCGAGGTAACCGCTGTCTGCCGTGACTTCTTGTTCCTGGAGAAAGTTAAGCGTCAATGCGAGAAAACACTGCATCGTCCAGCCACAAGCGAGGAGATTGCTGCTGCCGTAGCAATGGATGTTGAGAGTCTTAAGCTGCGCTATGACGCTGGGCTGAAAGCCAAGGAACTTCTTCTTAAGTCCAATTATAAGCTGGTAATGACGGTATGTAAGAGCTTCGTCGGCAAAGGGCCCCACATTCAAGATTTGGTATCGGAAGGTGTCAAGGGATTACTTAAGGGCGTTGAGAAGTACGACGCTACTAAGGGATTCCGCTTCGGAACATATGCACATTGGTGGATCCGCCAAGCGGTGAGTCGTAGTCTTGCCGAAACGGGACGCGCCGTTCGCTTGCCGATGCACATGATCGAGCAATTGACTCGTTTAAAAAACTTAAGTGCAAAGCTGCAAACGCAACTGGCCCGCGAGCCTACCTTACCAGAACTGGCAAAAGCGGCGGGGTTGCCGGTTACCCGTGTTCAGATGCTGATGGAAACCGCTCGTTCAGCCGCTTCCCTTGACACACCAATCGGAGGTAACGAACTGGGACCCACCGTTAAAGACAGCGTCGAGGACGAGCGCGAAGCAGCGGATGAAGAATTCGGATCTGACTCTCTGCGCAACGATATGGAAGCAATGTTACTGGAGCTTCCGGAACGTGAGGCGCGTGTAGTCCGCCTGCGTTTCGGCTTAGATGATGGTAAGGAGTGGACACTGGAAGAAATCGGAGAAGCCCTGAATGTCACCCGCGAGCGTATTCGCCAGATCGAGGCGAAAGCACTGCGCAAGCTTCGTGTGAAAACGATCGACGTTTCGGGCAAATTAATGGAATACGGAGAAAACTTAGAAATGTTGATGGATGGCAGCCGTGAGATGGCCGCACGTACATCTTCTGGTACCCGTAAGACATAG

#### Electrophoretic Mobility Shift Assay

A 5’ FAM-labeled DNA probe corresponding to the Chlamydomonas chloroplast 16s rRNA promoter sequence was generated by PCR from wild-type genomic DNA using primers oRS1000 and oRS1002 (Table S12). The resulting 147 bp probe was verified by electrophoresis in a 1% agarose gel, purified and quantified using a Nanodrop ND-100 Spectrophotometer (ThermoFisher).

Binding reactions were assembled on ice in a total volume of 15 µl in binding buffer (20 mM Tris, pH 8.0; 50 mM NaCl; 1 mM DTT; 10% glycerol) containing 10 nM DNA probe and varying concentrations of the relevant protein, and subsequently incubated at 30°C for 20 min. After incubation, 3.75 µl of Novex Hi-Density 5× TBE Sample Buffer (Invitrogen) was added, and samples were resolved on 6% DNA Retardation Gels (Invitrogen) by electrophoresis at 80 V for 3 h at 4°C. Fluorescent signals corresponding to free and protein-bound FAM-labelled DNA were visualised with an Azure Sapphire biomolecular imager.

**Table S12:**
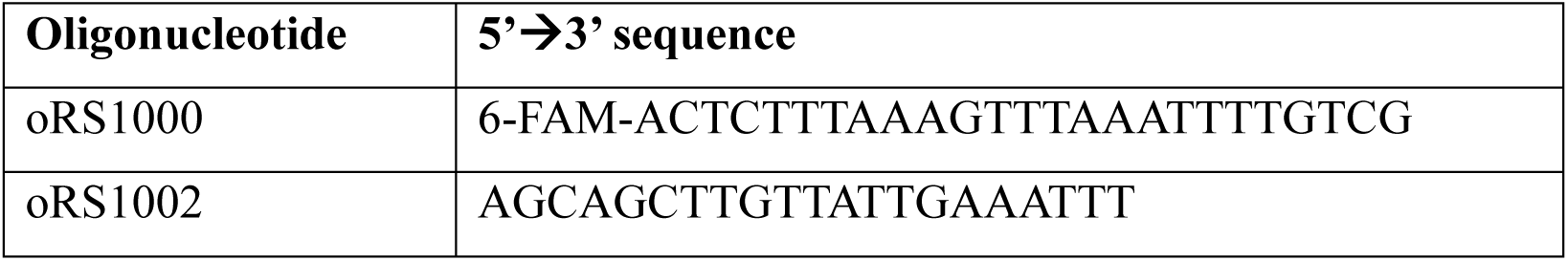
Primers used for the generation of the 16s rRNA promoter EMSA probe.

#### Phylogenetic analyses and evolutionary inference

We searched for homologs of each of the PEPS proteins identified from the *rpoA*-FLAG IP and subsequent mass spectrometry analysis. Corresponding protein sequences were queried against a dataset of predicted proteomes of a phylodivese set of Chloroplastida species with sequenced genomes using BLASTp with an e-value threshold <1e^-6^. The predicted proteomes of the following species were considered: chlorophyte algae (*Chlamydomonas reinhardtii, Chlorella variabilis, Coccomyxa subellipsoidea, Micromonas pusilla, Ostreococcus lucimarinus, Ulva mutabilis*), streptophyte algae (*Chara braunii, Chlorokybus melkonianii, Klebsormidium nitens, Mesostigma viride, Mesotaenium endlicherianum, Penium margaritaceum, Spirogloea muscicola*), and embryophytes (*Anthoceros agrestis, Arabidopsis thaliana, Gnetum montanum, Physcomitrium patens, Amborella trichopoda, Isoetes taiwanensis, Marchantia polymorpha, Oryza sativa, Selaginella moellendorffii*). All significant hits were aligned together with the query sequences using mafft (FFT-NS-2) and maximum likelihood phylogenies were inferred using IQ-TREE under BIC-selected best-fit substitution models and 1000 replicates of non-parametric bootstrapping. Resultant phylogenetic trees were visualized and annotated using iTOL (Data S6). Homologues from *A. thaliana* were annotated based on their TAIR annotations. Orthology of the Chlamydomonas PEPSs to land plant PAPs was assessed using a curated list of *A. thaliana* PAPs from (*26*).

Vice versa, homology of the identified PEPSs was determined and plotted as in Figure 6b. The phylogenetic distribution of the 12 PEPSs was then mapped onto a cladogram of the green lineage, and their evolutionary history was inferred.

