## Supplementary material for "The structure of a 2-MDa chloroplast RNA polymerase reveals unexpected evolutionary complexity": Data S2

### Full wwPDB EM Validation Report (i)

Apr 21, 2026 – 11:38 am BST

| | | |
|---|---|---|
| PDB ID | : | 28RF / pdb_000028rf |
| EMDB ID | : | EMD-56766 |
| Title | : | Cryo-EM single particle structure of the Plastid-Encoded RNA Polymerase (PEP)from Chlamydomonas reinhardtii. |
| Deposited on | : | 2026-02-15 |
| Resolution | : | 2.65 Å(reported) |

**This wwPDB validation report is for manuscript review**

This is a Full wwPDB EM Validation Report.

This report is produced by the wwPDB biocuration pipeline after annotation of the structure.

We welcome your comments at **
A user guide is available at
https://www.wwpdb.org/validation/2017/EMValidationReportHelp
with specific help available everywhere you see the (i) symbol.

The types of validation reports are described at
http://www.wwpdb.org/validation/2017/FAQs#types.

The following versions of software and data (see references (i)) were used in the production of this report:

| | | |
|---|---|---|
| EMDB validation analysis | : | 0.0.1.dev132 |
| MolProbity | : | 4-5-2 with Phenix2.0 |
| Percentile statistics | : | 20250101.v01 (using entries in the PDB archive January 1st 2025) |
| EM percentile statistics | : | 202505.v01 (Using data in the EMDB archive up until May 2025) |
| MapQ | : | 1.9.13 |
| Ideal geometry (proteins) | : | Engh & Huber (2001) |
| Ideal geometry (DNA, RNA) | : | Parkinson et al. (1996) |
| Validation Pipeline (wwPDB-VP) | : | 2.49 |

For Manuscript Review

### 1 Overall quality at a glance (i)

The following experimental techniques were used to determine the structure:
*ELECTRON MICROSCOPY*

The reported resolution of this entry is 2.65 Å.

Percentile scores (ranging between 0-100) for global validation metrics of the entry are shown in the following graphic. The table shows the number of entries on which the scores are based.

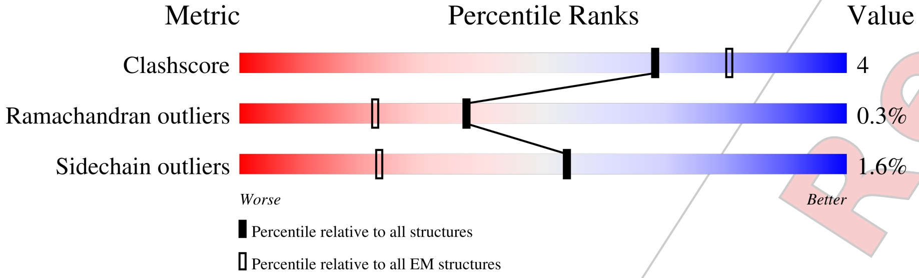


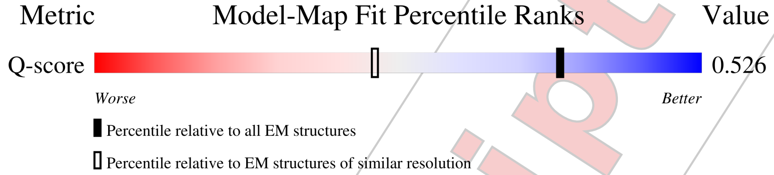


| Metric | Whole archive (#Entries) | EM structures (#Entries) | Similar EM resolution (#Entries, resolution range(Å)) |
|---|---|---|---|
| Clashscore | 229148 | 23984 | - |
| Ramachandran outliers | 224038 | 23583 | - |
| Sidechain outliers | 223484 | 23102 | - |
| Q-score | - | 25397 | 9050 ( 2.15 - 3.15 ) |

The table below summarises the geometric issues observed across the polymeric chains and their fit to the map. The red, orange, yellow and green segments of the bar indicate the fraction of residues that contain outliers for >=3, 2, 1 and 0 types of geometric quality criteria respectively. A grey segment represents the fraction of residues that are not modelled. The numeric value for each fraction is indicated below the corresponding segment, with a dot representing fractions <=5% The upper red bar (where present) indicates the fraction of residues that have poor fit to the EM map (all-atom inclusion < 40%). The numeric value is given above the bar.

| Mol | Chain | Length | Quality of chain |
|---|---|---|---|
| 1 | A | 737 | 52% 5% 43% |
| 1 | Z | 737 | • 56% 5% 39% |
| 2 | B | 822 | 70% 5% 25% |

*Continued on next page...*

For Manuscript Review

*Continued from previous page...*

| Mol | Chain | Length | Quality of chain |
|---|---|---|---|
| 3 | C | 626 | 60% 8% 32% |
| 4 | D | 3120 | 51% • 45% |
| 5 | E | 1932 | • 27% 8% 64% |
| 6 | F | 648 | 65% 5% 30% |
| 7 | G | 834 | 74% 6% 20% |
| 8 | H | 609 | 66% 6% 28% |
| 8 | I | 609 | 47% 6% 47% |
| 9 | J | 343 | 35% • 61% |
| 10 | K | 2788 | • 58% • 38% |
| 11 | M | 1801 | 7% 18% • 79% |
| 12 | N | 803 | 11% 58% 19% • 22% |
| 13 | Q | 414 | 18% 47% 7% 46% |
| 14 | R | 356 | 81% • 15% |
| 15 | S | 359 | 73% 6% 20% |
| 16 | T | 381 | 19% 6% • 73% |
| 17 | X | 410 | 81% 5% 14% |
| 17 | Y | 410 | 85% 5% 10% |

For Manuscript Review

### 2 Entry composition (i)

There are 17 unique types of molecules in this entry. The entry contains 83535 atoms, of which 0 are hydrogens and 0 are deuteriums.

In the tables below, the AltConf column contains the number of residues with at least one atom in alternate conformation and the Trace column contains the number of residues modelled with at most 2 atoms.

- Molecule 1 is a protein called DNA-directed RNA polymerase.

| Mol | Chain | Residues | Atoms | | | | | AltConf | Trace |
|---|---|---|---|---|---|---|---|---|---|
| | | | Total | C | N | O | S | | |
| 1 | A | 421 | 3368 | 2181 | 558 | 615 | 14 | 0 | 0 |
| 1 | Z | 453 | 3590 | 2321 | 594 | 664 | 11 | 0 | 0 |

- Molecule 2 is a protein called DNA-directed RNA polymerase subunit beta N-terminal section.

| Mol | Chain | Residues | Atoms | | | | | AltConf | Trace |
|---|---|---|---|---|---|---|---|---|---|
| | | | Total | C | N | O | S | | |
| 2 | B | 617 | 4970 | 3213 | 849 | 890 | 18 | 0 | 0 |

- Molecule 3 is a protein called DNA-directed RNA polymerase subunit beta C-terminal section.

| Mol | Chain | Residues | Atoms | | | | | AltConf | Trace |
|---|---|---|---|---|---|---|---|---|---|
| | | | Total | C | N | O | S | | |
| 3 | C | 424 | 3368 | 2156 | 572 | 625 | 15 | 0 | 0 |

- Molecule 4 is a protein called DNA-directed RNA polymerase subunit beta".

| Mol | Chain | Residues | Atoms | | | | | AltConf | Trace |
|---|---|---|---|---|---|---|---|---|---|
| | | | Total | C | N | O | S | | |
| 4 | D | 1725 | 14136 | 9263 | 2372 | 2469 | 32 | 0 | 0 |

- Molecule 5 is a protein called DNA-directed RNA polymerase subunit.

| Mol | Chain | Residues | Atoms | | | | | AltConf | Trace |
|---|---|---|---|---|---|---|---|---|---|
| | | | Total | C | N | O | S | | |
| 5 | E | 696 | 5698 | 3684 | 1001 | 992 | 21 | 0 | 0 |

- Molecule 6 is a protein called Glucose-6-phosphate 1-epimerase.

For Manuscript Review

| Mol | Chain | Residues | Atoms | | | | | AltConf | Trace |
|---|---|---|---|---|---|---|---|---|---|
| 6 | F | 452 | Total 3517 | C 2234 | N 596 | O 672 | S 15 | 0 | 0 |

- Molecule 7 is a protein called Rubisco LSMT substrate-binding domain-containing protein.

| Mol | Chain | Residues | Atoms | | | | | AltConf | Trace |
|---|---|---|---|---|---|---|---|---|---|
| 7 | G | 664 | Total 5136 | C 3246 | N 887 | O 980 | S 23 | 0 | 0 |

- Molecule 8 is a protein called UDP-3-O-acyl-N-acetylglucosamine deacetylase.

| Mol | Chain | Residues | Atoms | | | | | AltConf | Trace |
|---|---|---|---|---|---|---|---|---|---|
| 8 | H | 441 | Total 3419 | C 2148 | N 593 | O 665 | S 13 | 0 | 0 |
| 8 | I | 324 | Total 2506 | C 1590 | N 431 | O 474 | S 11 | 0 | 0 |

- Molecule 9 is a protein called Uncharacterized protein A0A2K3DRS4.

| Mol | Chain | Residues | Atoms | | | | | AltConf | Trace |
|---|---|---|---|---|---|---|---|---|---|
| 9 | J | 135 | Total 1105 | C 697 | N 189 | O 212 | S 7 | 0 | 0 |

- Molecule 10 is a protein called SAP domain-containing protein.

| Mol | Chain | Residues | Atoms | | | | | AltConf | Trace |
|---|---|---|---|---|---|---|---|---|---|
| 10 | K | 1726 | Total 12323 | C 7722 | N 2159 | O 2397 | S 45 | 0 | 0 |

- Molecule 11 is a protein called Uncharacterized protein A0A2K3D3G3.

| Mol | Chain | Residues | Atoms | | | | | AltConf | Trace |
|---|---|---|---|---|---|---|---|---|---|
| 11 | M | 375 | Total 2794 | C 1739 | N 485 | O 556 | S 14 | 0 | 0 |

- Molecule 12 is a protein called Mur ligase central domain-containing protein.

| Mol | Chain | Residues | Atoms | | | | | AltConf | Trace |
|---|---|---|---|---|---|---|---|---|---|
| 12 | N | 623 | Total 4733 | C 2987 | N 814 | O 921 | S 11 | 0 | 0 |

- Molecule 13 is a protein called Coenzyme Q-binding protein COQ10 START domain-containing

For Manuscript Review

protein.

| Mol | Chain | Residues | Atoms | | | | | AltConf | Trace |
|---|---|---|---|---|---|---|---|---|---|
| | | | Total | C | N | O | S | | |
| 13 | Q | 225 | 1730 | 1098 | 295 | 331 | 6 | 0 | 0 |

- Molecule 14 is a protein called S1 motif domain-containing protein.

| Mol | Chain | Residues | Atoms | | | | | AltConf | Trace |
|---|---|---|---|---|---|---|---|---|---|
| | | | Total | C | N | O | S | | |
| 14 | R | 301 | 2438 | 1534 | 401 | 492 | 11 | 0 | 0 |

- Molecule 15 is a protein called Uncharacterized protein.

| Mol | Chain | Residues | Atoms | | | | | AltConf | Trace |
|---|---|---|---|---|---|---|---|---|---|
| | | | Total | C | N | O | S | | |
| 15 | S | 287 | 2374 | 1467 | 414 | 484 | 9 | 0 | 0 |

- Molecule 16 is a protein called Uncharacterized protein.

| Mol | Chain | Residues | Atoms | | | | | AltConf | Trace |
|---|---|---|---|---|---|---|---|---|---|
| | | | Total | C | N | O | S | | |
| 16 | T | 104 | 809 | 523 | 134 | 146 | 6 | 0 | 0 |

- Molecule 17 is a protein called Uncharacterized protein.

| Mol | Chain | Residues | Atoms | | | | | AltConf | Trace |
|---|---|---|---|---|---|---|---|---|---|
| | | | Total | C | N | O | S | | |
| 17 | X | 353 | 2700 | 1706 | 467 | 510 | 17 | 0 | 0 |
| | | | Total | C | N | O | S | | |
| 17 | Y | 368 | 2821 | 1788 | 486 | 530 | 17 | 0 | 0 |

For Manuscript Review

### 3 Residue-property plots ⓘ

These plots are drawn for all protein, RNA, DNA and oligosaccharide chains in the entry. The first graphic for a chain summarises the proportions of the various outlier classes displayed in the second graphic. The second graphic shows the sequence view annotated by issues in geometry and atom inclusion in map density. Residues are color-coded according to the number of geometric quality criteria for which they contain at least one outlier: green = 0, yellow = 1, orange = 2 and red = 3 or more. A red diamond above a residue indicates a poor fit to the EM map for this residue (all-atom inclusion < 40%). Stretches of 2 or more consecutive residues without any outlier are shown as a green connector. Residues present in the sample, but not in the model, are shown in grey.

- Molecule 1: DNA-directed RNA polymerase

Chain A:

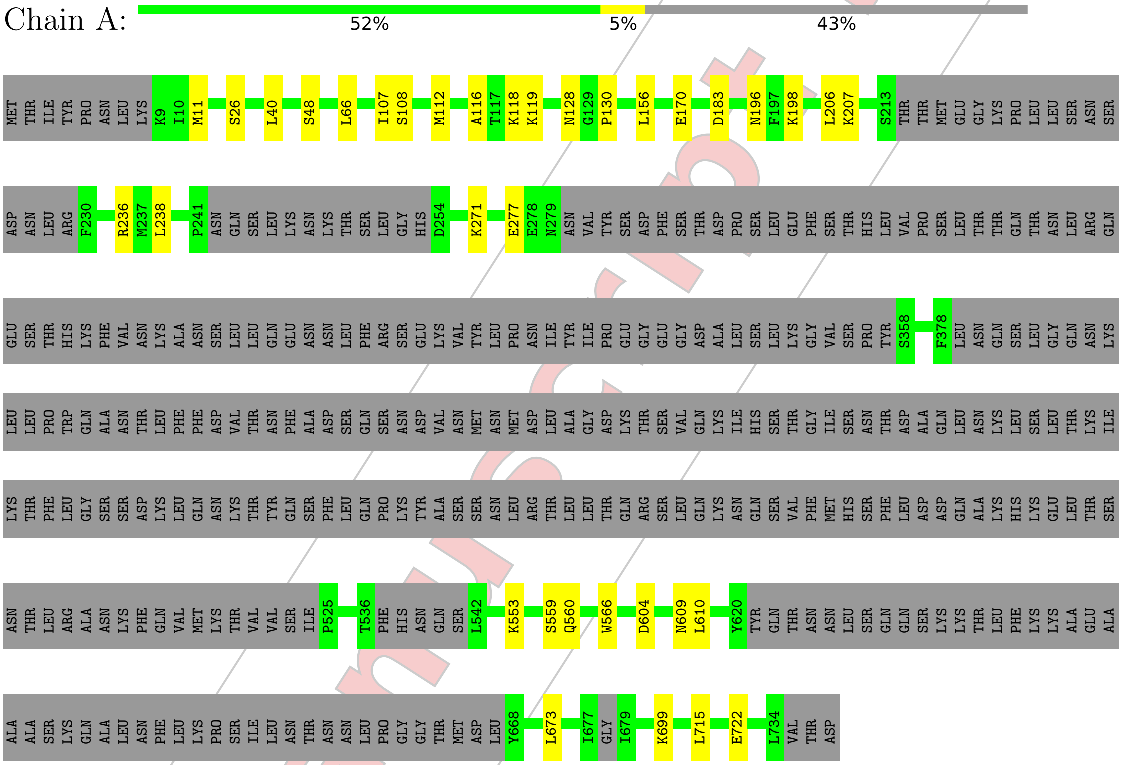


- Molecule 1: DNA-directed RNA polymerase

Chain Z:

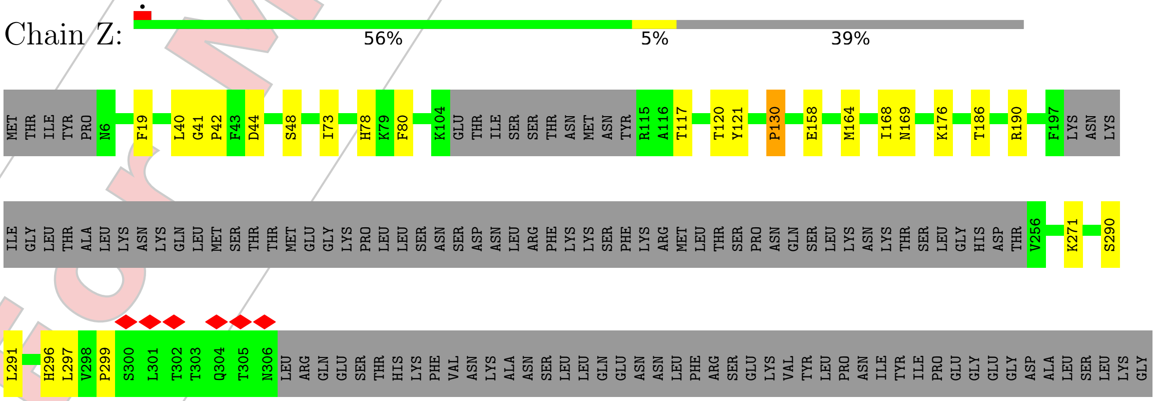

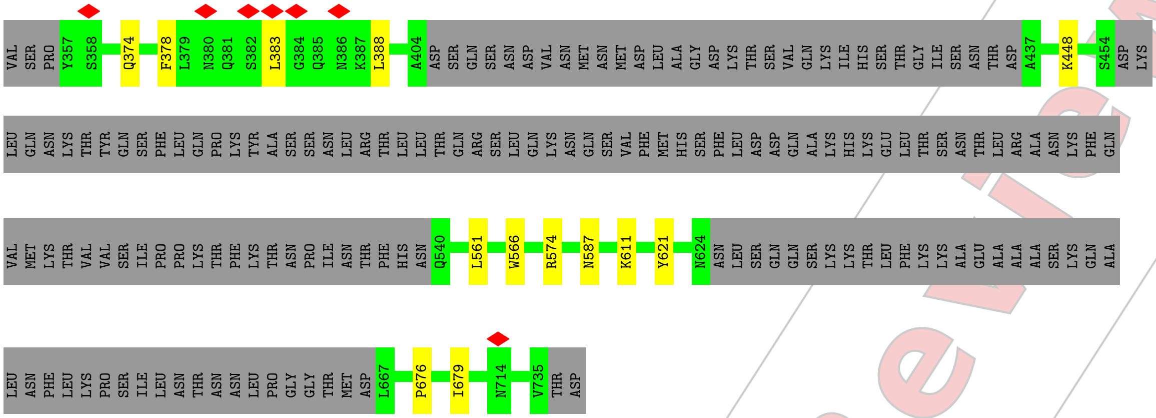


• Molecule 2: DNA-directed RNA polymerase subunit beta N-terminal section

Chain B:

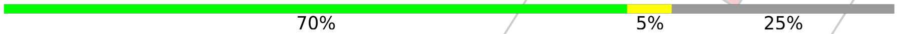


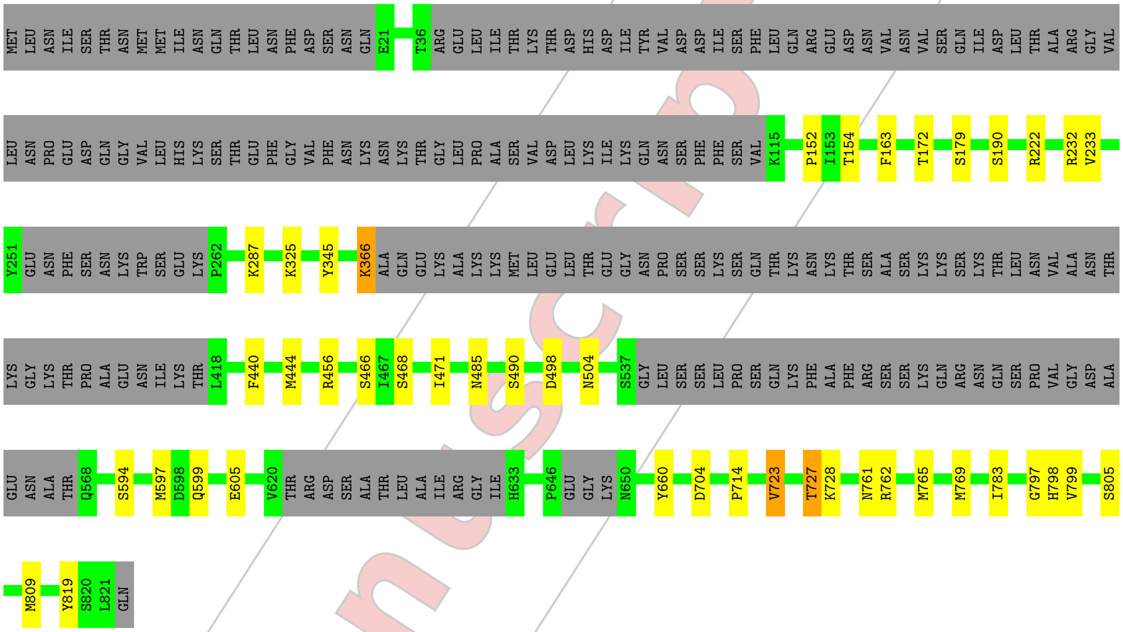


• Molecule 3: DNA-directed RNA polymerase subunit beta C-terminal section

Chain C:

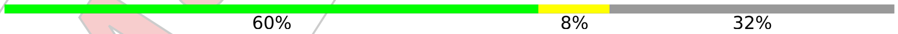


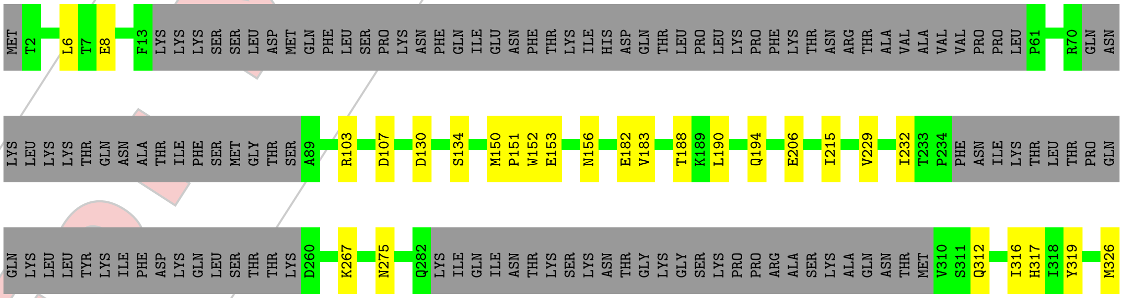


For Manuscript Review

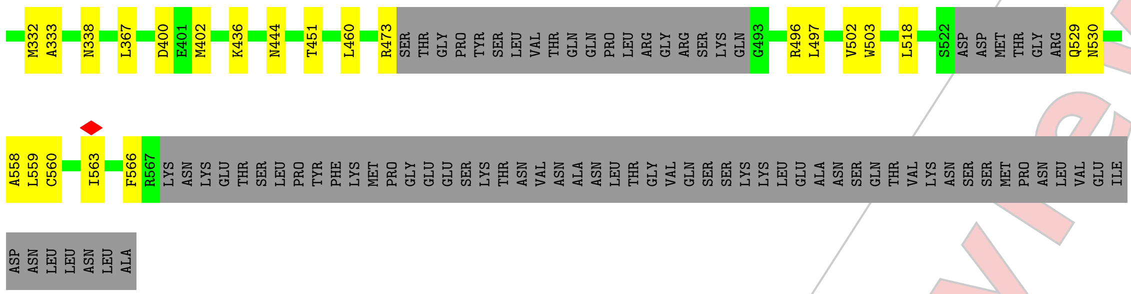

• Molecule 4: DNA-directed RNA polymerase subunit beta”

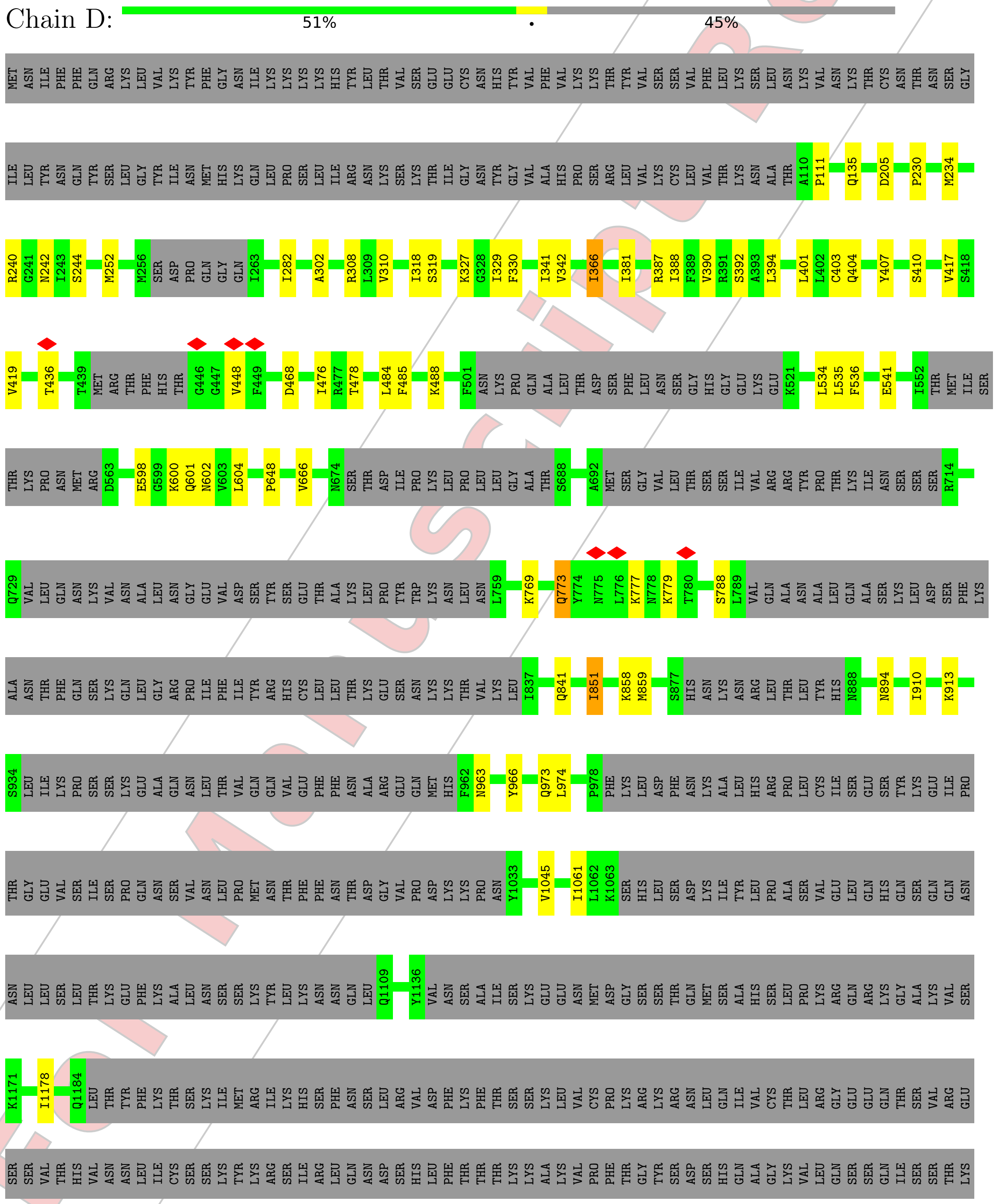


For Manuscript Review

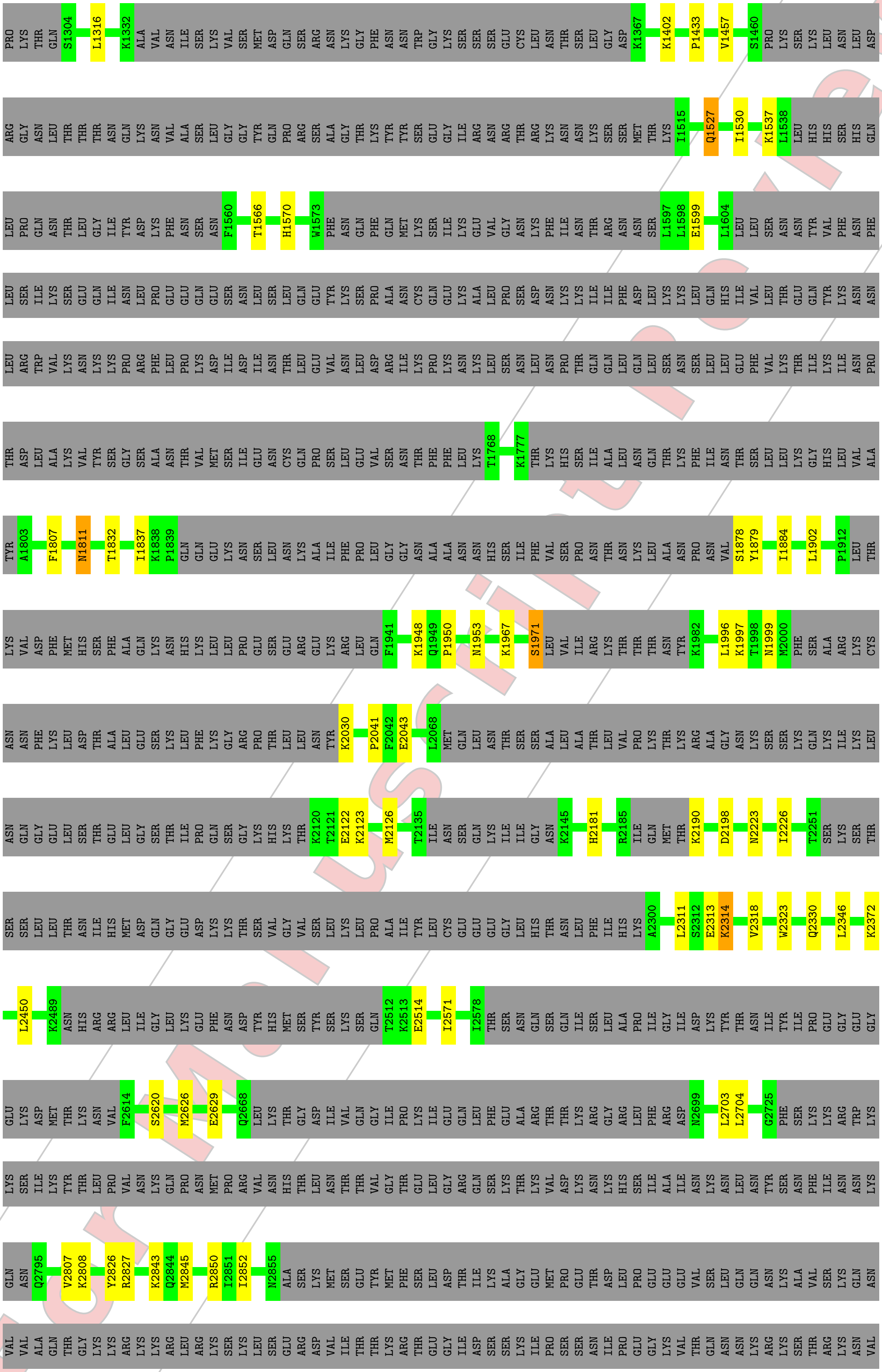


For Manuscript Review

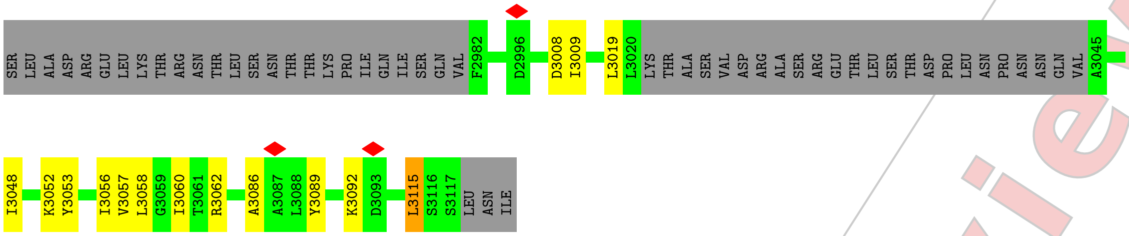


• Molecule 5: DNA-directed RNA polymerase subunit

Chain E:

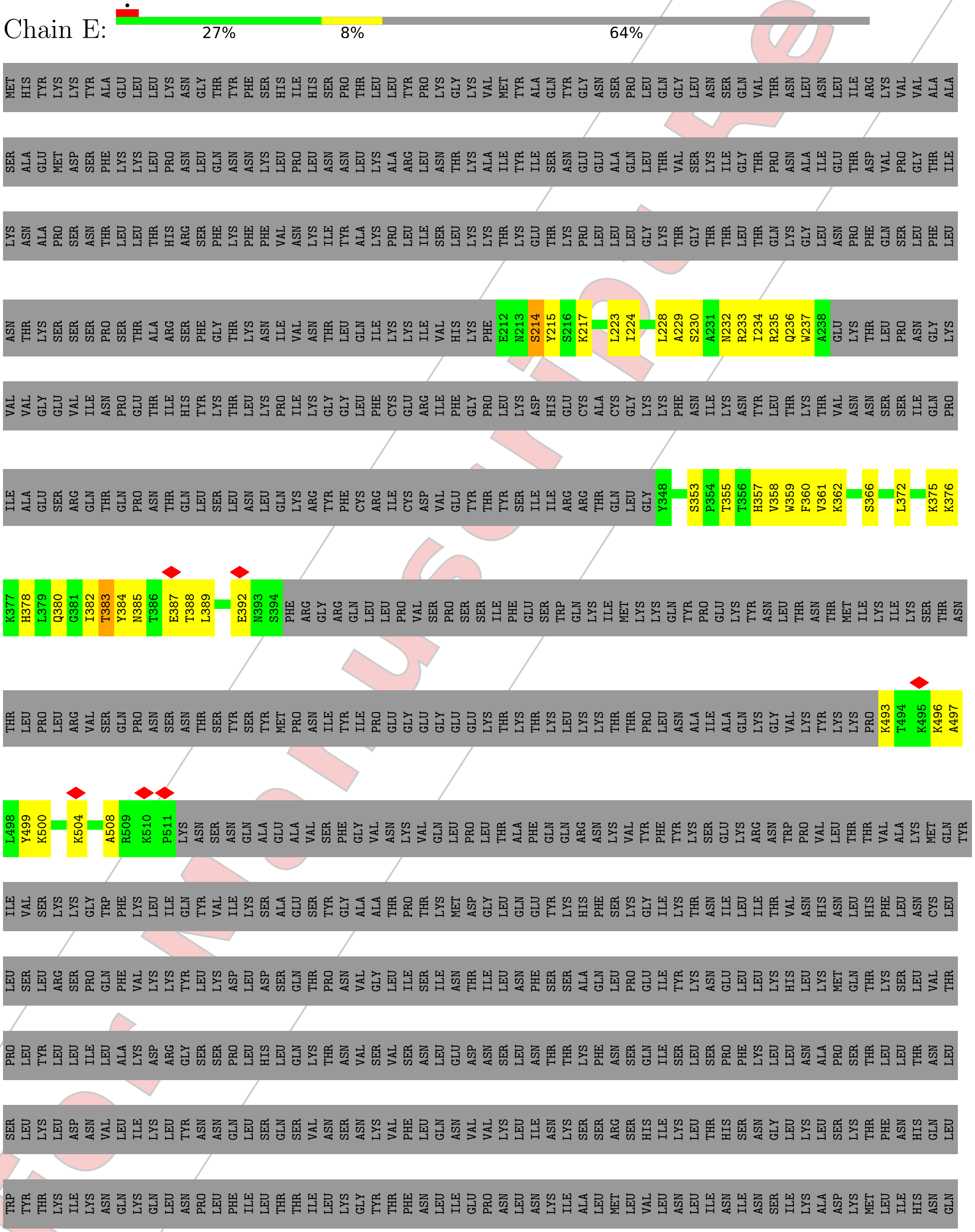


For Manuscript Review

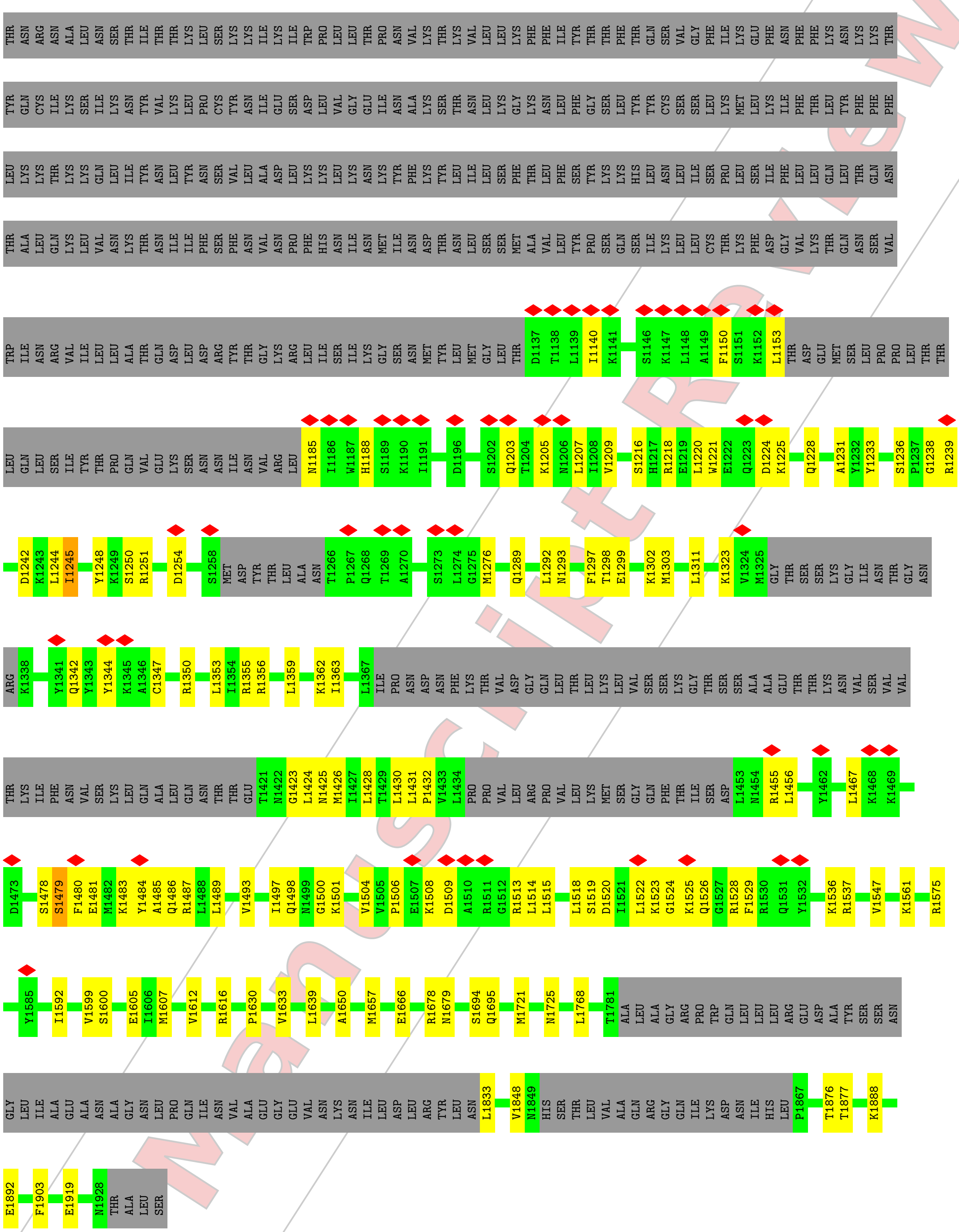


• Molecule 6: Glucose-6-phosphate 1-epimerase

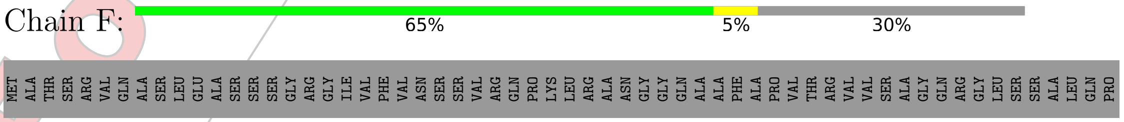


For Manuscript Review

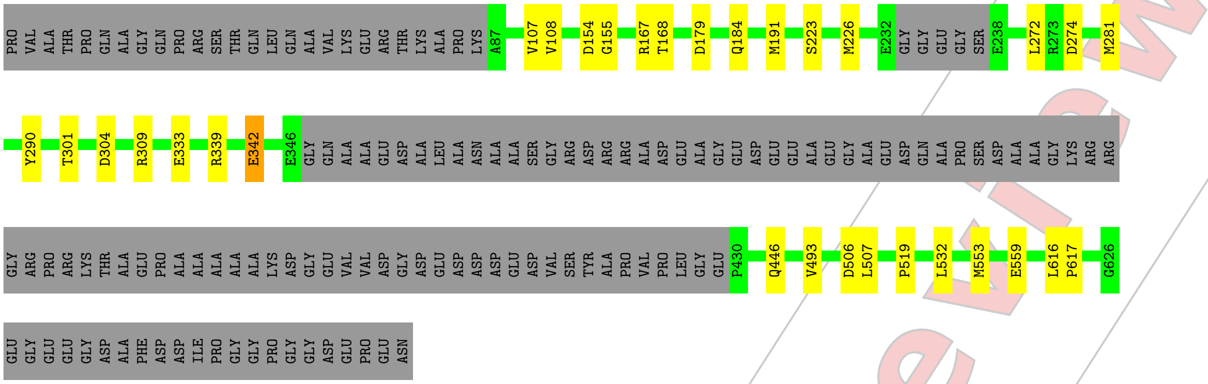


• Molecule 7: Rubisco LSMT substrate-binding domain-containing protein

Chain G:

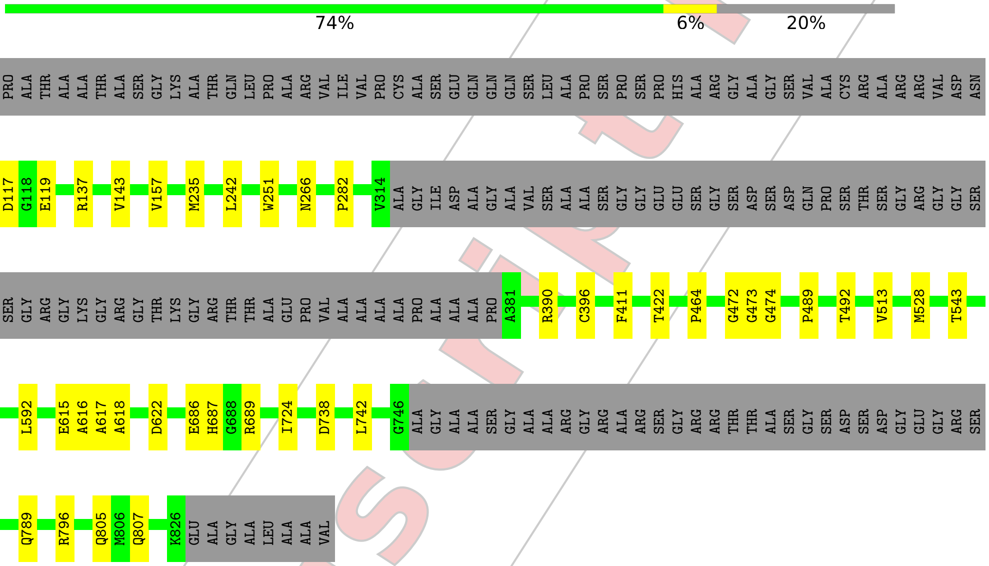


• Molecule 8: UDP-3-O-acyl-N-acetylglucosamine deacetylase

Chain H:

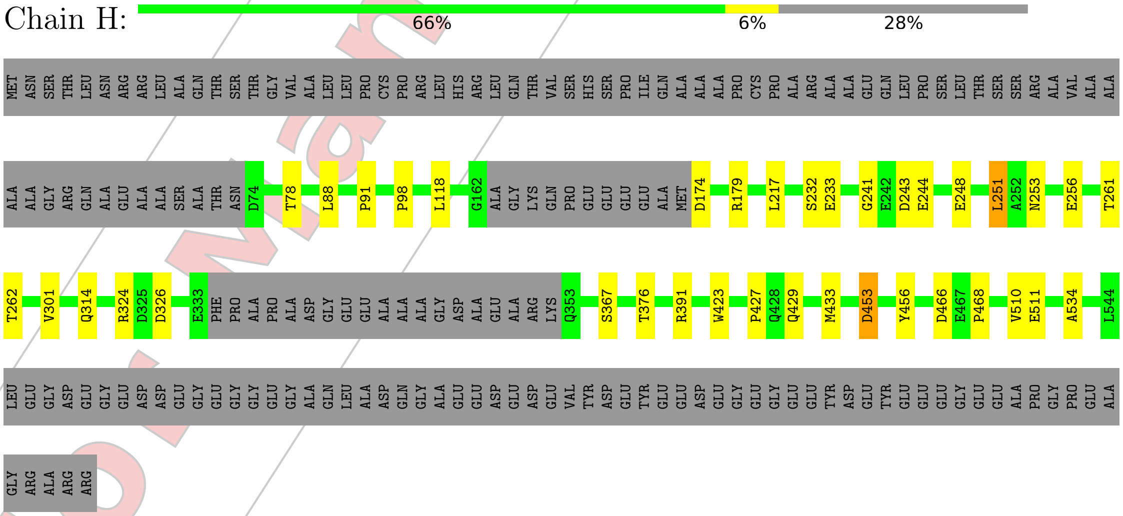


• Molecule 8: UDP-3-O-acyl-N-acetylglucosamine deacetylase

For Manuscript Review

Chain I: 47% 6% 47%

MET ASN SER THR LEU ASN ARG ARG LEU ALA GLN THR SER THR GLY VAL ALA LEU LEU PRO CYS PRO ARG LEU HIS ARG LEU GLN THR VAL SER HIS SER PRO ILE GLN ALA ALA ALA PRO CYS PRO ALA ARG ALA ALA GLU GLN LEU PRO SER LEU THR SER SER ARG ALA VAL ALA ALA

ALA ALA GLY ARG GLN ALA GLU ALA ALA SER ALA THR ASN ASP ARG LYS ARG THR ARG LEU ALA ALA ASP LYS SER PRO GLU LEU I89 D90 P91 Q92 M99 Q104 Q105 V116 G117 L118 H119 A133 V142 P143 E144 GLY THR ASN ALA ALA ARG GLY GLU GLU SER LEU VAL

ARG MET LEU GLU ALA GLY ALA GLY LYS GLN PRO GLU GLU GLU GLU ALA MET ASP ASP THR ALA THR ARG GLU TYR LYS ARG GLU LEU ILE ALA ALA TYR GLN GLU TYR VAL ASP ALA GLN GLU ASP ASP GLY PHE ASN GLY THR PHE ALA ASP TYR MET ASN GLY LYS SER TYR VAL ALA

LEU LEU GLN ARG LEU ARG GLU GLU GLY PRO PRO ARG PHE SER PHE SER GLU ALA GLU ALA PRO GLN PRO ARG GLY GLU ASP GLU PRO TYR VAL GLU ALA ALA LEU ALA ASN ILE T255 K266 I293 GLU GLY GLY ALA GLU V299 P300 V301 L318 A321 D325 D326 L327

G332 E333 PHE PRO ALA PRO ALA ASP GLY GLU GLU ALA ALA GLY ASP ALA GLU ALA ARG LYS Q353 K357 M358 R361 Q369 Q429 M433 W438 N451 D466 M473 H486 G487 H488 S489 G490 L491 D502 H503 E504 L505 V523 E546 GLY ASP GLU

GLY GLU ASP ASP GLU GLY GLU GLY GLY GLU GLY ALA GLN LEU ALA ASP GLN GLY ALA GLU GLU ASP GLU ASP GLU VAL TYR ASP GLU TYR GLU GLU ASP GLU GLY GLU GLY GLU GLU TYR ASP GLU TYR GLU GLU GLU GLY GLU ALA PRO GLY PRO GLU ALA GLY ARG ALA ARG ARG

• Molecule 9: Uncharacterized protein A0A2K3DRS4

Chain J: 35% • 61%

MET PRO ALA ALA ARG SER LEU ASP SER SER GLU CYS SER THR SER TYR SER ALA GLN HIS TYR ARG ARG THR SER ARG GLY GLU ARG MET ARG THR CYS ALA ALA ALA ASN ASP ASP ASP PHE LYS PRO ALA GLY ALA ALA ARG ARG GLY ARG GLY LYS ALA ARG ALA PRO PRO PRO PRO

PRO PRO PRO PRO GLN GLN GLY ALA GLU ASP GLU GLU GLU TYR GLU GLU SER LEU GLY ALA SER LEU GLU ASP PRO ALA GLN GLY GLN LEU GLN ARG LEU LEU SER ARG SER GLY PRO GLY GLU SER VAL ALA ALA ASP VAL GLY PRO ALA ILE LYS ARG PRO P116 Q130 L137

D172 Y190 R194 R195 M199 E206 A211 Q214 F234 V238 N241 I242 V246 R247 Q248 P249 L250 GLU GLU GLU LEU ARG GLU ARG LEU GLU GLU ALA ILE GLY GLU ALA THR LEU LEU PRO PRO THR GLU ALA GLU GLN GLN ALA PRO LYS PHE PRO SER ASP

ALA ASP VAL SER ALA TYR MET MET GLU ASN PHE LEU GLU GLY VAL LEU PRO GLY VAL SER LEU THR ASN PRO ASN GLN ARG TYR ALA ASP ASP ILE TYR GLY GLY ASP GLU GLU GLY ALA ASP GLU LEU LEU ASN MET VAL GLU GLU GLN GLU GLY ASP GLY GLU ALA GLU PRO ALA ASP

GLU

• Molecule 10: SAP domain-containing protein

Chain K: • 58% • 38%

MET LEU HIS VAL GLY GLN ALA SER ALA GLY ARG PRO VAL ALA SER SER ARG SER VAL ALA ARG GLY PRO ARG PRO LEU GLN CYS PRO VAL SER THR PHE ALA PRO SER SER ARG ARG LEU ILE ALA GLU ALA LYS ARG GLY GLY ARG PRO THR SER GLU PRO SER ALA ALA GLY ASP ASP

ALA ASP PRO LEU ALA ALA VAL ALA GLU LEU ASN PHE ARG GLU LEU GLN ALA GLU LEU LYS ALA ARG GLY LEU PRO ALA GLY GLY LYS ARG GLU GLU LEU ALA GLU ARG LEU ALA ALA ALA ALA LEU ALA ASP GLU GLY ALA GLY ALA GLY ALA SER ALA THR GLY GLY SER SER SER SER SER

GLY SER GLY SER GLY ASN ALA LYS ALA ALA ALA GLY ILE ASP LEU ASP SER ALA LEU ASP LEU LEU GLU GLU VAL PRO TYR ALA GLU LEU LYS ALA LEU CYS ALA GLU ALA GLY LEU SER ALA ALA GLY LYS LYS ALA GLU LEU GLN GLN ARG LEU ALA GLU ALA ALA VAL ALA GLY ASP

ILE ASP LEU VAL SER PRO LEU GLU GLY GLY GLU ASP LEU GLU GLN PRO GLU ALA ALA ARG ALA ALA SER HIS GLY SER ALA GLN SER SER ALA ALA ALA ALA ALA ASP ALA ALA ALA LEU LEU PRO GLU LEU LEU ALA LEU SER ALA GLU GLU LEU GLU ALA ALA SER LEU SER GLU TYR GLY VAL

For Manuscript Review

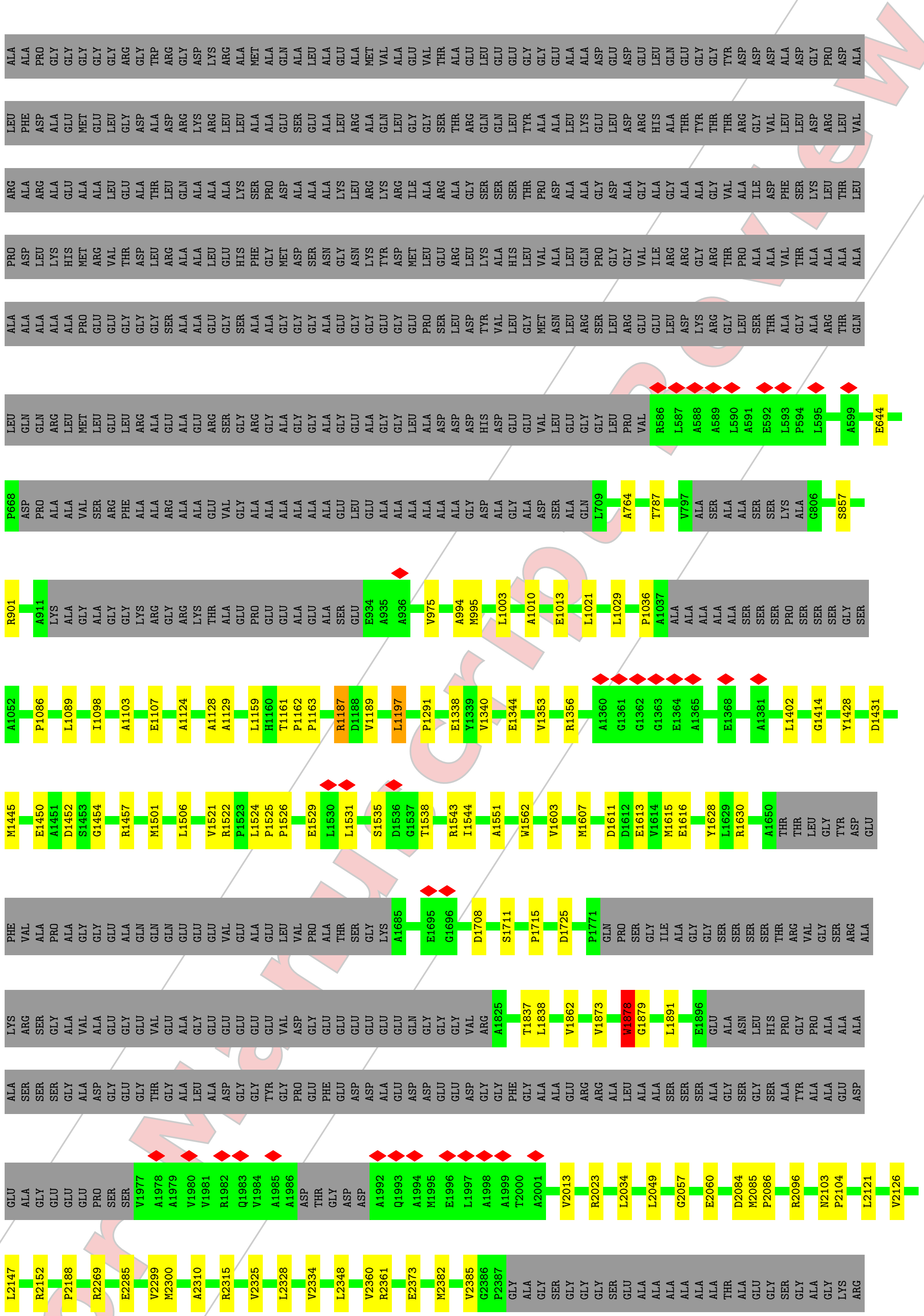


For Manuscript Review

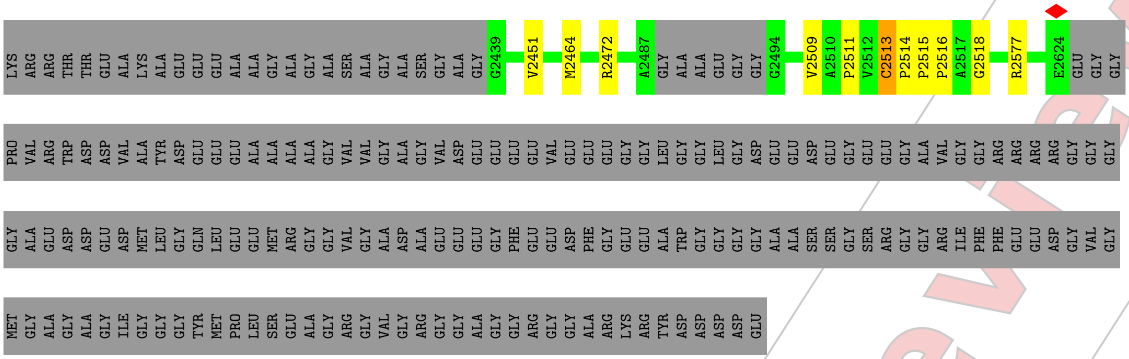


- Molecule 11: Uncharacterized protein A0A2K3D3G3

Chain M:

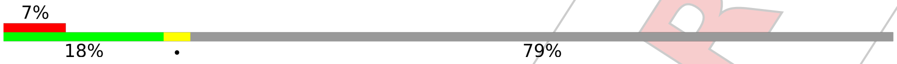


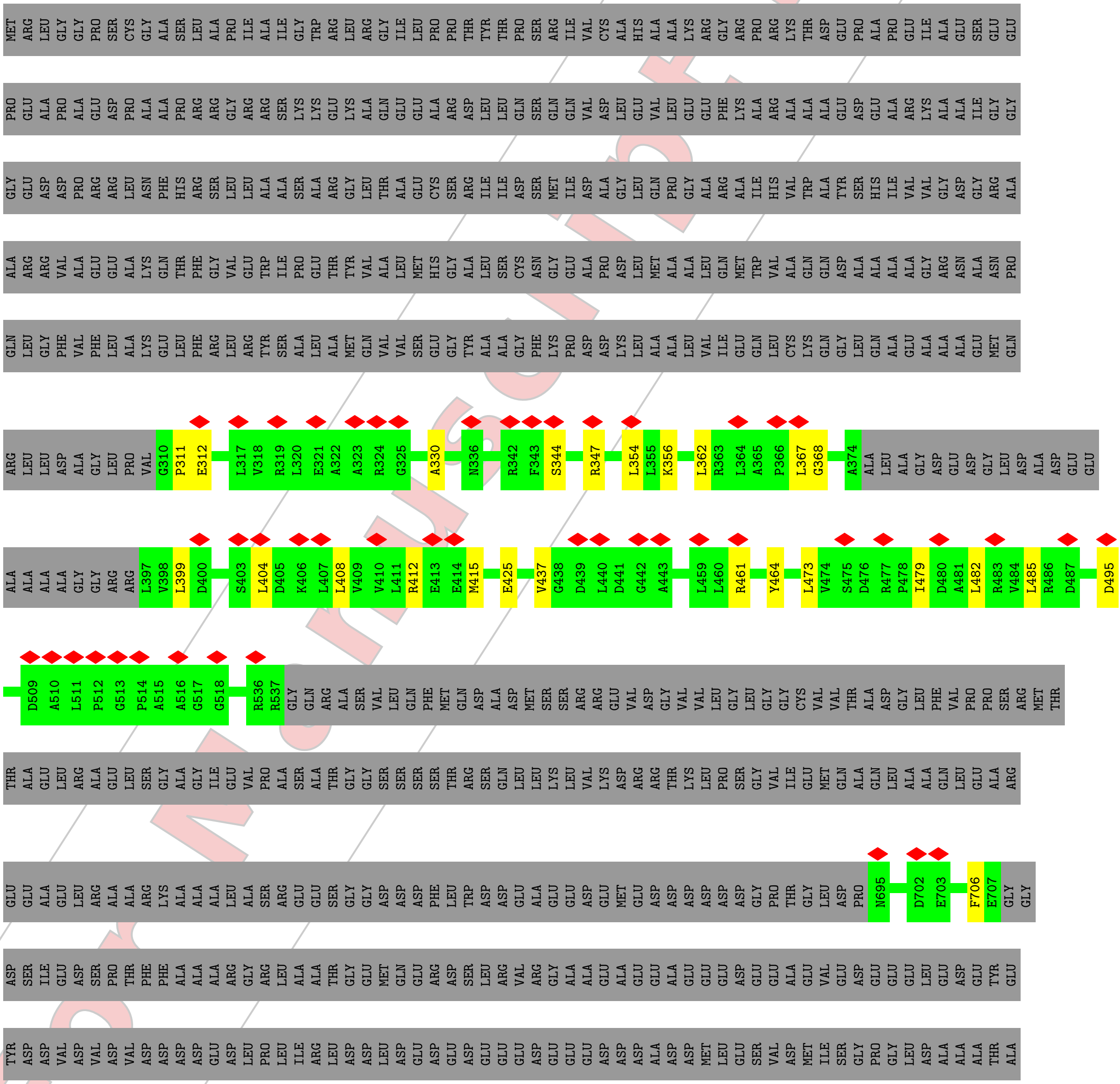


For Manuscript Review

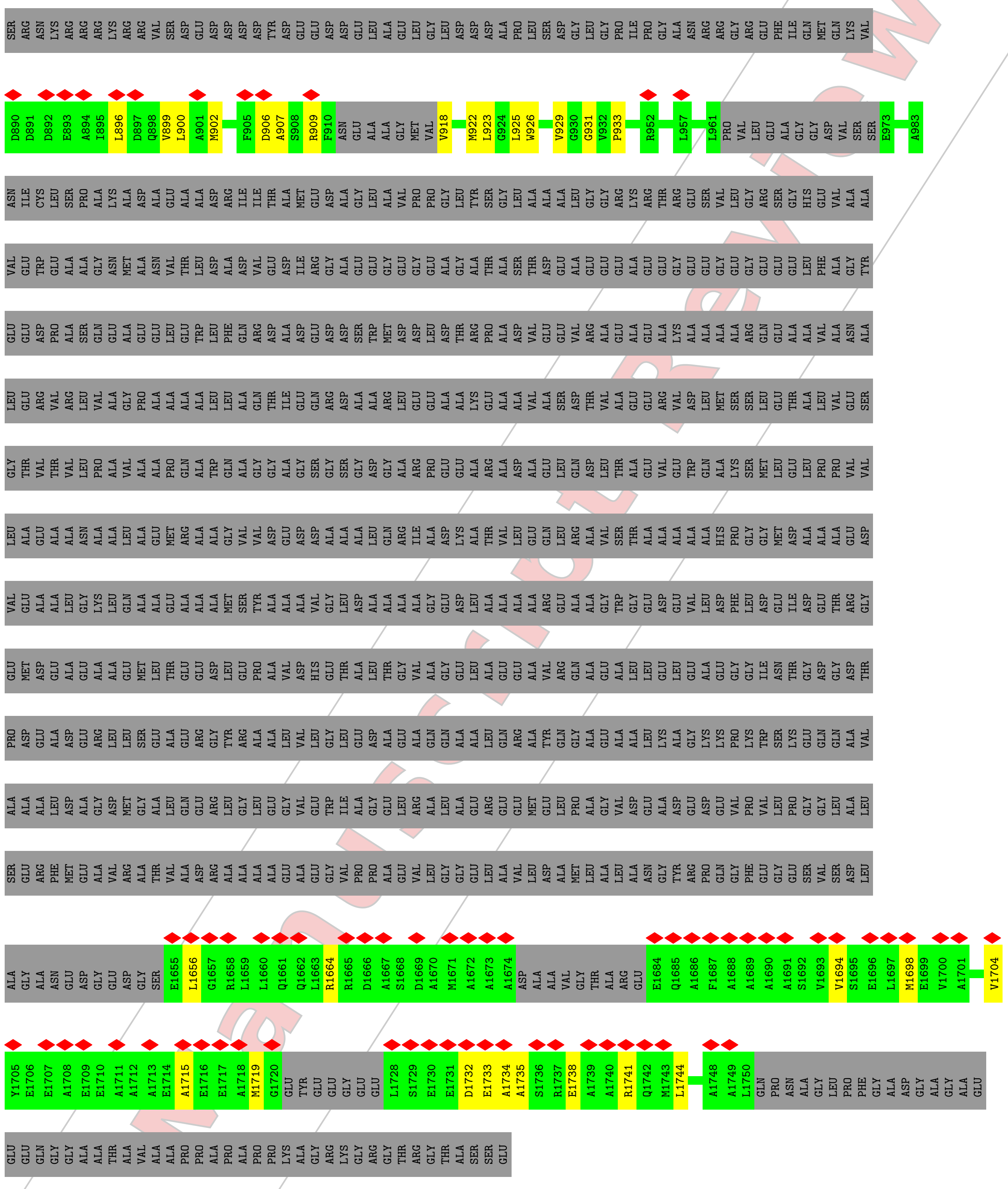

- Molecule 12: Mur ligase central domain-containing protein

Chain N:

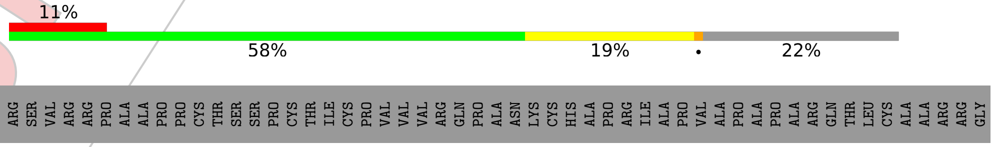


For Manuscript Review

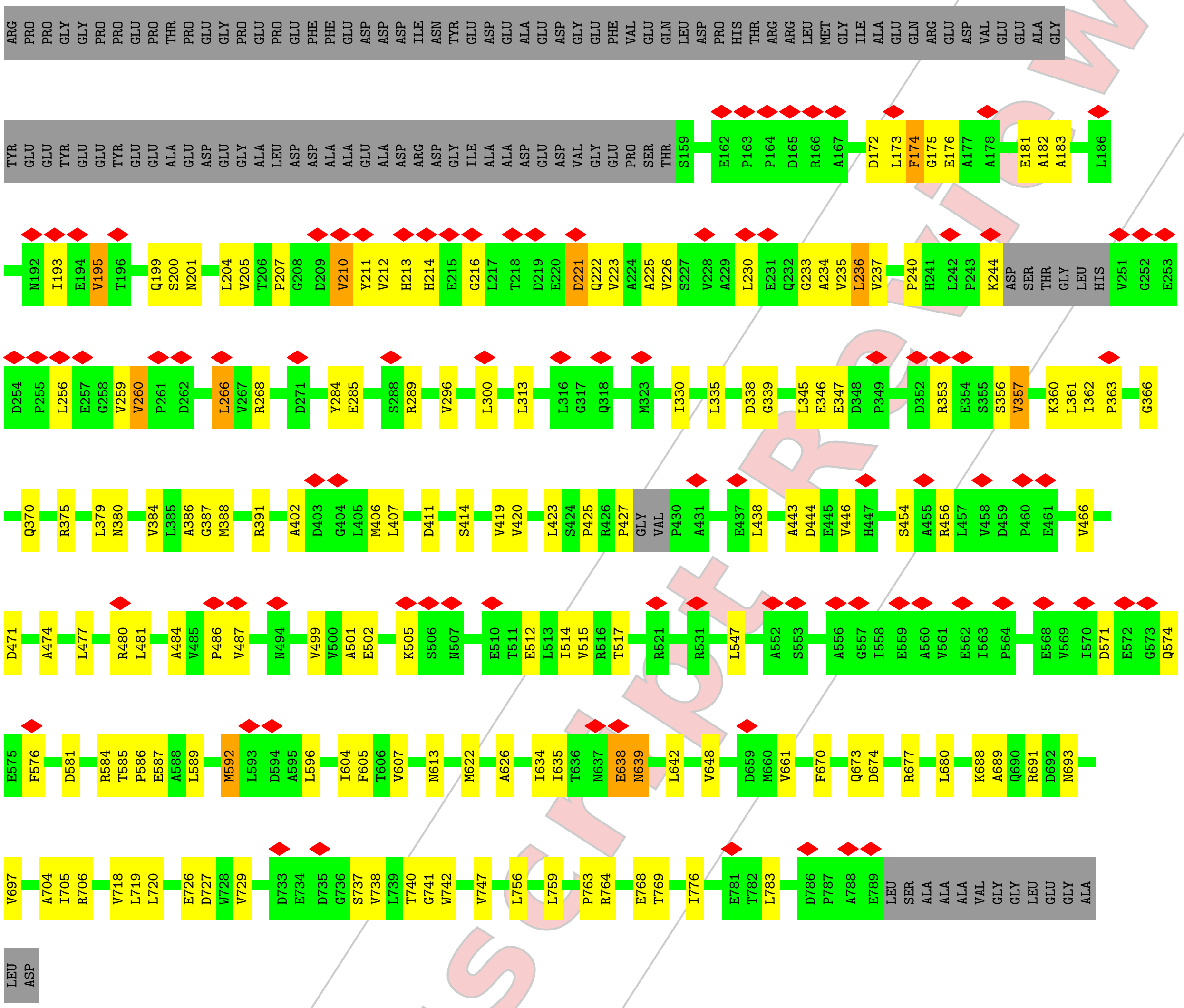


• Molecule 13: Coenzyme Q-binding protein COQ10 START domain-containing protein

Chain Q:

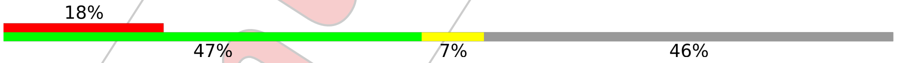


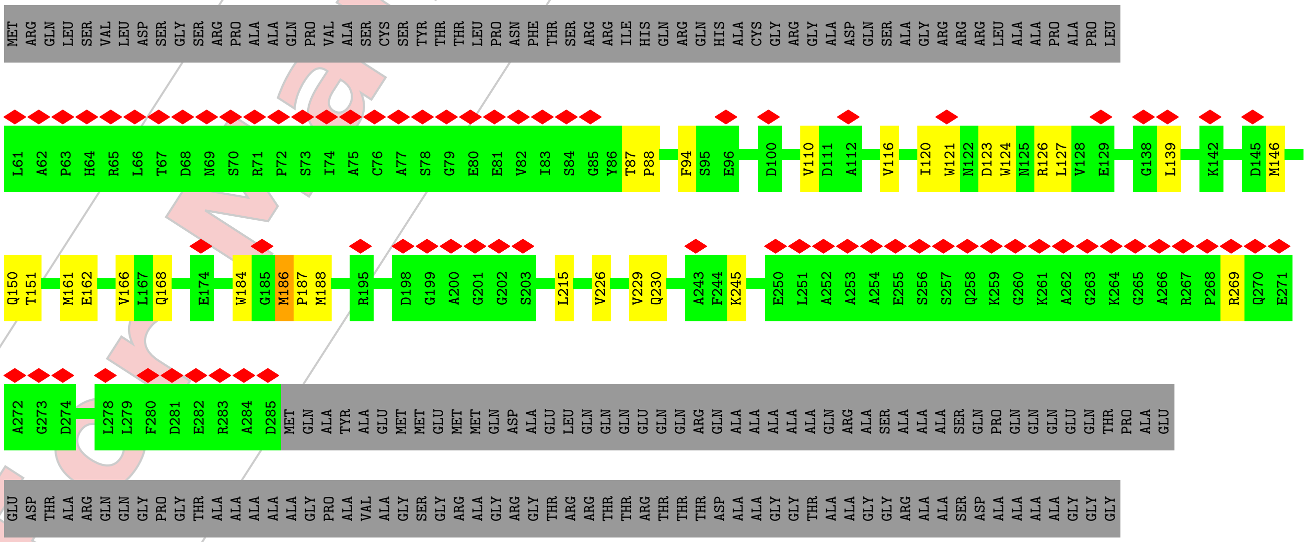


For Manuscript Review

GLN GLY PRO ASP ASP ARG SER SER GLY ARG LYS ARG SER VAL ARG SER THR ARG GLY ALA SER

• Molecule 14: S1 motif domain-containing protein

Chain R: 81% • 15%

MET GLN GLY ALA LEU ARG GLY LEU GLY ALA ARG PRO PRO THR ASN CYS GLY GLY ALA SER CYS SER GLY ARG VAL GLY PRO CYS SER ALA GLN LEU ASN PRO ARG GLY MET VAL PRO THR VAL GLY ARG GLY HIS GLN HIS ARG ILE LEU PRO PRO CYS ARG I55 T58 D91

R100 D106 V114 W130 N131 S132 H153 M180 W185 H188 I194 D199 Q297 G355 LEU

• Molecule 15: Uncharacterized protein

Chain S: 73% 6% 20%

MET ASP CYS SER ALA LEU ALA ALA ARG ARG ALA LEU SER GLN CYS ALA GLY PRO SER SER ARG CYS LEU VAL PRO ALA ALA LEU PRO GLN SER ARG PRO ARG THR CYS SER ARG SER THR GLY HIS ALA ALA GLY HIS ARG ALA THR THR ILE VAL ALA ARG ALA ASN GLY LYS THR VAL

THR PRO SER PRO LEU PHE ASP GLU VAL ALA E71 S83 D98 N108 E112 I116 E120 P126 V137 M172 W186 D187 K194 H261 D273 L288 R291 D318 E319 A320 E321 A322 E323 L324 E334 I337 R357 LEU GLY

• Molecule 16: Uncharacterized protein

Chain T: 19% 6% • 73%

MET GLN LEU GLY ARG GLN LEU SER ALA HIS ALA GLY THR CYS ARG GLN GLN ALA LEU ARG CYS SER SER LEU ALA ARG SER ALA PRO PRO PRO ALA ILE CYS PRO SER ARG ALA PRO THR GLN PRO CYS SER ARG GLY SER VAL GLN LEU HIS ALA LYS ARG THR GLY GLY ARG SER LYS

THR GLN GLU GLY SER ASP ASP ALA SER GLY TRP ASP ALA ALA LEU ALA ASP ILE PRO THR GLU ASP ASP GLU ALA ALA SER GLU ASP PHE GLU GLU SER ASP ALA GLU ARG SER ALA GLY GLN ALA GLY GLU ALA GLY GLU PRO SER THR SER TYR ALA PRO VAL GLY LYS VAL SER SER

GLY PRO ARG ALA PRO LYS ALA PRO SER SER LEU LEU SER LEU ALA GLU TYR ASN MET LEU MET GLU ALA LYS ALA ILE ARG GLU ALA GLN GLU GLU ALA GLU ARG LEU GLU ARG GLU ILE ALA GLU GLY ARG GLY VAL VAL VAL THR GLU PRO VAL PRO ILE ASP VAL ASN ARG GLN GLY

PHE GLN VAL F184 E192 L193 L194 VAL ALA GLN ASP GLY GLN GLY LEU LEU ALA LEU ALA ASP SER TYR VAL ALA ALA ARG GLN PRO ALA ALA GLU GLY GLY TRP ALA TYR SER ARG VAL LYS LEU GLU GLY GLU VAL PRO THR GLY ALA SER VAL TYR THR V241 M252 M270 A271

P272 A273 ASP ASP ASP SER GLY GLU A280 V287 F288 D291 G292 K293 S294 R295 T296 M297 A298 L299 S300 E301 V302 R303 L304 V305 G306 Q307 P308 P309 L310 P313 V314 T323 L324 L330 P331 I336 N339 THR ALA VAL ASP LEU ALA ASP MET SER GLU GLU GLU LEU ALA

GLU ARG LEU ALA GLU LEU GLU ALA GLN ALA GLU ALA LEU ASP GLU LEU ASP GLU ALA PRO ASP VAL PRO SER VAL ASP ASN TYR

• Molecule 17: Uncharacterized protein

Chain X: 81% 5% 14%

MET GLU ALA SER CYS SER SER HIS CYS GLN GLY ALA ALA THR SER ARG LEU GLY GLN ARG GLY TRP ARG MET PRO PRO ALA ARG VAL LEU ARG LEU ARG MET GLN ALA SER MET PRO VAL GLU ILE GLY LYS LEU GLU LYS GLN PRO ILE PRO VAL PRO LYS ASP TYR LYS P58 S59

For Manuscript Review

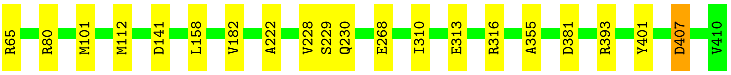


• Molecule 17: Uncharacterized protein

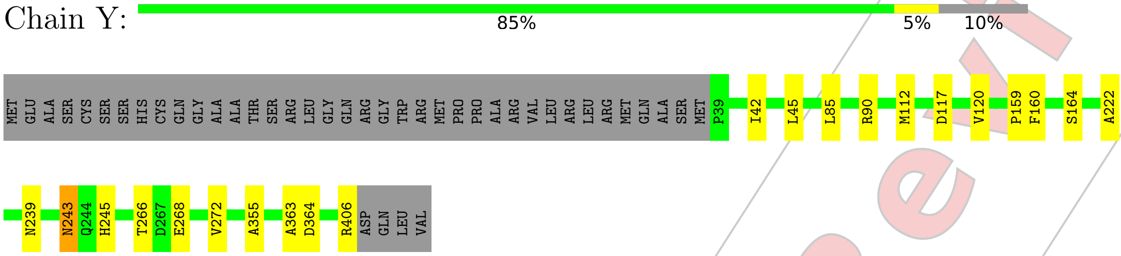


For Manuscript Review

### 4 Experimental information (i)

| Property | Value | Source |
|---|---|---|
| EM reconstruction method | SINGLE PARTICLE | Depositor |
| Imposed symmetry | POINT, Not provided | |
| Number of particles used | 650394 | Depositor |
| Resolution determination method | FSC 0.143 CUT-OFF | Depositor |
| CTF correction method | PHASE FLIPPING AND AMPLITUDE CORRECTION | Depositor |
| Microscope | TFS KRIOS | Depositor |
| Voltage (kV) | 300 | Depositor |
| Electron dose ($e^-/\text{Å}^2$) | 60 | Depositor |
| Minimum defocus (nm) | 500 | Depositor |
| Maximum defocus (nm) | 2000 | Depositor |
| Magnification | Not provided | |
| Image detector | FEI FALCON IV (4k x 4k) | Depositor |
| Maximum map value | 0.623 | Depositor |
| Minimum map value | 0.093 | Depositor |
| Average map value | 0.201 | Depositor |
| Map value standard deviation | 0.008 | Depositor |
| Recommended contour level | 0.235 | Depositor |
| Map size (Å) | 558.6, 558.6, 558.6 | wwPDB |
| Map dimensions | 588, 588, 588 | wwPDB |
| Map angles (°) | 90.0, 90.0, 90.0 | wwPDB |
| Pixel spacing (Å) | 0.95, 0.95, 0.95 | Depositor |

For Manuscript Review

### 5 Model quality (i)

#### 5.1 Standard geometry (i)

The Z score for a bond length (or angle) is the number of standard deviations the observed value is removed from the expected value. A bond length (or angle) with $|Z| > 5$ is considered an outlier worth inspection. RMSZ is the root-mean-square of all Z scores of the bond lengths (or angles).

| Mol | Chain | Bond lengths | | Bond angles | |
|---|---|---|---|---|---|
| | | RMSZ | #$|Z|$ >5 | RMSZ | #$|Z|$ >5 |
| 1 | A | 0.15 | 0/3422 | 0.35 | 0/4605 |
| 1 | Z | 0.15 | 0/3649 | 0.41 | 1/4929 (0.0%) |
| 2 | B | 0.17 | 0/5073 | 0.36 | 1/6846 (0.0%) |
| 3 | C | 0.16 | 0/3434 | 0.38 | 0/4633 |
| 4 | D | 0.16 | 0/14417 | 0.38 | 0/19397 |
| 5 | E | 0.20 | 0/5810 | 0.51 | 0/7825 |
| 6 | F | 0.18 | 0/3607 | 0.40 | 1/4909 (0.0%) |
| 7 | G | 0.15 | 0/5252 | 0.32 | 0/7154 |
| 8 | H | 0.14 | 0/3499 | 0.34 | 0/4760 |
| 8 | I | 0.14 | 0/2568 | 0.38 | 0/3500 |
| 9 | J | 0.16 | 0/1131 | 0.37 | 0/1530 |
| 10 | K | 0.15 | 0/12566 | 0.34 | 0/17224 |
| 11 | M | 0.13 | 0/2824 | 0.36 | 0/3817 |
| 12 | N | 0.17 | 0/4829 | 0.55 | 2/6586 (0.0%) |
| 13 | Q | 0.12 | 0/1768 | 0.35 | 0/2400 |
| 14 | R | 0.15 | 0/2502 | 0.32 | 0/3404 |
| 15 | S | 0.29 | 0/2433 | 0.36 | 0/3308 |
| 16 | T | 0.47 | 1/825 (0.1%) | 0.80 | 4/1114 (0.4%) |
| 17 | X | 0.18 | 0/2761 | 0.37 | 0/3767 |
| 17 | Y | 0.16 | 0/2887 | 0.37 | 0/3938 |
| All | All | 0.17 | 1/85257 (0.0%) | 0.39 | 9/115646 (0.0%) |

Chiral center outliers are detected by calculating the chiral volume of a chiral center and verifying if the center is modelled as a planar moiety or with the opposite hand.A planarity outlier is detected by checking planarity of atoms in a peptide group, atoms in a mainchain group or atoms of a sidechain that are expected to be planar.

| Mol | Chain | #Chirality outliers | #Planarity outliers |
|---|---|---|---|
| 1 | A | 0 | 1 |
| 5 | E | 0 | 2 |
| 8 | I | 0 | 1 |
| 10 | K | 0 | 3 |
| 12 | N | 0 | 1 |

*Continued on next page...*

For Manuscript Review

*Continued from previous page...*

| Mol | Chain | #Chirality outliers | #Planarity outliers |
|---|---|---|---|
| All | All | 0 | 8 |

All (1) bond length outliers are listed below:

| Mol | Chain | Res | Type | Atoms | Z | Observed(Å) | Ideal(Å) |
|---|---|---|---|---|---|---|---|
| 16 | T | 305 | VAL | CA-C | -7.21 | 1.44 | 1.52 |

All (9) bond angle outliers are listed below:

| Mol | Chain | Res | Type | Atoms | Z | Observed($^o$) | Ideal($^o$) |
|---|---|---|---|---|---|---|---|
| 16 | T | 305 | VAL | N-CA-C | -13.36 | 97.59 | 110.42 |
| 12 | N | 638 | GLU | CA-C-N | 8.76 | 137.47 | 121.70 |
| 12 | N | 638 | GLU | C-N-CA | 8.76 | 137.47 | 121.70 |
| 16 | T | 305 | VAL | CB-CA-C | -7.03 | 102.97 | 111.97 |
| 16 | T | 298 | ALA | N-CA-C | 5.64 | 118.42 | 107.71 |
| 1 | Z | 41 | GLY | N-CA-C | -5.56 | 97.18 | 113.30 |
| 2 | B | 366 | LYS | CB-CG-CD | 5.17 | 123.18 | 111.30 |
| 16 | T | 301 | GLU | N-CA-C | 5.07 | 117.00 | 108.73 |
| 6 | F | 342 | GLU | CA-CB-CG | 5.01 | 124.13 | 114.10 |

There are no chirality outliers.

All (8) planarity outliers are listed below:

| Mol | Chain | Res | Type | Group |
|---|---|---|---|---|
| 1 | A | 40 | LEU | Peptide |
| 5 | E | 1224 | ASP | Peptide |
| 5 | E | 383 | THR | Peptide |
| 8 | I | 142 | VAL | Peptide |
| 10 | K | 1161 | THR | Peptide |
| 10 | K | 1187 | ARG | Peptide |
| 10 | K | 2513 | CYS | Peptide |
| 12 | N | 639 | ASN | Peptide |

#### 5.2 Too-close contacts (i)

In the following table, the Non-H and H(model) columns list the number of non-hydrogen atoms and hydrogen atoms in the chain respectively. The H(added) column lists the number of hydrogen atoms added and optimized by MolProbity. The Clashes column lists the number of clashes within the asymmetric unit, whereas Symm-Clashes lists symmetry-related clashes.

For Manuscript Review

| Mol | Chain | Non-H | H(model) | H(added) | Clashes | Symm-Clashes |
|---|---|---|---|---|---|---|
| 1 | A | 3368 | 0 | 3524 | 21 | 0 |
| 1 | Z | 3590 | 0 | 3698 | 22 | 0 |
| 2 | B | 4970 | 0 | 5159 | 27 | 0 |
| 3 | C | 3368 | 0 | 3379 | 32 | 0 |
| 4 | D | 14136 | 0 | 14711 | 93 | 0 |
| 5 | E | 5698 | 0 | 5930 | 108 | 0 |
| 6 | F | 3517 | 0 | 3392 | 17 | 0 |
| 7 | G | 5136 | 0 | 5032 | 22 | 0 |
| 8 | H | 3419 | 0 | 3274 | 23 | 0 |
| 8 | I | 2506 | 0 | 2418 | 19 | 0 |
| 9 | J | 1105 | 0 | 1045 | 11 | 0 |
| 10 | K | 12323 | 0 | 12188 | 57 | 0 |
| 11 | M | 2794 | 0 | 2758 | 31 | 0 |
| 12 | N | 4733 | 0 | 4669 | 105 | 0 |
| 13 | Q | 1730 | 0 | 1693 | 17 | 0 |
| 14 | R | 2438 | 0 | 2273 | 10 | 0 |
| 15 | S | 2374 | 0 | 2168 | 15 | 0 |
| 16 | T | 809 | 0 | 826 | 30 | 0 |
| 17 | X | 2700 | 0 | 2694 | 11 | 0 |
| 17 | Y | 2821 | 0 | 2829 | 15 | 0 |
| All | All | 83535 | 0 | 83660 | 621 | 0 |

The all-atom clashscore is defined as the number of clashes found per 1000 atoms (including hydrogen atoms). The all-atom clashscore for this structure is 4.

All (621) close contacts within the same asymmetric unit are listed below, sorted by their clash magnitude.

| Atom-1 | Atom-2 | Interatomic distance (Å) | Clash overlap (Å) |
|---|---|---|---|
| 8:I:353:GLN:HE21 | 8:I:353:GLN:N | 1.33 | 1.27 |
| 8:I:353:GLN:N | 8:I:353:GLN:NE2 | 2.12 | 0.98 |
| 16:T:307:GLN:HB2 | 16:T:308:PRO:HD2 | 1.45 | 0.94 |
| 5:E:380:GLN:O | 5:E:384:TYR:HB2 | 1.71 | 0.89 |
| 5:E:380:GLN:O | 5:E:383:THR:O | 1.93 | 0.87 |
| 12:N:638:GLU:HB3 | 12:N:639:ASN:HB2 | 1.60 | 0.82 |
| 4:D:2372:LYS:H | 4:D:2626:MET:HE1 | 1.43 | 0.82 |
| 16:T:306:GLY:O | 16:T:307:GLN:HB3 | 1.81 | 0.81 |
| 6:F:226:MET:HE3 | 6:F:226:MET:H | 1.46 | 0.80 |
| 16:T:307:GLN:CB | 16:T:308:PRO:HD2 | 2.07 | 0.78 |
| 13:Q:126:ARG:HH21 | 13:Q:269:ARG:HH21 | 1.34 | 0.74 |
| 16:T:304:LEU:HD12 | 16:T:324:LEU:HD23 | 1.69 | 0.74 |
| 16:T:299:LEU:HG | 16:T:300:SER:H | 1.52 | 0.74 |
| 4:D:666:VAL:HB | 4:D:1832:THR:HG22 | 1.70 | 0.72 |

Continued on next page...

For Manuscript Review

*Continued from previous page...*

| Atom-1 | Atom-2 | Interatomic distance (Å) | Clash overlap (Å) |
|---|---|---|---|
| 16:T:288:PHE:HA | 16:T:299:LEU:HA | 1.70 | 0.72 |
| 5:E:1292:LEU:HB3 | 5:E:1423:GLY:HA3 | 1.71 | 0.71 |
| 10:K:995:MET:HE1 | 10:K:1129:ALA:HB1 | 1.75 | 0.69 |
| 4:D:2311:LEU:HD23 | 4:D:2346:LEU:HD13 | 1.75 | 0.69 |
| 5:E:1239:ARG:HH21 | 12:N:353:ARG:HB2 | 1.58 | 0.68 |
| 5:E:1721:MET:HE2 | 1:Z:176:LYS:HG2 | 1.74 | 0.68 |
| 6:F:532:LEU:HD13 | 6:F:553:MET:HE2 | 1.77 | 0.67 |
| 12:N:571:ASP:HA | 12:N:574:GLN:HE22 | 1.61 | 0.66 |
| 12:N:345:LEU:HB2 | 12:N:366:GLY:HA2 | 1.77 | 0.66 |
| 11:M:1694:VAL:HG13 | 11:M:1698:MET:HE2 | 1.77 | 0.65 |
| 11:M:1741:ARG:HE | 11:M:1744:LEU:HD12 | 1.60 | 0.65 |
| 16:T:307:GLN:NE2 | 16:T:310:LEU:HG | 2.11 | 0.65 |
| 8:I:357:LYS:HE2 | 8:I:489:SER:HB2 | 1.79 | 0.65 |
| 10:K:2315:ARG:HG3 | 10:K:2325:VAL:HG12 | 1.79 | 0.65 |
| 17:X:112:MET:HB3 | 17:Y:112:MET:HB3 | 1.77 | 0.65 |
| 15:S:323:GLU:O | 15:S:324:LEU:C | 2.36 | 0.65 |
| 10:K:2300:MET:HE1 | 10:K:2310:ALA:HB3 | 1.79 | 0.65 |
| 10:K:2013:VAL:O | 10:K:2049:LEU:HA | 1.97 | 0.65 |
| 4:D:598:GLU:HB2 | 4:D:600:LYS:HG2 | 1.79 | 0.64 |
| 1:Z:378:PHE:HB2 | 1:Z:383:LEU:HD13 | 1.79 | 0.64 |
| 3:C:518:LEU:O | 5:E:1536:LYS:NZ | 2.29 | 0.64 |
| 8:I:104:GLN:NE2 | 8:I:487:GLY:O | 2.30 | 0.64 |
| 5:E:375:LYS:HB3 | 5:E:378:HIS:HD2 | 1.62 | 0.64 |
| 4:D:3086:ALA:HA | 4:D:3089:TYR:HB2 | 1.80 | 0.64 |
| 5:E:1251:ARG:NH2 | 12:N:674:ASP:OD2 | 2.31 | 0.63 |
| 11:M:1656:LEU:HD23 | 11:M:1704:VAL:HG22 | 1.80 | 0.63 |
| 11:M:330:ALA:HB1 | 11:M:354:LEU:HD11 | 1.81 | 0.63 |
| 3:C:275:ASN:HB2 | 3:C:319:TYR:HB2 | 1.80 | 0.63 |
| 1:A:116:ALA:O | 1:A:118:LYS:NZ | 2.32 | 0.63 |
| 11:M:311:PRO:HD3 | 11:M:344:SER:HB2 | 1.81 | 0.63 |
| 1:A:609:ASN:ND2 | 9:J:130:GLN:O | 2.32 | 0.63 |
| 12:N:296:VAL:HA | 12:N:419:VAL:HB | 1.81 | 0.62 |
| 13:Q:146:MET:HE1 | 13:Q:168:GLN:HG3 | 1.81 | 0.62 |
| 4:D:2181:HIS:HB3 | 4:D:2629:GLU:HG3 | 1.82 | 0.62 |
| 16:T:305:VAL:O | 16:T:306:GLY:C | 2.42 | 0.62 |
| 12:N:259:VAL:HG13 | 12:N:260:VAL:HG23 | 1.82 | 0.61 |
| 5:E:232:ASN:HA | 5:E:235:ARG:HG2 | 1.81 | 0.61 |
| 12:N:673:GLN:OE1 | 12:N:677:ARG:NH1 | 2.33 | 0.61 |
| 10:K:1107:GLU:HB2 | 10:K:1124:ALA:HB2 | 1.82 | 0.61 |
| 4:D:2850:ARG:HE | 4:D:3056:ILE:HD12 | 1.64 | 0.61 |
| 15:S:319:GLU:CD | 15:S:319:GLU:H | 2.05 | 0.61 |

*Continued on next page...*

For Manuscript Review

*Continued from previous page...*

| Atom-1 | Atom-2 | Interatomic distance (Å) | Clash overlap (Å) |
|---|---|---|---|
| 16:T:308:PRO:HB3 | 16:T:309:PRO:HD2 | 1.82 | 0.61 |
| 5:E:366:SER:HB3 | 5:E:376:LYS:HE3 | 1.81 | 0.61 |
| 3:C:8:GLU:HG3 | 10:K:2516:PRO:HB3 | 1.83 | 0.61 |
| 1:Z:374:GLN:HE21 | 1:Z:388:LEU:HD22 | 1.66 | 0.61 |
| 5:E:1236:SER:HB2 | 12:N:357:VAL:HA | 1.82 | 0.61 |
| 2:B:714:PRO:HG2 | 14:R:58:THR:HG21 | 1.83 | 0.60 |
| 5:E:1480:PHE:HA | 5:E:1483:LYS:HD2 | 1.84 | 0.60 |
| 12:N:375:ARG:HB3 | 12:N:407:LEU:HD13 | 1.83 | 0.60 |
| 4:D:1599:GLU:OE2 | 6:F:309:ARG:NH1 | 2.34 | 0.60 |
| 5:E:389:LEU:H | 5:E:392:GLU:HB3 | 1.66 | 0.60 |
| 5:E:1299:GLU:OE1 | 5:E:1302:LYS:NZ | 2.35 | 0.59 |
| 4:D:135:GLN:HE22 | 5:E:1679:ASN:HA | 1.67 | 0.59 |
| 4:D:2198:ASP:HB3 | 14:R:180:MET:HE1 | 1.83 | 0.59 |
| 12:N:425:PRO:HB3 | 12:N:438:LEU:HB3 | 1.82 | 0.59 |
| 17:X:65:ARG:NH2 | 17:Y:406:ARG:O | 2.35 | 0.59 |
| 10:K:1526:PRO:O | 10:K:1543:ARG:NH2 | 2.33 | 0.59 |
| 3:C:188:THR:HG22 | 3:C:190:LEU:H | 1.67 | 0.59 |
| 10:K:1531:LEU:HA | 10:K:1535:SER:HB2 | 1.85 | 0.59 |
| 3:C:400:ASP:OD1 | 15:S:261:HIS:NE2 | 2.36 | 0.59 |
| 7:G:738:ASP:OD1 | 7:G:796:ARG:NH2 | 2.34 | 0.59 |
| 5:E:234:ILE:HA | 5:E:237:TRP:HB2 | 1.85 | 0.58 |
| 5:E:1919:GLU:HG2 | 9:J:206:GLU:HG2 | 1.84 | 0.58 |
| 1:A:277:GLU:OE2 | 1:A:560:GLN:NE2 | 2.36 | 0.58 |
| 4:D:2223:ASN:ND2 | 14:R:194:ILE:O | 2.36 | 0.58 |
| 2:B:805:SER:OG | 15:S:187:ASP:OD1 | 2.20 | 0.58 |
| 15:S:318:ASP:HB3 | 15:S:321:GLU:HG2 | 1.84 | 0.58 |
| 2:B:783:ILE:HD11 | 17:Y:42:ILE:HD13 | 1.85 | 0.58 |
| 8:H:179:ARG:NH2 | 17:Y:159:PRO:O | 2.37 | 0.58 |
| 10:K:1344:GLU:N | 10:K:1344:GLU:OE2 | 2.36 | 0.58 |
| 10:K:1402:LEU:HD21 | 10:K:2188:PRO:HG3 | 1.85 | 0.58 |
| 12:N:230:LEU:HD23 | 12:N:234:ALA:HB3 | 1.84 | 0.58 |
| 17:X:313:GLU:OE1 | 17:X:316:ARG:NH2 | 2.37 | 0.58 |
| 1:A:130:PRO:HA | 1:A:156:LEU:O | 2.03 | 0.57 |
| 8:I:105:GLN:HB3 | 8:I:318:LEU:HD12 | 1.87 | 0.57 |
| 12:N:604:ILE:HG23 | 12:N:719:LEU:HD23 | 1.86 | 0.57 |
| 12:N:581:ASP:HB3 | 12:N:592:MET:HE1 | 1.87 | 0.57 |
| 5:E:1289:GLN:HA | 5:E:1292:LEU:HB2 | 1.87 | 0.57 |
| 5:E:1311:LEU:HD21 | 5:E:1353:LEU:HG | 1.87 | 0.57 |
| 11:M:495:ASP:OD1 | 11:M:495:ASP:N | 2.38 | 0.57 |
| 4:D:319:SER:HB3 | 4:D:401:LEU:HD12 | 1.85 | 0.57 |
| 4:D:3115:LEU:O | 9:J:194:ARG:NH1 | 2.38 | 0.57 |

*Continued on next page...*

For Manuscript Review

*Continued from previous page...*

| Atom-1 | Atom-2 | Interatomic distance (Å) | Clash overlap (Å) |
|---|---|---|---|
| 6:F:191:MET:HE2 | 6:F:272:LEU:HD11 | 1.87 | 0.57 |
| 10:K:2269:ARG:HG2 | 10:K:2373:GLU:HG2 | 1.87 | 0.57 |
| 3:C:367:LEU:HD12 | 4:D:240:ARG:HB3 | 1.87 | 0.56 |
| 5:E:358:VAL:HA | 5:E:361:VAL:HB | 1.87 | 0.56 |
| 4:D:2450:LEU:HD22 | 7:G:464:PRO:HG2 | 1.85 | 0.56 |
| 6:F:506:ASP:OD1 | 6:F:507:LEU:N | 2.38 | 0.56 |
| 8:H:248:GLU:O | 8:H:253:ASN:ND2 | 2.39 | 0.56 |
| 14:R:91:ASP:HB2 | 14:R:100:ARG:HH12 | 1.70 | 0.56 |
| 4:D:601:GLN:HG3 | 4:D:602:ASN:H | 1.70 | 0.56 |
| 8:H:118:LEU:HB2 | 8:H:301:VAL:HG21 | 1.86 | 0.56 |
| 8:H:427:PRO:HG3 | 8:H:456:TYR:HE2 | 1.70 | 0.56 |
| 4:D:2807:VAL:HG11 | 4:D:3053:TYR:HE2 | 1.69 | 0.56 |
| 16:T:299:LEU:HG | 16:T:300:SER:N | 2.20 | 0.56 |
| 4:D:2122:GLU:OE1 | 4:D:2122:GLU:N | 2.34 | 0.56 |
| 16:T:303:ARG:NH1 | 16:T:305:VAL:HG22 | 2.20 | 0.56 |
| 5:E:1185:ASN:HD22 | 12:N:216:GLY:HA3 | 1.71 | 0.56 |
| 11:M:1734:ALA:O | 11:M:1738:GLU:N | 2.34 | 0.56 |
| 11:M:1732:ASP:O | 11:M:1735:ALA:HB3 | 2.06 | 0.55 |
| 12:N:357:VAL:HG21 | 12:N:360:LYS:HE2 | 1.88 | 0.55 |
| 12:N:574:GLN:HG2 | 12:N:576:PHE:H | 1.72 | 0.55 |
| 4:D:648:PRO:HD3 | 4:D:2043:GLU:HB2 | 1.87 | 0.55 |
| 12:N:768:GLU:HG2 | 12:N:769:THR:HG23 | 1.86 | 0.55 |
| 16:T:299:LEU:CG | 16:T:300:SER:H | 2.19 | 0.55 |
| 2:B:345:TYR:H | 15:S:126:PRO:HB3 | 1.72 | 0.55 |
| 4:D:1457:VAL:HG22 | 4:D:1527:GLN:HB3 | 1.89 | 0.55 |
| 12:N:705:ILE:HG12 | 12:N:720:LEU:HD22 | 1.89 | 0.55 |
| 6:F:301:THR:OG1 | 6:F:304:ASP:OD2 | 2.25 | 0.55 |
| 7:G:569:PRO:HD3 | 7:G:687:HIS:HE2 | 1.72 | 0.55 |
| 4:D:135:GLN:NE2 | 5:E:1678:ARG:O | 2.39 | 0.55 |
| 5:E:1353:LEU:HD13 | 5:E:1356:ARG:HH21 | 1.71 | 0.55 |
| 5:E:1561:LYS:NZ | 5:E:1600:SER:OG | 2.40 | 0.55 |
| 7:G:618:ALA:HA | 7:G:622:ASP:HB2 | 1.89 | 0.55 |
| 12:N:670:PHE:HD2 | 12:N:673:GLN:HE21 | 1.55 | 0.55 |
| 16:T:331:PRO:HG2 | 16:T:336:ILE:HD11 | 1.87 | 0.55 |
| 1:Z:130:PRO:HD3 | 1:Z:158:GLU:HA | 1.87 | 0.54 |
| 10:K:1103:ALA:HB2 | 10:K:1128:ALA:HB2 | 1.89 | 0.54 |
| 13:Q:150:GLN:NE2 | 13:Q:162:GLU:OE2 | 2.40 | 0.54 |
| 5:E:382:ILE:O | 5:E:385:ASN:HA | 2.07 | 0.54 |
| 1:Z:40:LEU:HD12 | 1:Z:561:LEU:HD23 | 1.88 | 0.54 |
| 12:N:199:GLN:OE1 | 12:N:201:ASN:ND2 | 2.39 | 0.54 |
| 12:N:300:LEU:HD21 | 12:N:427:PRO:HD3 | 1.89 | 0.54 |

*Continued on next page...*

For Manuscript Review

Continued from previous page...

| Atom-1 | Atom-2 | Interatomic distance (Å) | Clash overlap (Å) |
|---|---|---|---|
| 2:B:172:THR:O | 2:B:190:SER:OG | 2.25 | 0.54 |
| 5:E:1575:ARG:NH2 | 5:E:1605:GLU:OE1 | 2.40 | 0.54 |
| 17:X:268:GLU:OE2 | 17:X:268:GLU:N | 2.33 | 0.54 |
| 5:E:1245:ILE:HD12 | 5:E:1248:TYR:HD2 | 1.73 | 0.54 |
| 11:M:479:ILE:HG21 | 11:M:907:ALA:HB1 | 1.90 | 0.54 |
| 1:A:673:LEU:HB3 | 1:A:699:LYS:HB2 | 1.90 | 0.53 |
| 12:N:584:ARG:NH2 | 12:N:726:GLU:OE2 | 2.42 | 0.53 |
| 17:Y:85:LEU:O | 17:Y:90:ARG:NH2 | 2.41 | 0.53 |
| 10:K:901:ARG:NH2 | 10:K:1013:GLU:OE2 | 2.41 | 0.53 |
| 2:B:232:ARG:NH1 | 2:B:594:SER:OG | 2.40 | 0.53 |
| 4:D:2030:LYS:N | 4:D:2030:LYS:HE2 | 2.23 | 0.53 |
| 7:G:616:ALA:O | 7:G:618:ALA:N | 2.42 | 0.53 |
| 11:M:399:LEU:HD13 | 11:M:404:LEU:HD21 | 1.91 | 0.53 |
| 1:A:128:ASN:OD1 | 10:K:2103:ASN:ND2 | 2.40 | 0.53 |
| 1:A:206:LEU:HD22 | 10:K:2104:PRO:HB3 | 1.90 | 0.53 |
| 4:D:484:LEU:HD22 | 4:D:536:PHE:HA | 1.90 | 0.52 |
| 4:D:1566:THR:HG23 | 4:D:1599:GLU:HG2 | 1.91 | 0.52 |
| 4:D:2514:GLU:CD | 4:D:2514:GLU:H | 2.17 | 0.52 |
| 10:K:1603:VAL:O | 10:K:1607:MET:HG3 | 2.09 | 0.52 |
| 5:E:359:TRP:CD1 | 5:E:1498:GLN:HE21 | 2.28 | 0.52 |
| 8:I:466:ASP:N | 8:I:466:ASP:OD1 | 2.42 | 0.52 |
| 11:M:367:LEU:HD22 | 11:M:368:GLY:H | 1.73 | 0.52 |
| 11:M:479:ILE:HD13 | 11:M:899:VAL:HG11 | 1.92 | 0.52 |
| 12:N:200:SER:OG | 12:N:214:HIS:NE2 | 2.41 | 0.52 |
| 15:S:288:LEU:HD12 | 15:S:291:ARG:HH21 | 1.74 | 0.52 |
| 12:N:362:ILE:HD11 | 12:N:742:TRP:HB3 | 1.92 | 0.52 |
| 17:X:381:ASP:OD1 | 17:X:393:ARG:NH1 | 2.42 | 0.52 |
| 4:D:476:ILE:HD11 | 4:D:534:LEU:HD21 | 1.91 | 0.52 |
| 4:D:3092:LYS:HB2 | 5:E:224:ILE:H | 1.74 | 0.52 |
| 10:K:2472:ARG:HH11 | 10:K:2509:VAL:HG11 | 1.75 | 0.52 |
| 1:Z:73:ILE:HG12 | 1:Z:164:MET:HG3 | 1.92 | 0.52 |
| 5:E:383:THR:C | 5:E:385:ASN:H | 2.18 | 0.52 |
| 5:E:1289:GLN:O | 5:E:1293:ASN:N | 2.41 | 0.52 |
| 12:N:756:LEU:HA | 12:N:759:LEU:HB3 | 1.91 | 0.52 |
| 12:N:586:PRO:HA | 12:N:589:LEU:HD12 | 1.92 | 0.52 |
| 5:E:1508:LYS:HA | 5:E:1514:LEU:HA | 1.92 | 0.52 |
| 6:F:223:SER:H | 6:F:226:MET:HE1 | 1.74 | 0.52 |
| 8:I:361:ARG:HD3 | 8:I:523:VAL:HG21 | 1.91 | 0.52 |
| 5:E:1725:ASN:HD22 | 10:K:1715:PRO:HB2 | 1.74 | 0.51 |
| 4:D:1061:ILE:HG13 | 8:H:256:GLU:HB2 | 1.91 | 0.51 |
| 8:H:98:PRO:HD3 | 8:H:314:GLN:HG3 | 1.93 | 0.51 |

Continued on next page...

For Manuscript Review

*Continued from previous page...*

| Atom-1 | Atom-2 | Interatomic distance (Å) | Clash overlap (Å) |
|---|---|---|---|
| 12:N:737:SER:OG | 12:N:738:VAL:N | 2.44 | 0.51 |
| 13:Q:161:MET:HE2 | 13:Q:161:MET:HA | 1.91 | 0.51 |
| 12:N:240:PRO:O | 12:N:268:ARG:NH1 | 2.43 | 0.51 |
| 12:N:607:VAL:HG22 | 12:N:634:ILE:HD12 | 1.92 | 0.51 |
| 5:E:1216:SER:OG | 5:E:1218:ARG:NH1 | 2.40 | 0.51 |
| 12:N:639:ASN:H | 12:N:729:VAL:HG23 | 1.76 | 0.51 |
| 1:A:198:LYS:HD2 | 1:A:238:LEU:HD12 | 1.92 | 0.51 |
| 11:M:356:LYS:NZ | 11:M:425:GLU:O | 2.43 | 0.51 |
| 4:D:910:ILE:HG13 | 4:D:913:LYS:HE2 | 1.93 | 0.51 |
| 5:E:1150:PHE:H | 12:N:444:ASP:HB3 | 1.75 | 0.51 |
| 5:E:1520:ASP:HA | 5:E:1523:LYS:HG2 | 1.91 | 0.51 |
| 8:I:90:ASP:OD2 | 8:I:92:GLN:NE2 | 2.39 | 0.51 |
| 8:I:332:GLY:HA2 | 8:I:358:MET:HE3 | 1.92 | 0.51 |
| 7:G:235:MET:HE3 | 7:G:411:PHE:HB2 | 1.93 | 0.51 |
| 5:E:387:GLU:HB3 | 5:E:1220:LEU:HD23 | 1.92 | 0.51 |
| 10:K:1089:LEU:HD23 | 10:K:1159:LEU:HD22 | 1.93 | 0.51 |
| 10:K:2328:LEU:HD13 | 10:K:2334:VAL:HG22 | 1.93 | 0.51 |
| 11:M:926:TRP:CD1 | 11:M:931:GLY:HA3 | 2.46 | 0.51 |
| 12:N:502:GLU:HB2 | 12:N:514:ILE:HG22 | 1.93 | 0.51 |
| 3:C:130:ASP:HB2 | 3:C:134:SER:HB2 | 1.93 | 0.51 |
| 4:D:2323:TRP:O | 4:D:2330:GLN:NE2 | 2.40 | 0.51 |
| 10:K:1613:GLU:OE1 | 10:K:1630:ARG:NH2 | 2.44 | 0.51 |
| 1:Z:121:TYR:HB2 | 1:Z:168:ILE:HB | 1.91 | 0.51 |
| 5:E:217:LYS:NZ | 5:E:1666:GLU:OE2 | 2.43 | 0.50 |
| 10:K:2147:LEU:O | 10:K:2152:ARG:NH2 | 2.44 | 0.50 |
| 12:N:200:SER:HG | 12:N:214:HIS:HE2 | 1.59 | 0.50 |
| 16:T:304:LEU:CD1 | 16:T:324:LEU:HD23 | 2.40 | 0.50 |
| 2:B:498:ASP:O | 2:B:504:ASN:ND2 | 2.42 | 0.50 |
| 12:N:237:VAL:HG21 | 12:N:266:LEU:HD22 | 1.93 | 0.50 |
| 12:N:688:LYS:HB2 | 12:N:776:ILE:HG12 | 1.93 | 0.50 |
| 8:H:174:ASP:OD1 | 8:H:174:ASP:N | 2.41 | 0.50 |
| 16:T:305:VAL:O | 16:T:306:GLY:O | 2.29 | 0.50 |
| 5:E:1238:GLY:O | 12:N:353:ARG:NH2 | 2.45 | 0.50 |
| 9:J:248:GLN:OE1 | 9:J:248:GLN:N | 2.36 | 0.50 |
| 11:M:899:VAL:HA | 11:M:902:MET:HG3 | 1.94 | 0.50 |
| 8:H:243:ASP:OD1 | 8:H:243:ASP:N | 2.44 | 0.50 |
| 11:M:482:LEU:HG | 11:M:900:LEU:HD21 | 1.94 | 0.50 |
| 15:S:108:ASN:ND2 | 15:S:112:GLU:OE2 | 2.37 | 0.50 |
| 2:B:769:MET:HE3 | 3:C:333:ALA:HB1 | 1.92 | 0.50 |
| 5:E:234:ILE:HD13 | 5:E:1432:PRO:HG3 | 1.94 | 0.50 |
| 5:E:388:THR:OG1 | 5:E:1221:TRP:O | 2.29 | 0.50 |

*Continued on next page...*

For Manuscript Review

*Continued from previous page...*

| Atom-1 | Atom-2 | Interatomic distance (Å) | Clash overlap (Å) |
|---|---|---|---|
| 10:K:1086:PRO:HB2 | 10:K:1163:PRO:HG3 | 1.94 | 0.50 |
| 7:G:472:GLY:O | 7:G:474:GLY:N | 2.45 | 0.49 |
| 16:T:305:VAL:HG23 | 16:T:323:THR:HG21 | 1.94 | 0.49 |
| 5:E:499:TYR:HE1 | 12:N:204:LEU:HD21 | 1.78 | 0.49 |
| 13:Q:94:PHE:CZ | 13:Q:186:MET:HB2 | 2.47 | 0.49 |
| 1:A:183:ASP:OD1 | 1:A:183:ASP:N | 2.45 | 0.49 |
| 5:E:1359:LEU:O | 5:E:1363:ILE:HG12 | 2.12 | 0.49 |
| 7:G:577:ARG:NH1 | 7:G:805:GLN:OE1 | 2.46 | 0.49 |
| 15:S:83:SER:OG | 15:S:98:ASP:OD1 | 2.30 | 0.49 |
| 16:T:192:GLU:OE1 | 16:T:193:LEU:N | 2.46 | 0.49 |
| 1:Z:186:THR:HG22 | 1:Z:190:ARG:HD2 | 1.94 | 0.49 |
| 12:N:438:LEU:HD22 | 12:N:587:GLU:HG2 | 1.95 | 0.49 |
| 12:N:481:LEU:HG | 12:N:484:ALA:HB3 | 1.94 | 0.49 |
| 1:Z:676:PRO:HD2 | 1:Z:679:ILE:HD13 | 1.95 | 0.49 |
| 4:D:302:ALA:HB2 | 4:D:436:THR:HG22 | 1.94 | 0.49 |
| 12:N:402:ALA:O | 12:N:406:MET:HE3 | 2.13 | 0.49 |
| 1:Z:19:PHE:HA | 1:Z:42:PRO:HD2 | 1.94 | 0.49 |
| 4:D:468:ASP:OD1 | 4:D:468:ASP:N | 2.40 | 0.49 |
| 2:B:597:MET:HE2 | 2:B:599:GLN:HG2 | 1.95 | 0.48 |
| 4:D:859:MET:HE3 | 4:D:859:MET:HB3 | 1.76 | 0.48 |
| 4:D:963:ASN:HD21 | 4:D:966:TYR:H | 1.60 | 0.48 |
| 8:I:502:ASP:OD1 | 8:I:503:HIS:ND1 | 2.45 | 0.48 |
| 10:K:1616:GLU:OE2 | 1:Z:574:ARG:NH2 | 2.46 | 0.48 |
| 12:N:213:HIS:HD2 | 12:N:226:VAL:HG23 | 1.77 | 0.48 |
| 7:G:724:ILE:HG21 | 7:G:807:GLN:HB2 | 1.94 | 0.48 |
| 10:K:2057:GLY:HA2 | 10:K:2060:GLU:HG3 | 1.95 | 0.48 |
| 12:N:284:TYR:HB3 | 12:N:386:ALA:HB2 | 1.95 | 0.48 |
| 13:Q:166:VAL:HG23 | 13:Q:184:TRP:HD1 | 1.79 | 0.48 |
| 1:A:553:LYS:NZ | 1:Z:169:ASN:OD1 | 2.45 | 0.48 |
| 4:D:1967:LYS:HE3 | 4:D:2041:PRO:HA | 1.94 | 0.48 |
| 10:K:2023:ARG:HG2 | 10:K:2126:VAL:HG21 | 1.95 | 0.48 |
| 12:N:335:LEU:HD21 | 12:N:339:GLY:HA2 | 1.95 | 0.48 |
| 3:C:215:ILE:HG23 | 3:C:229:VAL:HG22 | 1.95 | 0.48 |
| 4:D:3092:LYS:HB3 | 5:E:223:LEU:HD12 | 1.95 | 0.48 |
| 3:C:338:ASN:OD1 | 3:C:338:ASN:N | 2.42 | 0.48 |
| 4:D:342:VAL:HG22 | 4:D:2827:ARG:HH12 | 1.78 | 0.48 |
| 4:D:1971:SER:O | 4:D:1971:SER:OG | 2.25 | 0.48 |
| 8:H:78:THR:HG22 | 8:H:534:ALA:HA | 1.95 | 0.48 |
| 9:J:195:ARG:O | 9:J:199:MET:HG3 | 2.14 | 0.48 |
| 12:N:691:ARG:HH12 | 12:N:776:ILE:H | 1.61 | 0.48 |
| 10:K:2299:VAL:HG23 | 10:K:2300:MET:HG3 | 1.95 | 0.48 |

*Continued on next page...*

For Manuscript Review

*Continued from previous page...*

| Atom-1 | Atom-2 | Interatomic distance (Å) | Clash overlap (Å) |
|---|---|---|---|
| 12:N:356:SER:HB2 | 12:N:362:ILE:HG12 | 1.95 | 0.48 |
| 12:N:635:ILE:HD12 | 12:N:697:VAL:HG22 | 1.95 | 0.48 |
| 2:B:727:THR:OG1 | 2:B:728:LYS:N | 2.45 | 0.48 |
| 3:C:558:ALA:HA | 5:E:1518:LEU:HD23 | 1.95 | 0.48 |
| 7:G:784:ILE:HG23 | 7:G:789:GLN:HB2 | 1.95 | 0.48 |
| 10:K:1428:TYR:O | 10:K:1431:ASP:HB2 | 2.14 | 0.48 |
| 8:H:453:ASP:OD1 | 8:H:453:ASP:N | 2.35 | 0.47 |
| 10:K:1445:MET:HE2 | 10:K:1445:MET:HB2 | 1.79 | 0.47 |
| 17:Y:266:THR:HG22 | 17:Y:272:VAL:HA | 1.96 | 0.47 |
| 1:Z:120:THR:HG23 | 1:Z:169:ASN:HD21 | 1.79 | 0.47 |
| 10:K:1837:THR:HG21 | 10:K:1891:LEU:HB2 | 1.96 | 0.47 |
| 12:N:313:LEU:HD21 | 12:N:547:LEU:HD23 | 1.96 | 0.47 |
| 12:N:419:VAL:HA | 12:N:466:VAL:HG22 | 1.96 | 0.47 |
| 4:D:485:PHE:HB3 | 4:D:535:LEU:HB2 | 1.95 | 0.47 |
| 5:E:500:LYS:O | 5:E:504:LYS:HG2 | 2.14 | 0.47 |
| 11:M:461:ARG:HB2 | 11:M:464:TYR:HD2 | 1.78 | 0.47 |
| 1:Z:271:LYS:HB3 | 1:Z:566:TRP:HB2 | 1.94 | 0.47 |
| 4:D:111:PRO:HG3 | 16:T:252:MET:HE1 | 1.96 | 0.47 |
| 4:D:394:LEU:HD23 | 4:D:2843:LYS:HB2 | 1.95 | 0.47 |
| 4:D:1402:LYS:HE2 | 4:D:1402:LYS:HB2 | 1.73 | 0.47 |
| 16:T:298:ALA:O | 16:T:299:LEU:C | 2.57 | 0.47 |
| 8:H:232:SER:OG | 8:H:233:GLU:N | 2.47 | 0.47 |
| 8:I:326:ASP:N | 8:I:326:ASP:OD1 | 2.46 | 0.47 |
| 10:K:1338:GLU:N | 10:K:1338:GLU:OE2 | 2.45 | 0.47 |
| 12:N:634:ILE:HG21 | 12:N:704:ALA:HB1 | 1.97 | 0.47 |
| 2:B:222:ARG:HG2 | 2:B:797:GLY:HA3 | 1.97 | 0.47 |
| 4:D:777:LYS:HZ2 | 4:D:779:LYS:HA | 1.79 | 0.47 |
| 4:D:366:ILE:HD13 | 4:D:381:ILE:HG22 | 1.96 | 0.47 |
| 4:D:604:LEU:HD23 | 4:D:604:LEU:HA | 1.83 | 0.47 |
| 4:D:3008:ASP:OD1 | 4:D:3009:ILE:N | 2.48 | 0.47 |
| 5:E:1233:TYR:OH | 12:N:706:ARG:O | 2.29 | 0.47 |
| 6:F:179:ASP:OD2 | 17:X:80:ARG:NH1 | 2.47 | 0.47 |
| 6:F:519:PRO:HD3 | 6:F:559:GLU:HG3 | 1.97 | 0.47 |
| 5:E:1363:ILE:HG21 | 5:E:1424:LEU:HD22 | 1.96 | 0.47 |
| 12:N:454:SER:HB3 | 12:N:480:ARG:HH11 | 1.80 | 0.47 |
| 1:Z:378:PHE:HA | 1:Z:383:LEU:HB2 | 1.96 | 0.47 |
| 3:C:473:ARG:O | 3:C:473:ARG:NE | 2.39 | 0.47 |
| 6:F:290:TYR:HE1 | 6:F:446:GLN:HB3 | 1.78 | 0.47 |
| 6:F:274:ASP:OD1 | 6:F:274:ASP:N | 2.47 | 0.47 |
| 8:I:118:LEU:HB2 | 8:I:301:VAL:HG21 | 1.96 | 0.47 |
| 2:B:456:ARG:HD3 | 2:B:471:ILE:O | 2.15 | 0.46 |

*Continued on next page...*

For Manuscript Review

*Continued from previous page...*

| Atom-1 | Atom-2 | Interatomic distance (Å) | Clash overlap (Å) |
|---|---|---|---|
| 12:N:207:PRO:HA | 12:N:233:GLY:HA3 | 1.97 | 0.46 |
| 16:T:291:ASP:OD1 | 16:T:293:LYS:N | 2.48 | 0.46 |
| 2:B:440:PHE:HD1 | 2:B:444:MET:HE2 | 1.80 | 0.46 |
| 14:R:153:HIS:NE2 | 14:R:199:ASP:OD1 | 2.42 | 0.46 |
| 2:B:761:ASN:O | 2:B:765:MET:HG3 | 2.15 | 0.46 |
| 8:H:466:ASP:OD1 | 8:H:466:ASP:N | 2.46 | 0.46 |
| 13:Q:121:TRP:HA | 13:Q:127:LEU:HD21 | 1.97 | 0.46 |
| 13:Q:215:LEU:HD22 | 13:Q:229:VAL:HG21 | 1.98 | 0.46 |
| 2:B:325:LYS:HG3 | 15:S:137:VAL:HG11 | 1.97 | 0.46 |
| 2:B:704:ASP:HA | 3:C:402:MET:HE3 | 1.96 | 0.46 |
| 5:E:1524:GLY:O | 5:E:1528:ARG:N | 2.48 | 0.46 |
| 12:N:210:VAL:HG22 | 12:N:236:LEU:HB2 | 1.98 | 0.46 |
| 17:Y:45:LEU:HD23 | 17:Y:45:LEU:H | 1.80 | 0.46 |
| 5:E:355:THR:HB | 5:E:1493:VAL:HG11 | 1.98 | 0.46 |
| 8:H:261:THR:OG1 | 8:H:262:THR:N | 2.49 | 0.46 |
| 4:D:327:LYS:HG2 | 11:M:706:PHE:HB3 | 1.96 | 0.46 |
| 8:H:429:GLN:O | 8:H:433:MET:HG3 | 2.16 | 0.46 |
| 12:N:222:GLN:HG3 | 12:N:223:VAL:HG23 | 1.97 | 0.46 |
| 5:E:384:TYR:HE2 | 5:E:1504:VAL:HG23 | 1.81 | 0.46 |
| 16:T:308:PRO:CB | 16:T:309:PRO:HD2 | 2.44 | 0.46 |
| 1:Z:291:LEU:HD23 | 1:Z:291:LEU:H | 1.81 | 0.46 |
| 3:C:326:MET:HE1 | 3:C:332:MET:HE3 | 1.98 | 0.46 |
| 3:C:563:ILE:HG21 | 3:C:566:PHE:HE1 | 1.81 | 0.46 |
| 4:D:1902:LEU:HB2 | 4:D:1953:ASN:HB3 | 1.97 | 0.46 |
| 5:E:1347:CYS:HA | 5:E:1350:ARG:HE | 1.81 | 0.46 |
| 5:E:1592:ILE:HA | 5:E:1599:VAL:HG21 | 1.97 | 0.46 |
| 10:K:1611:ASP:O | 10:K:1615:MET:HG3 | 2.15 | 0.46 |
| 12:N:175:GLY:HA3 | 12:N:183:ALA:HB2 | 1.96 | 0.46 |
| 1:Z:296:HIS:O | 1:Z:296:HIS:ND1 | 2.45 | 0.46 |
| 5:E:360:PHE:CD1 | 5:E:1426:MET:HE1 | 2.51 | 0.46 |
| 16:T:287:VAL:HG12 | 16:T:300:SER:HB2 | 1.98 | 0.46 |
| 4:D:242:ASN:ND2 | 4:D:244:SER:OG | 2.50 | 0.46 |
| 9:J:234:PHE:O | 9:J:238:VAL:HG12 | 2.16 | 0.46 |
| 15:S:318:ASP:O | 15:S:319:GLU:C | 2.59 | 0.46 |
| 9:J:211:ALA:O | 9:J:214:GLN:NE2 | 2.45 | 0.45 |
| 11:M:408:LEU:O | 11:M:412:ARG:HG2 | 2.15 | 0.45 |
| 12:N:471:ASP:HB3 | 12:N:474:ALA:HB2 | 1.98 | 0.45 |
| 12:N:505:LYS:HD3 | 12:N:512:GLU:HG2 | 1.96 | 0.45 |
| 12:N:689:ALA:O | 12:N:693:ASN:ND2 | 2.48 | 0.45 |
| 3:C:194:GLN:HA | 3:C:312:GLN:HE22 | 1.82 | 0.45 |
| 5:E:1520:ASP:OD1 | 5:E:1523:LYS:NZ | 2.48 | 0.45 |

*Continued on next page...*

For Manuscript Review

*Continued from previous page...*

| Atom-1 | Atom-2 | Interatomic distance (Å) | Clash overlap (Å) |
|---|---|---|---|
| 4:D:308:ARG:HB3 | 4:D:3060:ILE:HG21 | 1.98 | 0.45 |
| 4:D:541:GLU:HG3 | 14:R:130:TRP:HB2 | 1.99 | 0.45 |
| 4:D:2808:LYS:HG2 | 11:M:706:PHE:CZ | 2.51 | 0.45 |
| 5:E:233:ARG:O | 5:E:236:GLN:HG2 | 2.15 | 0.45 |
| 12:N:776:ILE:HG13 | 12:N:783:LEU:HD13 | 1.98 | 0.45 |
| 3:C:103:ARG:NH1 | 3:C:107:ASP:OD1 | 2.42 | 0.45 |
| 3:C:436:LYS:HB3 | 3:C:451:THR:HB | 1.98 | 0.45 |
| 5:E:1205:LYS:HE2 | 5:E:1207:LEU:HD21 | 1.98 | 0.45 |
| 5:E:1480:PHE:O | 5:E:1484:TYR:HD1 | 1.98 | 0.45 |
| 8:H:251:LEU:HD12 | 8:H:510:VAL:HG12 | 1.98 | 0.45 |
| 13:Q:226:VAL:O | 13:Q:230:GLN:HG3 | 2.17 | 0.45 |
| 1:A:604:ASP:OD1 | 1:A:604:ASP:N | 2.49 | 0.45 |
| 12:N:613:ASN:HD21 | 12:N:642:LEU:HB2 | 1.82 | 0.45 |
| 12:N:622:MET:O | 12:N:626:ALA:N | 2.48 | 0.45 |
| 16:T:305:VAL:HG23 | 16:T:323:THR:CG2 | 2.46 | 0.45 |
| 17:X:222:ALA:HB3 | 17:X:355:ALA:HB2 | 1.96 | 0.45 |
| 2:B:605:GLU:OE1 | 2:B:798:HIS:NE2 | 2.50 | 0.45 |
| 5:E:382:ILE:O | 5:E:385:ASN:CA | 2.65 | 0.45 |
| 10:K:1189:VAL:HG11 | 10:K:1291:PRO:HD3 | 1.99 | 0.45 |
| 7:G:686:GLU:OE2 | 7:G:689:ARG:NH2 | 2.39 | 0.45 |
| 12:N:174:PHE:CD2 | 12:N:182:ALA:HB1 | 2.52 | 0.45 |
| 12:N:406:MET:HE2 | 12:N:406:MET:HB3 | 1.82 | 0.45 |
| 1:A:236:ARG:HA | 1:A:236:ARG:HD3 | 1.83 | 0.45 |
| 10:K:1708:ASP:O | 10:K:1711:SER:OG | 2.35 | 0.45 |
| 10:K:2514:PRO:HA | 10:K:2515:PRO:HD3 | 1.92 | 0.45 |
| 4:D:3058:LEU:HD22 | 4:D:3062:ARG:HB2 | 1.99 | 0.44 |
| 5:E:1695:GLN:H | 5:E:1695:GLN:HG3 | 1.44 | 0.44 |
| 10:K:994:ALA:HB2 | 10:K:1003:LEU:HD12 | 1.99 | 0.44 |
| 10:K:1862:VAL:HG22 | 10:K:1873:VAL:HG22 | 2.00 | 0.44 |
| 12:N:338:ASP:OD1 | 12:N:380:ASN:ND2 | 2.50 | 0.44 |
| 12:N:420:VAL:HG21 | 12:N:477:LEU:HD21 | 1.99 | 0.44 |
| 16:T:307:GLN:CB | 16:T:308:PRO:CD | 2.88 | 0.44 |
| 4:D:1402:LYS:O | 15:S:357:ARG:NH1 | 2.51 | 0.44 |
| 5:E:499:TYR:CZ | 12:N:338:ASP:HB2 | 2.52 | 0.44 |
| 5:E:1833:LEU:HB3 | 5:E:1848:VAL:HG12 | 1.98 | 0.44 |
| 12:N:200:SER:HA | 12:N:212:VAL:HB | 2.00 | 0.44 |
| 15:S:172:MET:HE3 | 15:S:172:MET:HB3 | 1.86 | 0.44 |
| 2:B:233:VAL:HG23 | 2:B:597:MET:HG3 | 1.99 | 0.44 |
| 4:D:3019:LEU:HD22 | 4:D:3048:ILE:HG21 | 1.99 | 0.44 |
| 5:E:1526:GLN:HA | 5:E:1529:PHE:CD2 | 2.53 | 0.44 |
| 5:E:1768:LEU:HD12 | 16:T:313:PRO:HB2 | 2.00 | 0.44 |

*Continued on next page...*

For Manuscript Review

*Continued from previous page...*

| Atom-1 | Atom-2 | Interatomic distance (Å) | Clash overlap (Å) |
|---|---|---|---|
| 8:I:429:GLN:O | 8:I:433:MET:HG3 | 2.18 | 0.44 |
| 11:M:362:LEU:HD11 | 11:M:437:VAL:HG22 | 1.99 | 0.44 |
| 13:Q:123:ASP:HB3 | 13:Q:126:ARG:HG3 | 2.00 | 0.44 |
| 15:S:334:GLU:HA | 15:S:337:ILE:HG22 | 1.99 | 0.44 |
| 17:Y:243:ASN:O | 17:Y:245:HIS:ND1 | 2.51 | 0.44 |
| 2:B:809:MET:HG3 | 2:B:819:TYR:CD1 | 2.53 | 0.44 |
| 3:C:529:GLN:HB2 | 3:C:530:ASN:H | 1.63 | 0.44 |
| 4:D:1878:SER:OG | 4:D:1879:TYR:N | 2.50 | 0.44 |
| 11:M:925:LEU:O | 11:M:929:VAL:HG23 | 2.17 | 0.44 |
| 3:C:130:ASP:OD1 | 3:C:130:ASP:N | 2.49 | 0.44 |
| 3:C:150:MET:HE3 | 3:C:150:MET:HB2 | 1.80 | 0.44 |
| 4:D:2314:LYS:HE3 | 4:D:2314:LYS:HB2 | 1.67 | 0.44 |
| 5:E:362:LYS:HD2 | 5:E:1501:LYS:HD2 | 1.99 | 0.44 |
| 9:J:242:ILE:O | 9:J:246:VAL:HG22 | 2.17 | 0.44 |
| 4:D:1811:ASN:C | 4:D:1811:ASN:HD22 | 2.26 | 0.44 |
| 11:M:896:LEU:HD21 | 11:M:918:VAL:HG13 | 1.99 | 0.44 |
| 8:H:251:LEU:HD21 | 8:H:511:GLU:HG2 | 2.00 | 0.44 |
| 10:K:1098:ILE:HD11 | 10:K:2096:ARG:HB3 | 2.00 | 0.44 |
| 11:M:922:MET:HE2 | 11:M:922:MET:HA | 2.00 | 0.44 |
| 1:Z:44:ASP:OD1 | 1:Z:44:ASP:N | 2.49 | 0.44 |
| 4:D:851:ILE:HG22 | 4:D:1950:PRO:HB2 | 1.99 | 0.44 |
| 4:D:910:ILE:HB | 4:D:913:LYS:HB3 | 1.99 | 0.44 |
| 9:J:190:TYR:O | 9:J:194:ARG:HG2 | 2.18 | 0.44 |
| 10:K:1501:MET:HG2 | 10:K:1562:TRP:CD2 | 2.53 | 0.44 |
| 3:C:559:LEU:HD13 | 5:E:228:LEU:O | 2.17 | 0.43 |
| 5:E:1203:GLN:OE1 | 5:E:1250:SER:OG | 2.35 | 0.43 |
| 5:E:1225:LYS:HD2 | 5:E:1225:LYS:HA | 1.86 | 0.43 |
| 10:K:1353:VAL:HG13 | 10:K:1356:ARG:HH12 | 1.82 | 0.43 |
| 13:Q:110:VAL:HG13 | 13:Q:245:LYS:HG3 | 1.99 | 0.43 |
| 4:D:2123:LYS:H | 4:D:2123:LYS:HG2 | 1.55 | 0.43 |
| 12:N:221:ASP:N | 12:N:221:ASP:OD1 | 2.51 | 0.43 |
| 17:X:407:ASP:OD1 | 17:X:407:ASP:N | 2.50 | 0.43 |
| 17:Y:243:ASN:N | 17:Y:243:ASN:OD1 | 2.50 | 0.43 |
| 5:E:1607:MET:HB3 | 5:E:1630:PRO:HG2 | 1.99 | 0.43 |
| 8:I:505:LEU:HD12 | 8:I:505:LEU:HA | 1.84 | 0.43 |
| 11:M:311:PRO:HG2 | 11:M:347:ARG:HH21 | 1.82 | 0.43 |
| 12:N:756:LEU:HG | 12:N:759:LEU:HD23 | 2.00 | 0.43 |
| 3:C:267:LYS:HD2 | 3:C:267:LYS:HA | 1.75 | 0.43 |
| 4:D:777:LYS:HB3 | 4:D:777:LYS:HE3 | 1.80 | 0.43 |
| 5:E:372:LEU:HD23 | 5:E:372:LEU:HA | 1.79 | 0.43 |
| 8:H:324:ARG:NH2 | 8:H:326:ASP:OD2 | 2.51 | 0.43 |

*Continued on next page...*

For Manuscript Review

*Continued from previous page...*

| Atom-1 | Atom-2 | Interatomic distance (Å) | Clash overlap (Å) |
|---|---|---|---|
| 1:A:271:LYS:HG3 | 1:A:566:TRP:CE3 | 2.54 | 0.43 |
| 4:D:230:PRO:O | 4:D:234:MET:HG3 | 2.19 | 0.43 |
| 4:D:2704:LEU:HD22 | 4:D:3057:VAL:HG13 | 2.00 | 0.43 |
| 5:E:1228:GLN:HA | 5:E:1231:ALA:HB3 | 2.00 | 0.43 |
| 8:I:99:MET:HE2 | 8:I:99:MET:HB3 | 1.86 | 0.43 |
| 12:N:330:ILE:HD11 | 12:N:335:LEU:HD22 | 2.00 | 0.43 |
| 3:C:559:LEU:HD12 | 3:C:560:CYS:HA | 2.01 | 0.43 |
| 4:D:2827:ARG:O | 4:D:2827:ARG:HD3 | 2.19 | 0.43 |
| 5:E:1140:ILE:HD13 | 12:N:737:SER:HA | 2.00 | 0.43 |
| 5:E:1616:ARG:HD2 | 5:E:1650:ALA:HB2 | 2.00 | 0.43 |
| 11:M:1698:MET:SD | 11:M:1698:MET:N | 2.91 | 0.43 |
| 12:N:200:SER:HB2 | 12:N:379:LEU:HD13 | 2.00 | 0.43 |
| 1:Z:78:HIS:CE1 | 1:Z:80:PHE:HB2 | 2.54 | 0.43 |
| 3:C:182:GLU:HG2 | 3:C:317:HIS:CD2 | 2.54 | 0.43 |
| 4:D:3048:ILE:HD12 | 4:D:3048:ILE:HA | 1.87 | 0.43 |
| 5:E:1500:GLY:N | 5:E:1506:PRO:HB3 | 2.34 | 0.43 |
| 12:N:380:ASN:O | 12:N:384:VAL:HG23 | 2.19 | 0.43 |
| 12:N:605:PHE:HB2 | 12:N:718:VAL:HA | 2.01 | 0.43 |
| 1:A:107:ILE:HG22 | 1:A:108:SER:H | 1.83 | 0.43 |
| 2:B:287:LYS:H | 2:B:287:LYS:HG2 | 1.60 | 0.43 |
| 5:E:1209:VAL:HA | 12:N:680:LEU:HD13 | 1.99 | 0.43 |
| 10:K:1452:ASP:O | 10:K:1454:GLY:N | 2.47 | 0.43 |
| 12:N:244:LYS:HB3 | 12:N:244:LYS:HE3 | 1.66 | 0.43 |
| 12:N:613:ASN:ND2 | 12:N:642:LEU:HB2 | 2.34 | 0.43 |
| 16:T:314:VAL:HG21 | 16:T:330:LEU:HD11 | 1.99 | 0.43 |
| 5:E:496:LYS:HB3 | 5:E:496:LYS:HE2 | 1.81 | 0.43 |
| 13:Q:116:VAL:O | 13:Q:120:ILE:HG13 | 2.19 | 0.43 |
| 17:X:228:VAL:HG12 | 17:X:230:GLN:H | 1.83 | 0.43 |
| 17:Y:117:ASP:HB3 | 17:Y:120:VAL:HG23 | 2.01 | 0.43 |
| 17:Y:222:ALA:HB3 | 17:Y:355:ALA:HB2 | 2.00 | 0.43 |
| 4:D:769:LYS:O | 4:D:773:GLN:NE2 | 2.50 | 0.43 |
| 5:E:1509:ASP:N | 5:E:1513:ARG:O | 2.42 | 0.43 |
| 10:K:975:VAL:HG11 | 10:K:1029:LEU:HD23 | 2.00 | 0.43 |
| 10:K:1878:TRP:HB2 | 10:K:1879:GLY:H | 1.45 | 0.43 |
| 11:M:923:LEU:HD11 | 11:M:933:PRO:HG3 | 2.01 | 0.43 |
| 12:N:423:LEU:HB3 | 12:N:446:VAL:HG22 | 2.01 | 0.43 |
| 1:Z:44:ASP:O | 1:Z:48:SER:OG | 2.29 | 0.43 |
| 4:D:478:THR:HG21 | 4:D:484:LEU:HD12 | 2.00 | 0.42 |
| 5:E:384:TYR:O | 5:E:1487:ARG:NH2 | 2.52 | 0.42 |
| 5:E:496:LYS:O | 5:E:500:LYS:HG2 | 2.19 | 0.42 |
| 5:E:1478:SER:HB2 | 5:E:1479:SER:H | 1.61 | 0.42 |

*Continued on next page...*

For Manuscript Review

*Continued from previous page...*

| Atom-1 | Atom-2 | Interatomic distance (Å) | Clash overlap (Å) |
|---|---|---|---|
| 4:D:410:SER:HB2 | 4:D:417:VAL:HG22 | 2.00 | 0.42 |
| 6:F:155:GLY:HA3 | 6:F:168:THR:HG22 | 2.01 | 0.42 |
| 12:N:289:ARG:HA | 12:N:414:SER:HB2 | 2.01 | 0.42 |
| 17:X:141:ASP:N | 17:X:141:ASP:OD1 | 2.52 | 0.42 |
| 5:E:357:HIS:CE1 | 5:E:1497:ILE:HD13 | 2.55 | 0.42 |
| 5:E:1888:LYS:O | 5:E:1892:GLU:HG2 | 2.20 | 0.42 |
| 7:G:117:ASP:OD1 | 7:G:117:ASP:N | 2.52 | 0.42 |
| 12:N:729:VAL:O | 12:N:740:THR:HA | 2.19 | 0.42 |
| 16:T:271:ALA:O | 16:T:280:ALA:HA | 2.19 | 0.42 |
| 4:D:488:LYS:HG3 | 14:R:114:VAL:HG12 | 2.00 | 0.42 |
| 4:D:1045:VAL:HG13 | 4:D:1178:ILE:HD12 | 2.01 | 0.42 |
| 7:G:73:ARG:HA | 7:G:73:ARG:HD3 | 1.86 | 0.42 |
| 7:G:422:THR:HG23 | 7:G:492:THR:HG22 | 2.00 | 0.42 |
| 11:M:485:LEU:HD22 | 11:M:929:VAL:HG21 | 2.01 | 0.42 |
| 11:M:906:ASP:HB3 | 11:M:909:ARG:HH21 | 1.85 | 0.42 |
| 1:A:11:MET:HA | 1:A:11:MET:HE2 | 2.01 | 0.42 |
| 1:A:48:SER:OG | 1:A:559:SER:OG | 2.33 | 0.42 |
| 4:D:2223:ASN:HA | 4:D:2226:ILE:HG12 | 2.01 | 0.42 |
| 4:D:2571:ILE:HD12 | 4:D:2571:ILE:HA | 1.83 | 0.42 |
| 4:D:2703:LEU:HD12 | 4:D:2703:LEU:H | 1.84 | 0.42 |
| 5:E:1362:LYS:HD3 | 5:E:1362:LYS:HA | 1.87 | 0.42 |
| 7:G:242:LEU:HD21 | 7:G:251:TRP:CD2 | 2.55 | 0.42 |
| 8:I:473:MET:HE3 | 8:I:473:MET:HB2 | 1.79 | 0.42 |
| 12:N:411:ASP:C | 12:N:456:ARG:HH22 | 2.28 | 0.42 |
| 12:N:596:LEU:HD13 | 12:N:719:LEU:HG | 2.01 | 0.42 |
| 12:N:727:ASP:HB2 | 12:N:742:TRP:CD1 | 2.55 | 0.42 |
| 14:R:106:ASP:OD1 | 14:R:106:ASP:N | 2.51 | 0.42 |
| 16:T:294:SER:HB3 | 16:T:296:THR:HG23 | 2.01 | 0.42 |
| 5:E:1153:LEU:HD11 | 12:N:406:MET:HB2 | 2.02 | 0.42 |
| 12:N:501:ALA:HA | 12:N:515:VAL:HG22 | 2.01 | 0.42 |
| 17:Y:160:PHE:HB3 | 17:Y:164:SER:HB2 | 2.01 | 0.42 |
| 5:E:230:SER:HB3 | 5:E:1425:ASN:HD21 | 1.85 | 0.42 |
| 6:F:167:ARG:HH12 | 6:F:184:GLN:HG2 | 1.83 | 0.42 |
| 9:J:241:ASN:OD1 | 9:J:241:ASN:N | 2.52 | 0.42 |
| 10:K:1607:MET:HE1 | 10:K:1628:TYR:HA | 2.02 | 0.42 |
| 17:Y:42:ILE:H | 17:Y:42:ILE:HG13 | 1.73 | 0.42 |
| 3:C:6:LEU:HD22 | 10:K:2518:GLY:HA3 | 2.02 | 0.42 |
| 4:D:1996:LEU:HD23 | 4:D:1996:LEU:HA | 1.87 | 0.42 |
| 5:E:214:SER:OG | 5:E:215:TYR:N | 2.53 | 0.42 |
| 5:E:1876:THR:OG1 | 5:E:1877:THR:N | 2.53 | 0.42 |
| 10:K:1607:MET:HE1 | 10:K:1628:TYR:HD1 | 1.84 | 0.42 |

*Continued on next page...*

For Manuscript Review

*Continued from previous page...*

| Atom-1 | Atom-2 | Interatomic distance (Å) | Clash overlap (Å) |
|---|---|---|---|
| 12:N:173:LEU:HD11 | 12:N:195:VAL:HB | 2.02 | 0.42 |
| 2:B:762:ARG:HD3 | 2:B:762:ARG:HA | 1.86 | 0.42 |
| 4:D:330:PHE:CD1 | 4:D:387:ARG:HG3 | 2.55 | 0.42 |
| 4:D:407:TYR:HD1 | 4:D:417:VAL:HG21 | 1.84 | 0.42 |
| 4:D:1997:LYS:HE2 | 4:D:1997:LYS:HB2 | 1.85 | 0.42 |
| 12:N:172:ASP:O | 12:N:176:GLU:HB2 | 2.19 | 0.42 |
| 13:Q:124:TRP:HB2 | 13:Q:139:LEU:HD23 | 2.02 | 0.42 |
| 1:A:66:LEU:HD23 | 1:A:170:GLU:HB2 | 2.01 | 0.41 |
| 3:C:183:VAL:HB | 3:C:316:ILE:HB | 2.02 | 0.41 |
| 4:D:404:GLN:NE2 | 4:D:419:VAL:HG23 | 2.34 | 0.41 |
| 5:E:1467:LEU:HD13 | 5:E:1467:LEU:HA | 1.81 | 0.41 |
| 5:E:1547:VAL:HG23 | 5:E:1639:LEU:HD23 | 2.02 | 0.41 |
| 6:F:616:LEU:HD12 | 6:F:617:PRO:HD2 | 2.02 | 0.41 |
| 7:G:489:PRO:HG3 | 7:G:742:LEU:HD13 | 2.02 | 0.41 |
| 10:K:2085:MET:HG2 | 10:K:2086:PRO:HD2 | 2.02 | 0.41 |
| 12:N:346:GLU:H | 12:N:346:GLU:CD | 2.28 | 0.41 |
| 12:N:763:PRO:O | 12:N:764:ARG:HG3 | 2.20 | 0.41 |
| 1:A:207:LYS:HD3 | 1:A:207:LYS:HA | 1.80 | 0.41 |
| 2:B:809:MET:HE3 | 2:B:809:MET:HB3 | 1.93 | 0.41 |
| 4:D:341:ILE:HG23 | 4:D:2827:ARG:HG2 | 2.01 | 0.41 |
| 4:D:1807:PHE:HB2 | 8:H:88:LEU:HD23 | 2.02 | 0.41 |
| 5:E:1298:THR:O | 5:E:1302:LYS:HG2 | 2.20 | 0.41 |
| 5:E:1467:LEU:HG | 5:E:1489:LEU:HD13 | 2.03 | 0.41 |
| 5:E:1481:GLU:O | 5:E:1485:ALA:CB | 2.67 | 0.41 |
| 8:H:91:PRO:HB3 | 8:H:217:LEU:HD22 | 2.01 | 0.41 |
| 12:N:330:ILE:HD12 | 12:N:388:MET:HG3 | 2.02 | 0.41 |
| 3:C:496:ARG:HD3 | 5:E:1537:ARG:HD2 | 2.02 | 0.41 |
| 5:E:1353:LEU:HD13 | 5:E:1356:ARG:NH2 | 2.35 | 0.41 |
| 12:N:379:LEU:HD12 | 12:N:379:LEU:H | 1.84 | 0.41 |
| 12:N:638:GLU:HB3 | 12:N:639:ASN:CB | 2.41 | 0.41 |
| 17:Y:364:ASP:OD1 | 17:Y:364:ASP:N | 2.53 | 0.41 |
| 4:D:769:LYS:HE2 | 4:D:769:LYS:HB3 | 1.72 | 0.41 |
| 5:E:353:SER:C | 5:E:1276:MET:HE1 | 2.46 | 0.41 |
| 5:E:1481:GLU:O | 5:E:1485:ALA:HB3 | 2.21 | 0.41 |
| 8:H:423:TRP:CE2 | 8:H:468:PRO:HB3 | 2.56 | 0.41 |
| 12:N:438:LEU:HD12 | 12:N:438:LEU:HA | 1.89 | 0.41 |
| 15:S:194:LYS:NZ | 15:S:273:ASP:OD1 | 2.46 | 0.41 |
| 17:Y:239:ASN:OD1 | 17:Y:239:ASN:N | 2.50 | 0.41 |
| 1:Z:448:LYS:HA | 1:Z:448:LYS:HD3 | 1.90 | 0.41 |
| 2:B:769:MET:HB2 | 2:B:769:MET:HE2 | 1.84 | 0.41 |
| 3:C:503:TRP:CE2 | 4:D:310:VAL:HG11 | 2.55 | 0.41 |

*Continued on next page...*

For Manuscript Review

*Continued from previous page...*

| Atom-1 | Atom-2 | Interatomic distance (Å) | Clash overlap (Å) |
|---|---|---|---|
| 4:D:2190:LYS:HE2 | 4:D:2190:LYS:HB2 | 1.82 | 0.41 |
| 5:E:1355:ARG:O | 5:E:1359:LEU:HG | 2.19 | 0.41 |
| 12:N:213:HIS:CD2 | 12:N:225:ALA:HB3 | 2.55 | 0.41 |
| 12:N:499:VAL:HG12 | 12:N:517:THR:HB | 2.02 | 0.41 |
| 3:C:151:PRO:HA | 3:C:156:ASN:HD21 | 1.85 | 0.41 |
| 4:D:2313:GLU:OE1 | 4:D:2313:GLU:N | 2.47 | 0.41 |
| 5:E:1188:HIS:O | 12:N:375:ARG:NH1 | 2.53 | 0.41 |
| 7:G:390:ARG:HD2 | 7:G:390:ARG:HA | 1.95 | 0.41 |
| 12:N:205:VAL:HG21 | 12:N:211:TYR:CD2 | 2.55 | 0.41 |
| 12:N:256:LEU:HB3 | 12:N:266:LEU:HD11 | 2.02 | 0.41 |
| 17:X:158:LEU:HD12 | 17:X:158:LEU:HA | 1.96 | 0.41 |
| 2:B:468:SER:HB3 | 14:R:297:GLN:HE22 | 1.86 | 0.41 |
| 5:E:1299:GLU:O | 5:E:1303:MET:HE2 | 2.21 | 0.41 |
| 5:E:1323:LYS:HE2 | 5:E:1323:LYS:HB3 | 1.87 | 0.41 |
| 7:G:266:ASN:ND2 | 7:G:396:CYS:HB3 | 2.35 | 0.41 |
| 7:G:528:MET:HB3 | 7:G:528:MET:HE3 | 1.84 | 0.41 |
| 12:N:673:GLN:HB3 | 12:N:677:ARG:HD3 | 2.01 | 0.41 |
| 1:A:11:MET:HE1 | 10:K:1551:ALA:HA | 2.03 | 0.41 |
| 8:H:241:GLY:HA3 | 8:H:244:GLU:HG3 | 2.02 | 0.41 |
| 10:K:764:ALA:HB1 | 10:K:2385:VAL:HG21 | 2.03 | 0.41 |
| 10:K:2084:ASP:OD1 | 10:K:2084:ASP:N | 2.51 | 0.41 |
| 4:D:388:ILE:HG22 | 4:D:390:VAL:HG13 | 2.01 | 0.41 |
| 4:D:2126:MET:HE3 | 4:D:2126:MET:O | 2.21 | 0.41 |
| 7:G:589:PRO:HA | 7:G:592:LEU:HB2 | 2.01 | 0.41 |
| 8:H:367:SER:HB2 | 8:H:376:THR:HG23 | 2.03 | 0.41 |
| 8:I:491:LEU:HD23 | 8:I:491:LEU:HA | 1.94 | 0.41 |
| 10:K:1036:PRO:HD3 | 10:K:1197:LEU:HB3 | 2.03 | 0.41 |
| 10:K:2511:PRO:HG2 | 10:K:2577:ARG:HD3 | 2.02 | 0.41 |
| 12:N:335:LEU:HD23 | 12:N:384:VAL:HG13 | 2.02 | 0.41 |
| 13:Q:188:MET:HE2 | 13:Q:188:MET:HB2 | 1.93 | 0.41 |
| 13:Q:226:VAL:HG12 | 13:Q:230:GLN:HE21 | 1.85 | 0.41 |
| 16:T:304:LEU:HD23 | 16:T:304:LEU:O | 2.20 | 0.41 |
| 1:A:610:LEU:HD21 | 9:J:137:LEU:HD13 | 2.03 | 0.41 |
| 2:B:366:LYS:HA | 2:B:366:LYS:HE2 | 2.02 | 0.41 |
| 5:E:1455:ARG:HG2 | 5:E:1515:LEU:HD11 | 2.03 | 0.41 |
| 5:E:1498:GLN:HG3 | 5:E:1519:SER:HB3 | 2.02 | 0.41 |
| 8:I:119:HIS:HA | 8:I:438:TRP:O | 2.20 | 0.41 |
| 12:N:387:GLY:O | 12:N:391:ARG:HB2 | 2.21 | 0.41 |
| 13:Q:87:THR:HA | 13:Q:88:PRO:HD3 | 1.91 | 0.41 |
| 8:H:391:ARG:HE | 8:H:423:TRP:HH2 | 1.68 | 0.40 |
| 8:I:133:ALA:HB2 | 8:I:321:ALA:HB1 | 2.02 | 0.40 |

*Continued on next page...*

For Manuscript Review

*Continued from previous page...*

| Atom-1 | Atom-2 | Interatomic distance (Å) | Clash overlap (Å) |
|---|---|---|---|
| 10:K:2348:LEU:HB3 | 10:K:2360:VAL:HB | 2.04 | 0.40 |
| 11:M:1715:ALA:O | 11:M:1719:MET:HG2 | 2.21 | 0.40 |
| 12:N:193:ILE:HG21 | 12:N:236:LEU:HD12 | 2.02 | 0.40 |
| 12:N:361:LEU:HD22 | 12:N:741:GLY:HA3 | 2.03 | 0.40 |
| 1:Z:611:LYS:HB3 | 1:Z:621:TYR:HD2 | 1.87 | 0.40 |
| 1:A:119:LYS:HD3 | 10:K:1414:GLY:HA2 | 2.03 | 0.40 |
| 2:B:152:PRO:HB3 | 2:B:163:PHE:HD1 | 1.86 | 0.40 |
| 7:G:137:ARG:HH21 | 7:G:157:VAL:HG12 | 1.86 | 0.40 |
| 7:G:282:PRO:HD3 | 7:G:513:VAL:HG21 | 2.02 | 0.40 |
| 10:K:857:SER:HA | 10:K:1187:ARG:HB2 | 2.02 | 0.40 |
| 10:K:1525:PRO:HB3 | 10:K:1529:GLU:HG3 | 2.03 | 0.40 |
| 13:Q:186:MET:HA | 13:Q:187:PRO:HD3 | 1.97 | 0.40 |
| 2:B:799:VAL:HG12 | 3:C:130:ASP:HA | 2.03 | 0.40 |
| 4:D:1537:LYS:HB2 | 4:D:1537:LYS:HE2 | 1.91 | 0.40 |
| 4:D:1837:ILE:HG12 | 4:D:1884:ILE:HG12 | 2.04 | 0.40 |
| 5:E:385:ASN:ND2 | 5:E:1486:GLN:HB3 | 2.37 | 0.40 |
| 5:E:1242:ASP:HA | 12:N:363:PRO:HD3 | 2.03 | 0.40 |
| 11:M:312:GLU:H | 11:M:312:GLU:HG2 | 1.68 | 0.40 |
| 14:R:185:TRP:HB2 | 14:R:188:HIS:HB2 | 2.03 | 0.40 |
| 4:D:3058:LEU:HD23 | 4:D:3058:LEU:HA | 1.84 | 0.40 |
| 5:E:1525:LYS:HA | 5:E:1525:LYS:HD2 | 1.75 | 0.40 |
| 6:F:107:VAL:HG12 | 6:F:108:VAL:HG23 | 2.04 | 0.40 |
| 6:F:154:ASP:OD1 | 6:F:154:ASP:N | 2.47 | 0.40 |
| 6:F:339:ARG:HA | 6:F:342:GLU:OE2 | 2.21 | 0.40 |
| 3:C:152:TRP:CG | 3:C:153:GLU:H | 2.40 | 0.40 |
| 4:D:318:ILE:HD12 | 4:D:417:VAL:HG11 | 2.03 | 0.40 |
| 4:D:2852:ILE:HD12 | 4:D:3052:LYS:HB3 | 2.04 | 0.40 |
| 4:D:3052:LYS:HE3 | 4:D:3052:LYS:HB2 | 1.80 | 0.40 |
| 5:E:493:LYS:O | 5:E:497:ALA:HB3 | 2.21 | 0.40 |
| 5:E:1456:LEU:HD23 | 5:E:1456:LEU:HA | 1.91 | 0.40 |
| 5:E:1633:VAL:HG11 | 5:E:1639:LEU:HD12 | 2.04 | 0.40 |
| 10:K:1010:ALA:HB3 | 10:K:1021:LEU:HD11 | 2.02 | 0.40 |
| 16:T:270:MET:SD | 16:T:270:MET:N | 2.84 | 0.40 |

There are no symmetry-related clashes.

For Manuscript Review

#### 5.3 Torsion angles (i)

##### 5.3.1 Protein backbone (i)

In the following table, the Percentiles column shows the percent Ramachandran outliers of the chain as a percentile score with respect to all PDB entries followed by that with respect to all EM entries.

The Analysed column shows the number of residues for which the backbone conformation was analysed, and the total number of residues.

| Mol | Chain | Analysed | Favoured | Allowed | Outliers | Percentiles | |
|---|---|---|---|---|---|---|---|
| 1 | A | 405/737 (55%) | 397 (98%) | 8 (2%) | 0 | 100 | 100 |
| 1 | Z | 439/737 (60%) | 411 (94%) | 26 (6%) | 2 (0%) | 24 | 39 |
| 2 | B | 603/822 (73%) | 590 (98%) | 12 (2%) | 1 (0%) | 43 | 60 |
| 3 | C | 410/626 (66%) | 383 (93%) | 27 (7%) | 0 | 100 | 100 |
| 4 | D | 1655/3120 (53%) | 1591 (96%) | 62 (4%) | 2 (0%) | 48 | 66 |
| 5 | E | 674/1932 (35%) | 617 (92%) | 52 (8%) | 5 (1%) | 18 | 30 |
| 6 | F | 446/648 (69%) | 433 (97%) | 13 (3%) | 0 | 100 | 100 |
| 7 | G | 658/834 (79%) | 643 (98%) | 13 (2%) | 2 (0%) | 36 | 52 |
| 8 | H | 435/609 (71%) | 421 (97%) | 14 (3%) | 0 | 100 | 100 |
| 8 | I | 314/609 (52%) | 293 (93%) | 20 (6%) | 1 (0%) | 36 | 52 |
| 9 | J | 133/343 (39%) | 128 (96%) | 5 (4%) | 0 | 100 | 100 |
| 10 | K | 1704/2788 (61%) | 1658 (97%) | 44 (3%) | 2 (0%) | 48 | 66 |
| 11 | M | 357/1801 (20%) | 344 (96%) | 12 (3%) | 1 (0%) | 36 | 52 |
| 12 | N | 617/803 (77%) | 519 (84%) | 90 (15%) | 8 (1%) | 9 | 16 |
| 13 | Q | 223/414 (54%) | 222 (100%) | 1 (0%) | 0 | 100 | 100 |
| 14 | R | 299/356 (84%) | 296 (99%) | 3 (1%) | 0 | 100 | 100 |
| 15 | S | 285/359 (79%) | 285 (100%) | 0 | 0 | 100 | 100 |
| 16 | T | 98/381 (26%) | 88 (90%) | 4 (4%) | 6 (6%) | 1 | 1 |
| 17 | X | 351/410 (86%) | 342 (97%) | 9 (3%) | 0 | 100 | 100 |
| 17 | Y | 366/410 (89%) | 353 (96%) | 12 (3%) | 1 (0%) | 36 | 52 |
| All | All | 10472/18739 (56%) | 10014 (96%) | 427 (4%) | 31 (0%) | 37 | 52 |

All (31) Ramachandran outliers are listed below:

| Mol | Chain | Res | Type |
|---|---|---|---|
| 4 | D | 1433 | PRO |

*Continued on next page...*

For Manuscript Review

*Continued from previous page...*

| Mol | Chain | Res | Type |
|---|---|---|---|
| 5 | E | 508 | ALA |
| 5 | E | 1254 | ASP |
| 8 | I | 143 | PRO |
| 10 | K | 1878 | TRP |
| 11 | M | 1733 | GLU |
| 12 | N | 181 | GLU |
| 12 | N | 443 | ALA |
| 12 | N | 486 | PRO |
| 16 | T | 299 | LEU |
| 16 | T | 307 | GLN |
| 16 | T | 308 | PRO |
| 1 | Z | 130 | PRO |
| 1 | Z | 299 | PRO |
| 2 | B | 723 | VAL |
| 7 | G | 473 | GLY |
| 12 | N | 210 | VAL |
| 12 | N | 347 | GLU |
| 12 | N | 357 | VAL |
| 16 | T | 306 | GLY |
| 5 | E | 1479 | SER |
| 7 | G | 617 | ALA |
| 16 | T | 300 | SER |
| 17 | Y | 363 | ALA |
| 12 | N | 661 | VAL |
| 5 | E | 229 | ALA |
| 12 | N | 260 | VAL |
| 16 | T | 310 | LEU |
| 4 | D | 894 | ASN |
| 5 | E | 214 | SER |
| 10 | K | 1162 | PRO |

##### 5.3.2 Protein sidechains ⓘ

In the following table, the Percentiles column shows the percent sidechain outliers of the chain as a percentile score with respect to all PDB entries followed by that with respect to all EM entries.

The Analysed column shows the number of residues for which the sidechain conformation was analysed, and the total number of residues.

| Mol | Chain | Analysed | Rotameric | Outliers | Percentiles | |
|---|---|---|---|---|---|---|
| 1 | A | 384/674 (57%) | 379 (99%) | 5 (1%) | 61 | 77 |

*Continued on next page...*

For Manuscript Review

Continued from previous page...

| Mol | Chain | Analysed | Rotameric | Outliers | Percentiles | |
|---|---|---|---|---|---|---|
| 1 | Z | 405/674 (60%) | 401 (99%) | 4 (1%) | 68 | 81 |
| 2 | B | 554/737 (75%) | 546 (99%) | 8 (1%) | 59 | 76 |
| 3 | C | 364/551 (66%) | 358 (98%) | 6 (2%) | 55 | 74 |
| 4 | D | 1574/2864 (55%) | 1545 (98%) | 29 (2%) | 51 | 72 |
| 5 | E | 629/1769 (36%) | 616 (98%) | 13 (2%) | 47 | 69 |
| 6 | F | 362/499 (72%) | 359 (99%) | 3 (1%) | 73 | 85 |
| 7 | G | 526/629 (84%) | 520 (99%) | 6 (1%) | 65 | 80 |
| 8 | H | 354/473 (75%) | 352 (99%) | 2 (1%) | 78 | 88 |
| 8 | I | 264/473 (56%) | 256 (97%) | 8 (3%) | 36 | 57 |
| 9 | J | 114/277 (41%) | 112 (98%) | 2 (2%) | 51 | 72 |
| 10 | K | 1182/1841 (64%) | 1159 (98%) | 23 (2%) | 50 | 71 |
| 11 | M | 279/1296 (22%) | 276 (99%) | 3 (1%) | 65 | 80 |
| 12 | N | 495/634 (78%) | 482 (97%) | 13 (3%) | 40 | 63 |
| 13 | Q | 186/312 (60%) | 184 (99%) | 2 (1%) | 65 | 80 |
| 14 | R | 262/303 (86%) | 261 (100%) | 1 (0%) | 84 | 93 |
| 15 | S | 251/305 (82%) | 247 (98%) | 4 (2%) | 55 | 74 |
| 16 | T | 89/300 (30%) | 85 (96%) | 4 (4%) | 24 | 42 |
| 17 | X | 298/346 (86%) | 291 (98%) | 7 (2%) | 44 | 66 |
| 17 | Y | 312/346 (90%) | 310 (99%) | 2 (1%) | 78 | 88 |
| All | All | 8884/15303 (58%) | 8739 (98%) | 145 (2%) | 54 | 74 |

All (145) residues with a non-rotameric sidechain are listed below:

| Mol | Chain | Res | Type |
|---|---|---|---|
| 1 | A | 26 | SER |
| 1 | A | 112 | MET |
| 1 | A | 196 | ASN |
| 1 | A | 715 | LEU |
| 1 | A | 722 | GLU |
| 2 | B | 154 | THR |
| 2 | B | 179 | SER |
| 2 | B | 466 | SER |
| 2 | B | 485 | ASN |
| 2 | B | 490 | SER |
| 2 | B | 660 | TYR |

Continued on next page...

For Manuscript Review

*Continued from previous page...*

| Mol | Chain | Res | Type |
|---|---|---|---|
| 2 | B | 723 | VAL |
| 2 | B | 727 | THR |
| 3 | C | 206 | GLU |
| 3 | C | 232 | ILE |
| 3 | C | 444 | ASN |
| 3 | C | 460 | LEU |
| 3 | C | 497 | LEU |
| 3 | C | 502 | VAL |
| 4 | D | 205 | ASP |
| 4 | D | 252 | MET |
| 4 | D | 282 | ILE |
| 4 | D | 329 | ILE |
| 4 | D | 366 | ILE |
| 4 | D | 392 | SER |
| 4 | D | 403 | CYS |
| 4 | D | 448 | VAL |
| 4 | D | 773 | GLN |
| 4 | D | 788 | SER |
| 4 | D | 841 | GLN |
| 4 | D | 851 | ILE |
| 4 | D | 858 | LYS |
| 4 | D | 973 | GLN |
| 4 | D | 974 | LEU |
| 4 | D | 1316 | LEU |
| 4 | D | 1527 | GLN |
| 4 | D | 1530 | ILE |
| 4 | D | 1570 | HIS |
| 4 | D | 1811 | ASN |
| 4 | D | 1948 | LYS |
| 4 | D | 1971 | SER |
| 4 | D | 1999 | ASN |
| 4 | D | 2314 | LYS |
| 4 | D | 2318 | VAL |
| 4 | D | 2620 | SER |
| 4 | D | 2826 | TYR |
| 4 | D | 2845 | MET |
| 4 | D | 3115 | LEU |
| 5 | E | 1244 | LEU |
| 5 | E | 1245 | ILE |
| 5 | E | 1297 | PHE |
| 5 | E | 1342 | GLN |
| 5 | E | 1344 | TYR |

*Continued on next page...*

For Manuscript Review

Continued from previous page...

| Mol | Chain | Res | Type |
|---|---|---|---|
| 5 | E | 1428 | LEU |
| 5 | E | 1430 | LEU |
| 5 | E | 1431 | LEU |
| 5 | E | 1522 | LEU |
| 5 | E | 1612 | VAL |
| 5 | E | 1657 | MET |
| 5 | E | 1694 | SER |
| 5 | E | 1903 | PHE |
| 6 | F | 281 | MET |
| 6 | F | 333 | GLU |
| 6 | F | 493 | VAL |
| 7 | G | 71 | LEU |
| 7 | G | 119 | GLU |
| 7 | G | 143 | VAL |
| 7 | G | 543 | THR |
| 7 | G | 582 | GLU |
| 7 | G | 615 | GLU |
| 8 | H | 251 | LEU |
| 8 | H | 453 | ASP |
| 8 | I | 116 | VAL |
| 8 | I | 266 | LYS |
| 8 | I | 325 | ASP |
| 8 | I | 327 | LEU |
| 8 | I | 353 | GLN |
| 8 | I | 369 | GLN |
| 8 | I | 451 | ASN |
| 8 | I | 486 | HIS |
| 9 | J | 172 | ASP |
| 9 | J | 246 | VAL |
| 10 | K | 644 | GLU |
| 10 | K | 787 | THR |
| 10 | K | 1197 | LEU |
| 10 | K | 1340 | VAL |
| 10 | K | 1450 | GLU |
| 10 | K | 1457 | ARG |
| 10 | K | 1506 | LEU |
| 10 | K | 1521 | VAL |
| 10 | K | 1522 | ARG |
| 10 | K | 1524 | LEU |
| 10 | K | 1538 | THR |
| 10 | K | 1544 | ILE |
| 10 | K | 1725 | ASP |

Continued on next page...

For Manuscript Review

Continued from previous page...

| Mol | Chain | Res | Type |
|---|---|---|---|
| 10 | K | 1838 | LEU |
| 10 | K | 1878 | TRP |
| 10 | K | 2034 | LEU |
| 10 | K | 2121 | LEU |
| 10 | K | 2285 | GLU |
| 10 | K | 2361 | ARG |
| 10 | K | 2382 | MET |
| 10 | K | 2451 | VAL |
| 10 | K | 2464 | MET |
| 10 | K | 2513 | CYS |
| 11 | M | 415 | MET |
| 11 | M | 473 | LEU |
| 11 | M | 1664 | ARG |
| 12 | N | 174 | PHE |
| 12 | N | 195 | VAL |
| 12 | N | 221 | ASP |
| 12 | N | 235 | VAL |
| 12 | N | 236 | LEU |
| 12 | N | 266 | LEU |
| 12 | N | 285 | GLU |
| 12 | N | 370 | GLN |
| 12 | N | 487 | VAL |
| 12 | N | 585 | THR |
| 12 | N | 592 | MET |
| 12 | N | 648 | VAL |
| 12 | N | 747 | VAL |
| 13 | Q | 151 | THR |
| 13 | Q | 186 | MET |
| 14 | R | 132 | SER |
| 15 | S | 116 | ILE |
| 15 | S | 120 | GLU |
| 15 | S | 186 | TRP |
| 15 | S | 319 | GLU |
| 16 | T | 194 | LEU |
| 16 | T | 270 | MET |
| 16 | T | 303 | ARG |
| 16 | T | 310 | LEU |
| 17 | X | 59 | SER |
| 17 | X | 101 | MET |
| 17 | X | 182 | VAL |
| 17 | X | 229 | SER |
| 17 | X | 310 | ILE |

Continued on next page...

For Manuscript Review

*Continued from previous page...*

| Mol | Chain | Res | Type |
|---|---|---|---|
| 17 | X | 401 | TYR |
| 17 | X | 407 | ASP |
| 17 | Y | 243 | ASN |
| 17 | Y | 268 | GLU |
| 1 | Z | 117 | THR |
| 1 | Z | 290 | SER |
| 1 | Z | 297 | LEU |
| 1 | Z | 587 | ASN |

Sometimes sidechains can be flipped to improve hydrogen bonding and reduce clashes. All (66) such sidechains are listed below:

| Mol | Chain | Res | Type |
|---|---|---|---|
| 1 | A | 151 | GLN |
| 1 | A | 169 | ASN |
| 1 | A | 177 | GLN |
| 1 | A | 617 | ASN |
| 2 | B | 469 | GLN |
| 2 | B | 485 | ASN |
| 2 | B | 517 | GLN |
| 2 | B | 743 | GLN |
| 2 | B | 780 | GLN |
| 3 | C | 93 | GLN |
| 3 | C | 205 | ASN |
| 3 | C | 444 | ASN |
| 4 | D | 132 | ASN |
| 4 | D | 135 | GLN |
| 4 | D | 242 | ASN |
| 4 | D | 386 | ASN |
| 4 | D | 601 | GLN |
| 4 | D | 766 | ASN |
| 4 | D | 1970 | ASN |
| 4 | D | 2035 | GLN |
| 4 | D | 2054 | ASN |
| 4 | D | 2179 | GLN |
| 4 | D | 2317 | ASN |
| 4 | D | 2361 | ASN |
| 4 | D | 2364 | GLN |
| 4 | D | 2461 | HIS |
| 4 | D | 2809 | GLN |
| 5 | E | 378 | HIS |
| 5 | E | 385 | ASN |
| 5 | E | 501 | GLN |

*Continued on next page...*

For Manuscript Review

*Continued from previous page...*

| Mol | Chain | Res | Type |
|---|---|---|---|
| 5 | E | 505 | ASN |
| 5 | E | 1185 | ASN |
| 5 | E | 1210 | ASN |
| 5 | E | 1241 | ASN |
| 5 | E | 1306 | GLN |
| 5 | E | 1365 | ASN |
| 5 | E | 1458 | GLN |
| 5 | E | 1531 | GLN |
| 5 | E | 1601 | GLN |
| 5 | E | 1656 | GLN |
| 5 | E | 1660 | HIS |
| 5 | E | 1724 | HIS |
| 7 | G | 150 | HIS |
| 7 | G | 165 | GLN |
| 7 | G | 227 | GLN |
| 8 | H | 147 | ASN |
| 8 | I | 392 | HIS |
| 10 | K | 1277 | HIS |
| 12 | N | 213 | HIS |
| 12 | N | 232 | GLN |
| 12 | N | 380 | ASN |
| 12 | N | 382 | GLN |
| 12 | N | 687 | GLN |
| 14 | R | 88 | GLN |
| 14 | R | 138 | HIS |
| 14 | R | 294 | HIS |
| 15 | S | 105 | ASN |
| 16 | T | 269 | ASN |
| 16 | T | 283 | ASN |
| 17 | X | 145 | GLN |
| 17 | X | 245 | HIS |
| 17 | X | 328 | GLN |
| 17 | Y | 273 | GLN |
| 1 | Z | 6 | ASN |
| 1 | Z | 30 | ASN |
| 1 | Z | 592 | GLN |

##### 5.3.3 RNA ⓘ

There are no RNA molecules in this entry.

For Manuscript Review

#### 5.4 Non-standard residues in protein, DNA, RNA chains (i)

There are no non-standard protein/DNA/RNA residues in this entry.

#### 5.5 Carbohydrates (i)

There are no oligosaccharides in this entry.

#### 5.6 Ligand geometry (i)

There are no ligands in this entry.

#### 5.7 Other polymers (i)

There are no such residues in this entry.

#### 5.8 Polymer linkage issues (i)

The following chains have linkage breaks:

| Mol | Chain | Number of breaks |
|---|---|---|
| 8 | I | 1 |

All chain breaks are listed below:

| Model | Chain | Residue-1 | Atom-1 | Residue-2 | Atom-2 | Distance (Å) |
|---|---|---|---|---|---|---|
| 1 | I | 501:ALA | C | 502:ASP | N | 4.51 |

For Manuscript Review

### 6 Map visualisation (i)

This section contains visualisations of the EMDB entry EMD-56766. These allow visual inspection of the internal detail of the map and identification of artifacts.

No raw map or half-maps were deposited for this entry and therefore no images, graphs, etc. pertaining to the raw map can be shown.

#### 6.1 Orthogonal projections (i)

##### 6.1.1 Primary map

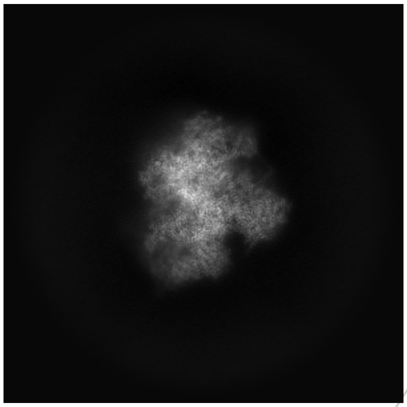

X

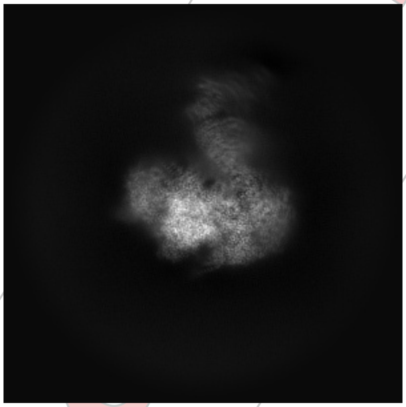

Y

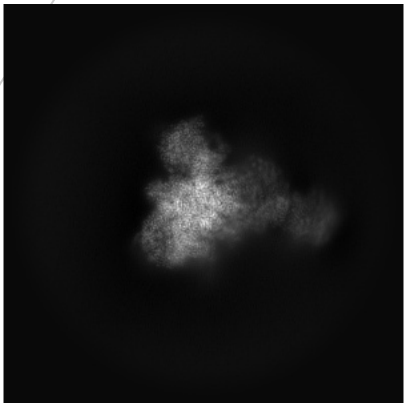

Z

The images above show the map projected in three orthogonal directions.

#### 6.2 Central slices (i)

##### 6.2.1 Primary map

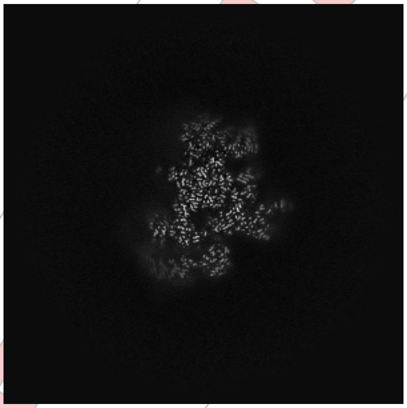

X Index: 294

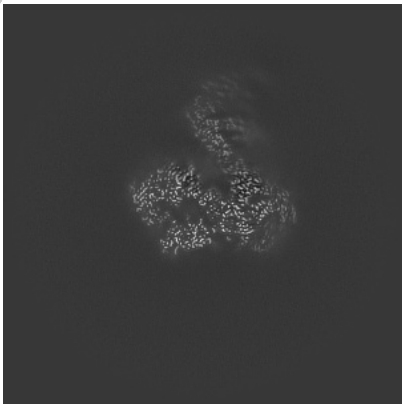

Y Index: 294

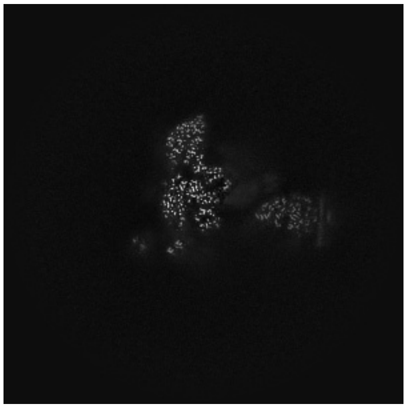

Z Index: 294

For Manuscript Review

The images above show central slices of the map in three orthogonal directions.

#### 6.3 Largest variance slices (i)

##### 6.3.1 Primary map

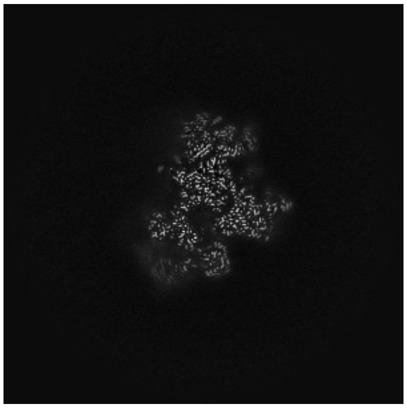
X Index: 289

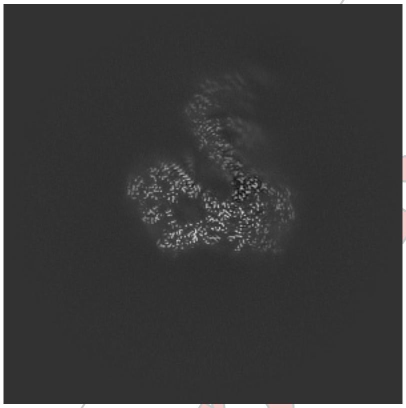
Y Index: 288

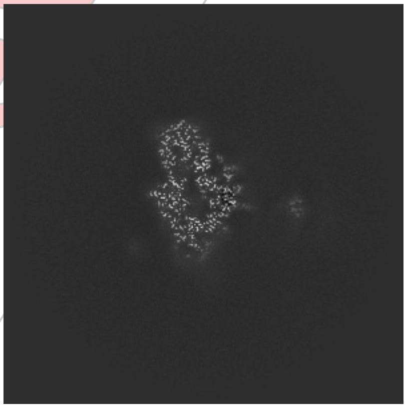
Z Index: 272

The images above show the largest variance slices of the map in three orthogonal directions.

#### 6.4 Orthogonal standard-deviation projections (False-color) (i)

##### 6.4.1 Primary map

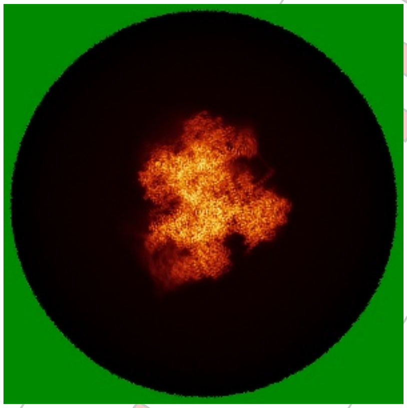
X

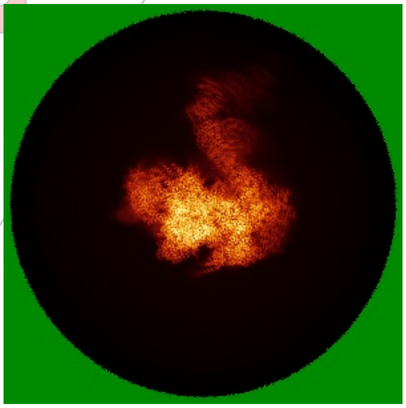
Y

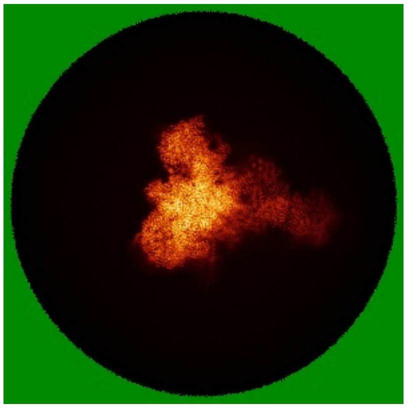
Z

The images above show the map standard deviation projections with false color in three orthogonal directions. Minimum values are shown in green, max in blue, and dark to light orange shades represent small to large values respectively.

For Manuscript Review

#### 6.5 Orthogonal surface views (i)

##### 6.5.1 Primary map

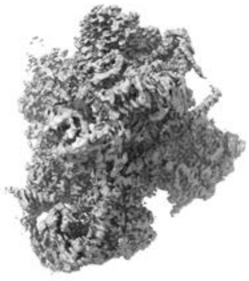

X

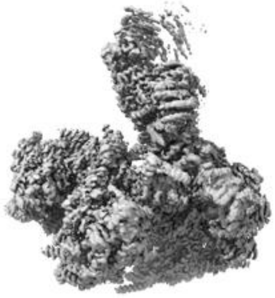

Y

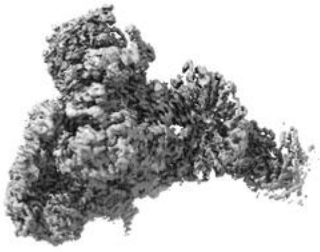

Z

The images above show the 3D surface view of the map at the recommended contour level 0.235. These images, in conjunction with the slice images, may facilitate assessment of whether an appropriate contour level has been provided.

#### 6.6 Mask visualisation (i)

This section was not generated. No masks/segmentation were deposited.

For Manuscript Review

### 7 Map analysis (i)

This section contains the results of statistical analysis of the map.

#### 7.1 Map-value distribution (i)

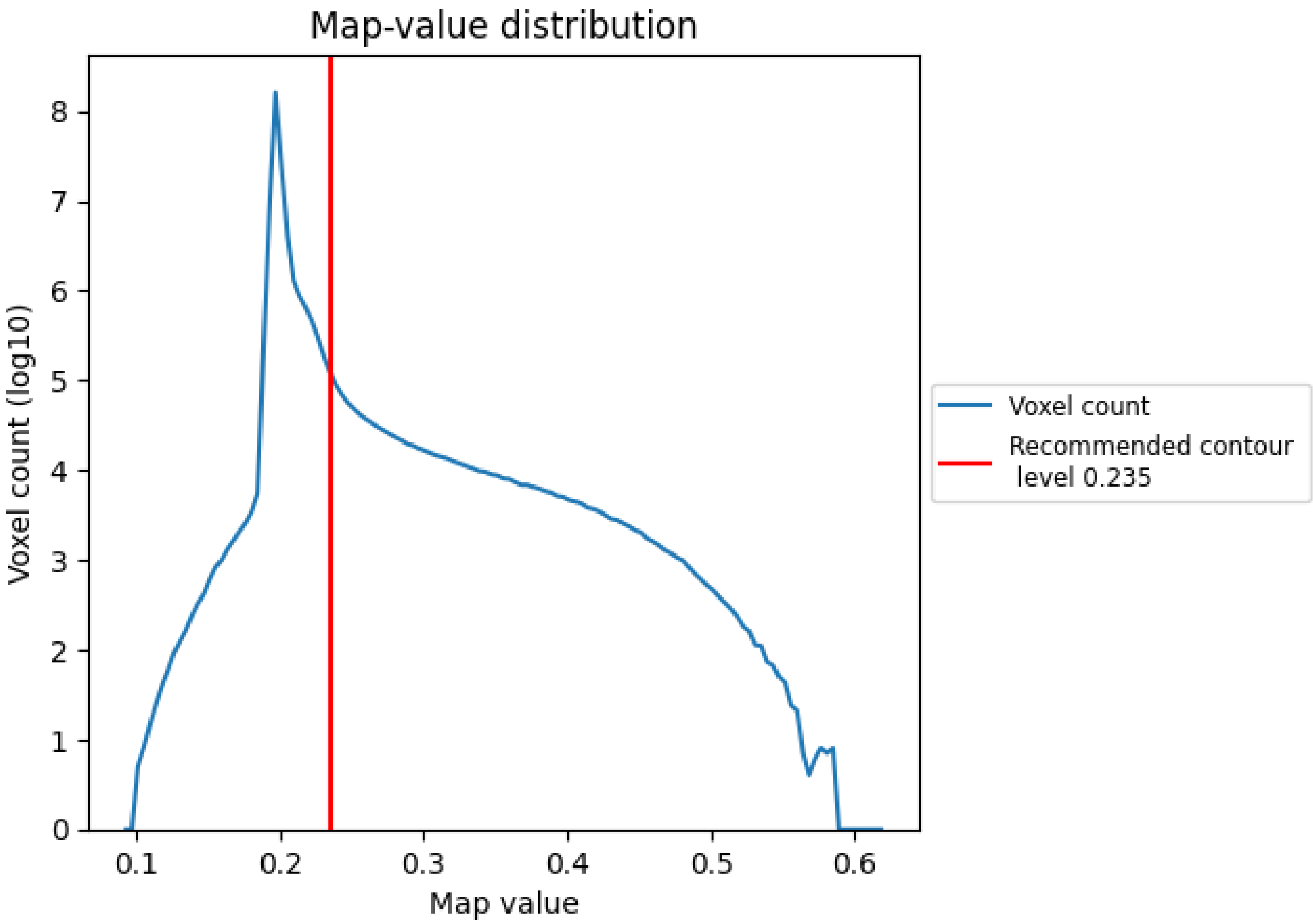


The map-value distribution is plotted in 128 intervals along the x-axis. The y-axis is logarithmic. A spike in this graph at zero usually indicates that the volume has been masked.

For Manuscript Review

#### 7.2 Volume estimate (i)

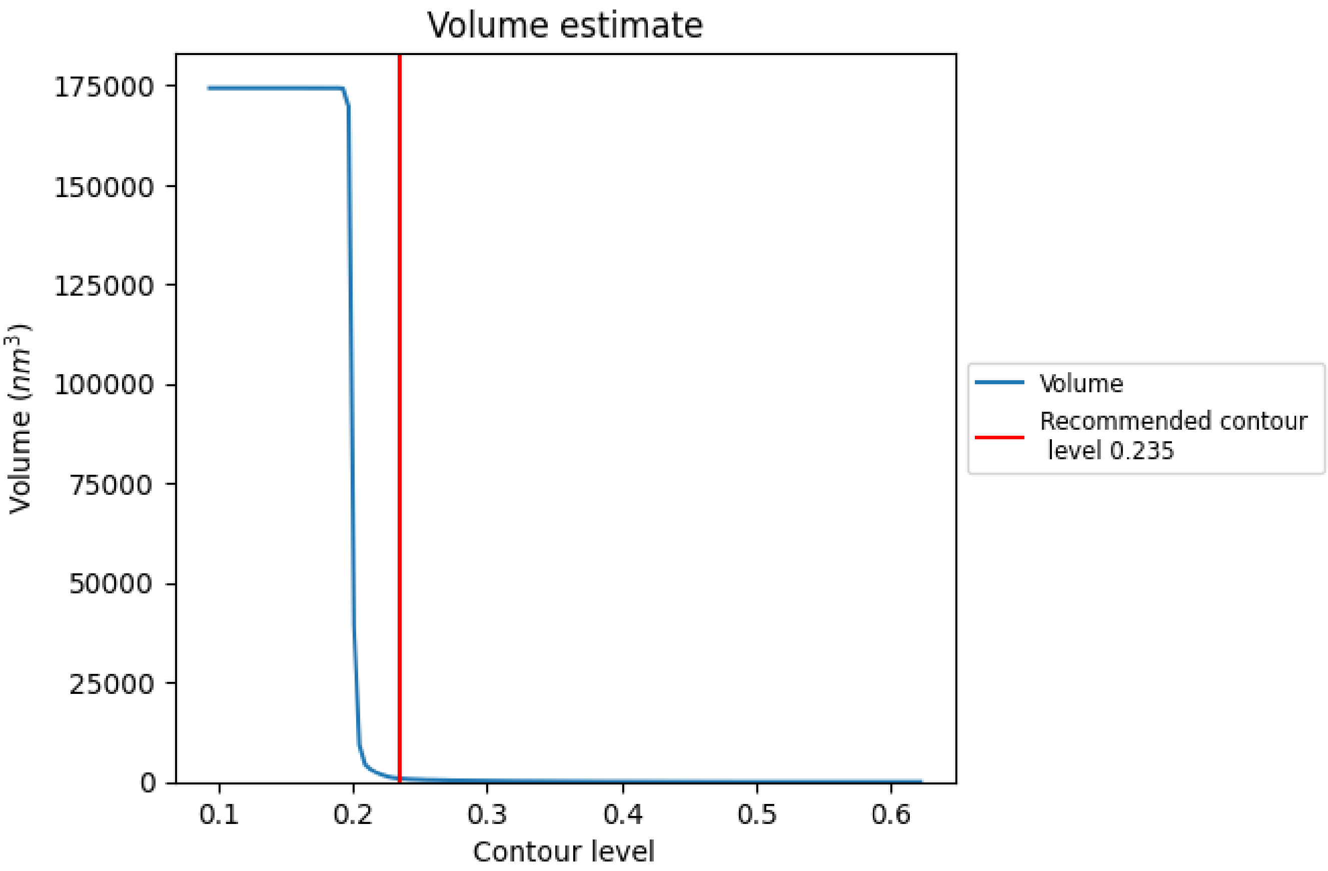


The volume at the recommended contour level is 833 $nm^3$; this corresponds to an approximate mass of 752 kDa.

The volume estimate graph shows how the enclosed volume varies with the contour level. The recommended contour level is shown as a vertical line and the intersection between the line and the curve gives the volume of the enclosed surface at the given level.

For Manuscript Review

#### 7.3 Rotationally averaged power spectrum (i)

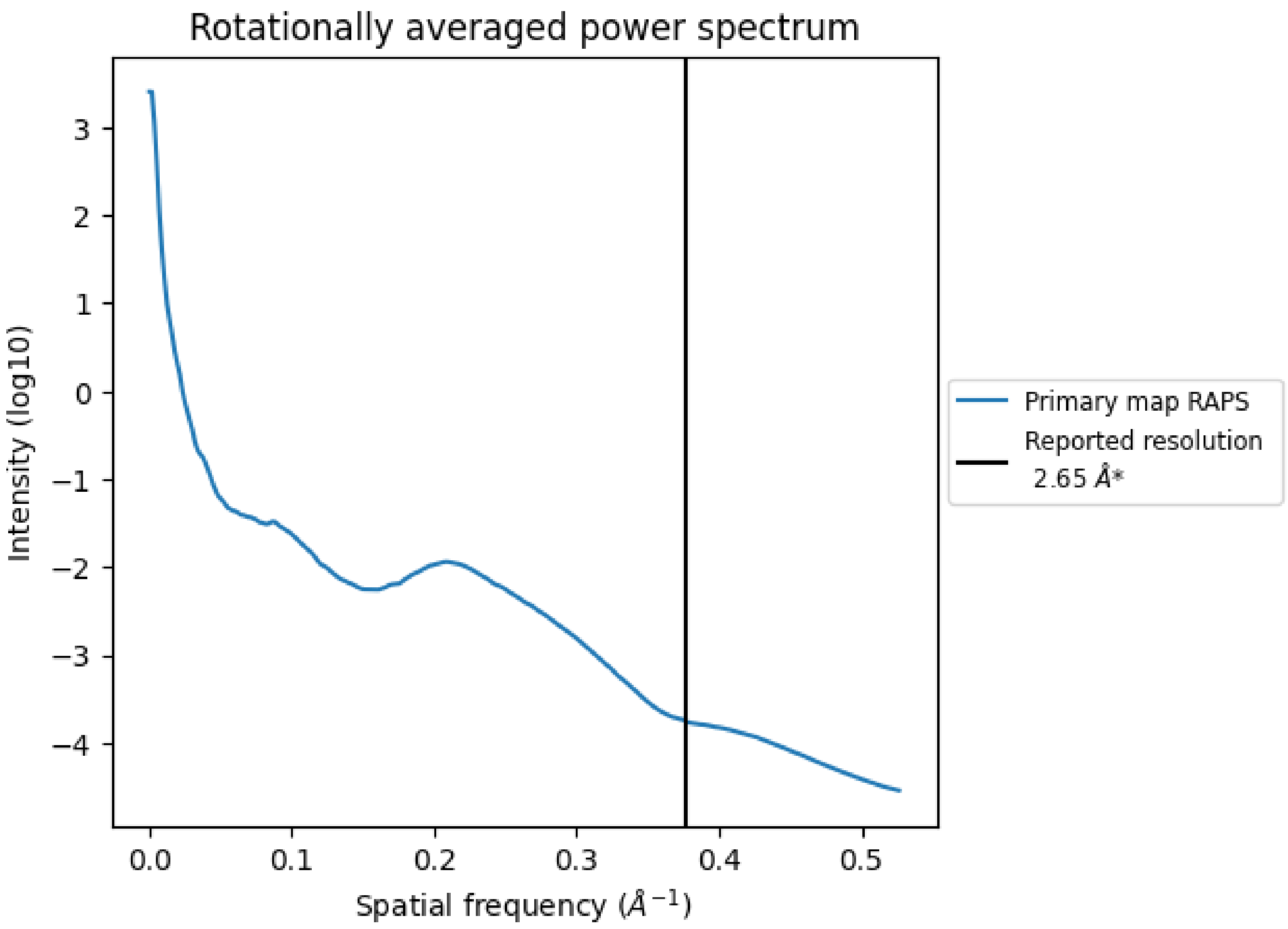


*Reported resolution corresponds to spatial frequency of 0.377 Å$^{-1}$

For Manuscript Review

### 8 Fourier-Shell correlation (i)

This section was not generated. No FSC curve or half-maps provided.

For Manuscript Review

### 9 Map-model fit (i)

This section contains information regarding the fit between EMDB map EMD-56766 and PDB model 28RF. Per-residue inclusion information can be found in section 3 on page 7.

#### 9.1 Map-model overlay (i)

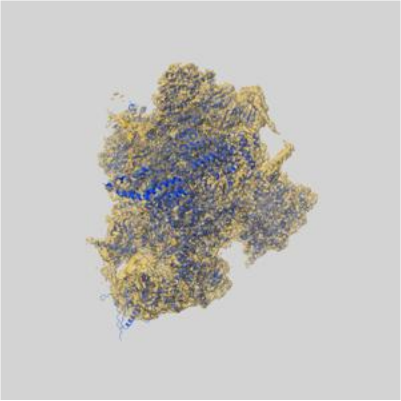

X

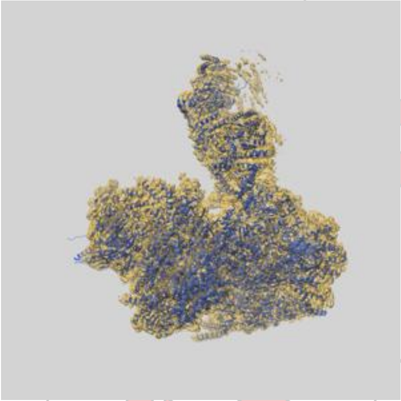

Y

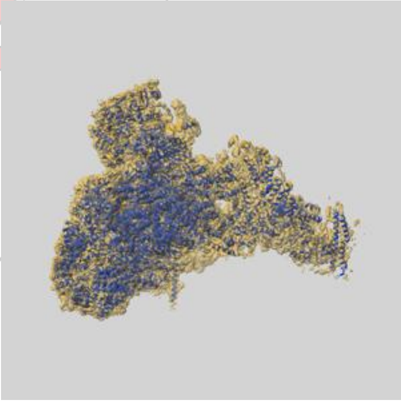

Z

The images above show the 3D surface view of the map at the recommended contour level 0.235 at 50% transparency in yellow overlaid with a ribbon representation of the model coloured in blue. These images allow for the visual assessment of the quality of fit between the atomic model and the map.

For Manuscript Review

#### 9.2 Q-score mapped to coordinate model (i)

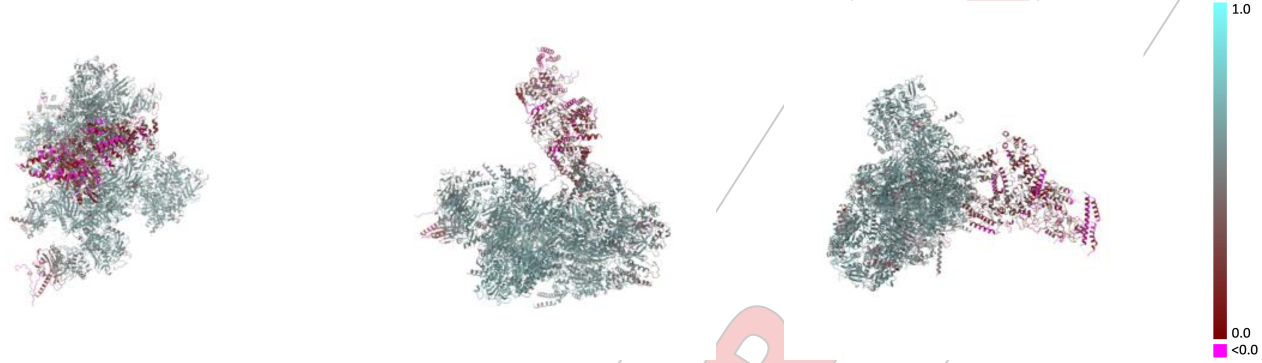


X Y Z

The images above show the model with each residue coloured according its Q-score. This shows their resolvability in the map with higher Q-score values reflecting better resolvability. Please note: Q-score is calculating the resolvability of atoms, and thus high values are only expected at resolutions at which atoms can be resolved. Low Q-score values may therefore be expected for many entries.

#### 9.3 Atom inclusion mapped to coordinate model (i)

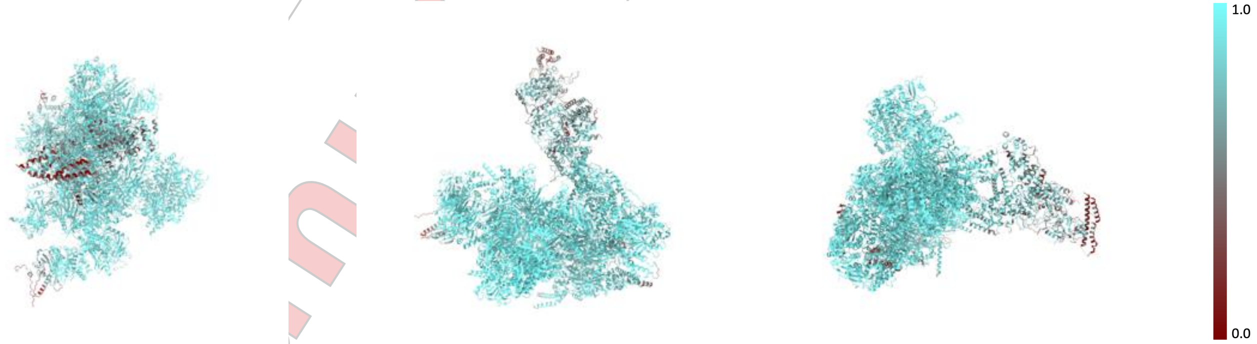


X Y Z

The images above show the model with each residue coloured according to its atom inclusion. This shows to what extent they are inside the map at the recommended contour level (0.235).

For Manuscript Review

#### 9.4 Atom inclusion (i)

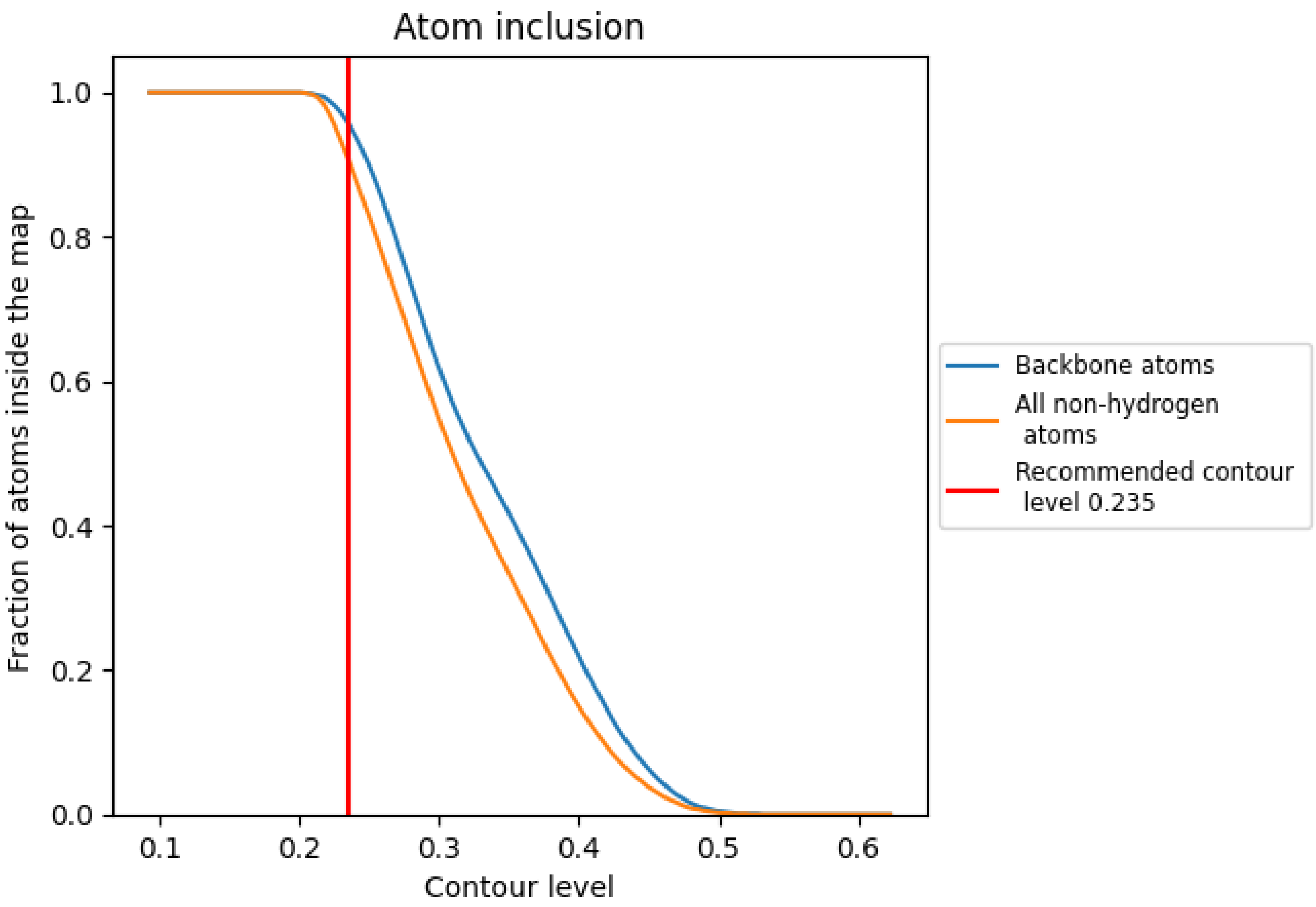


At the recommended contour level, 96% of all backbone atoms, 91% of all non-hydrogen atoms, are inside the map.

For Manuscript Review

#### 9.5 Map-model fit summary (i)

The table lists the average atom inclusion at the recommended contour level (0.235) and Q-score for the entire model and for each chain.

| Chain | Atom inclusion | Q-score |
|---|---|---|
| All | 0.9070 | 0.5260 |
| A | 0.9480 | 0.5850 |
| B | 0.9820 | 0.6160 |
| C | 0.9620 | 0.5870 |
| D | 0.9490 | 0.5580 |
| E | 0.8150 | 0.3960 |
| F | 0.9660 | 0.5960 |
| G | 0.9740 | 0.5870 |
| H | 0.9690 | 0.5760 |
| I | 0.9520 | 0.5230 |
| J | 0.9480 | 0.5540 |
| K | 0.9430 | 0.5740 |
| M | 0.5210 | 0.1720 |
| N | 0.6520 | 0.2340 |
| Q | 0.5390 | 0.2210 |
| R | 0.9670 | 0.6100 |
| S | 0.9610 | 0.6080 |
| T | 0.9520 | 0.4880 |
| X | 0.9900 | 0.6350 |
| Y | 0.9870 | 0.6190 |
| Z | 0.9060 | 0.5560 |

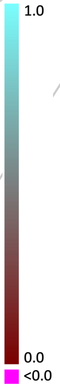


For Manuscript Review
