## Supplementary figures and images for "The structure of a 2-MDa chloroplast RNA polymerase reveals unexpected evolutionary complexity"

### Cre05.g234651.t1.1.hits.fa.mafft.treefile_color.tre.pdf

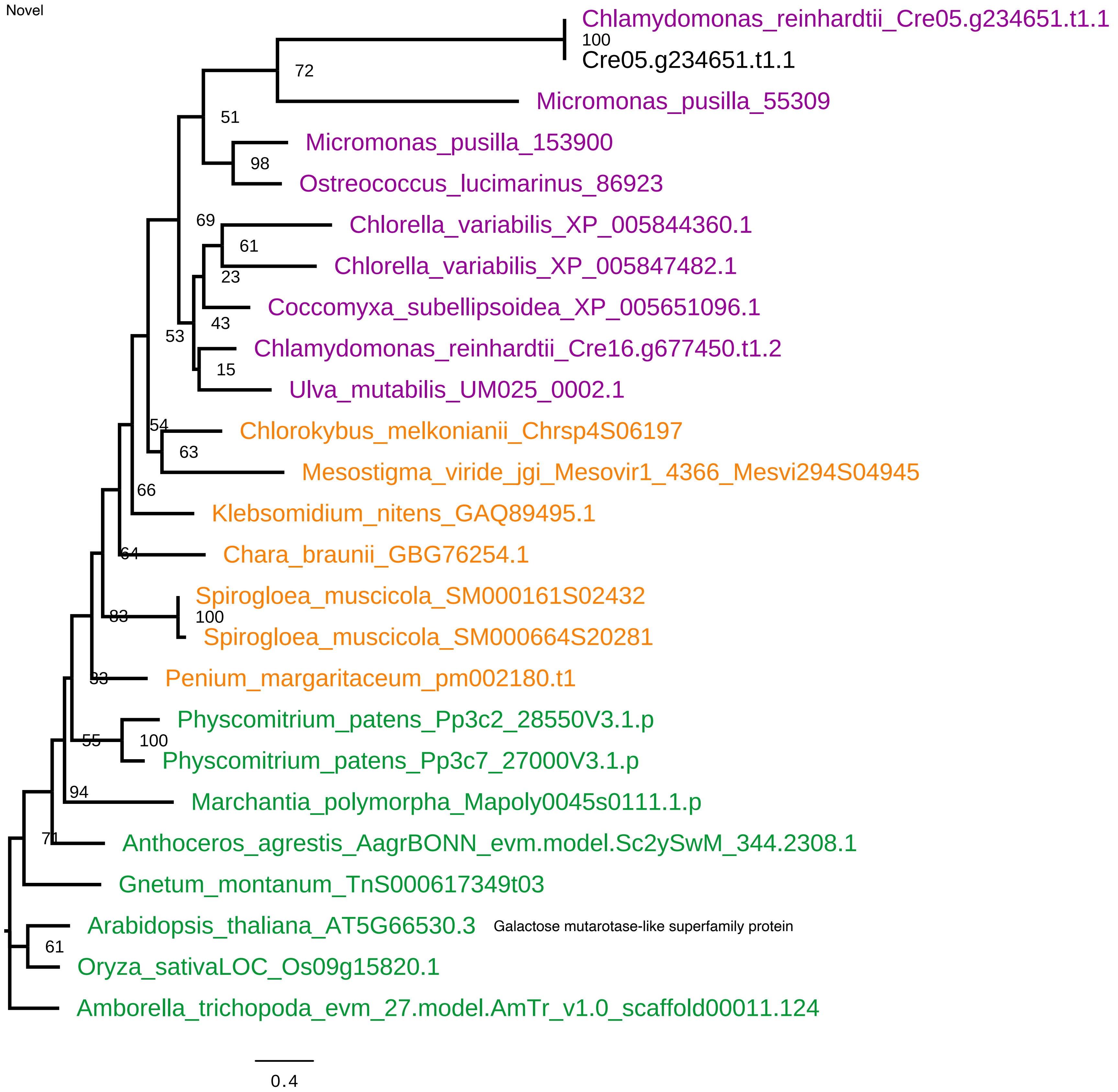

### Cre07.g348500.t1.1.hits.fa.mafft.treefile_color.tre.pdf

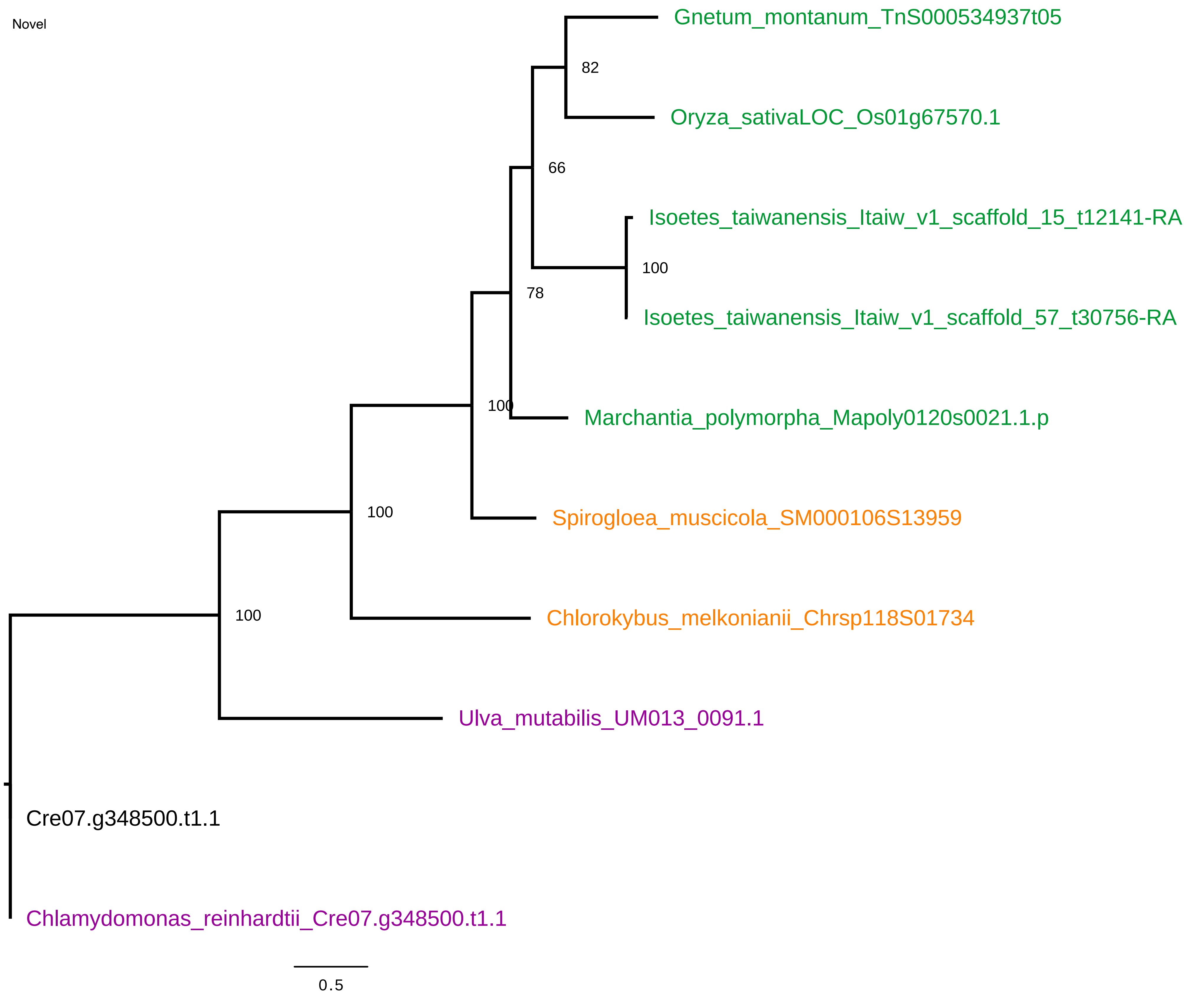

### Cre09.g400627.t1.1.hits.fa.mafft.treefile_color.tre.pdf

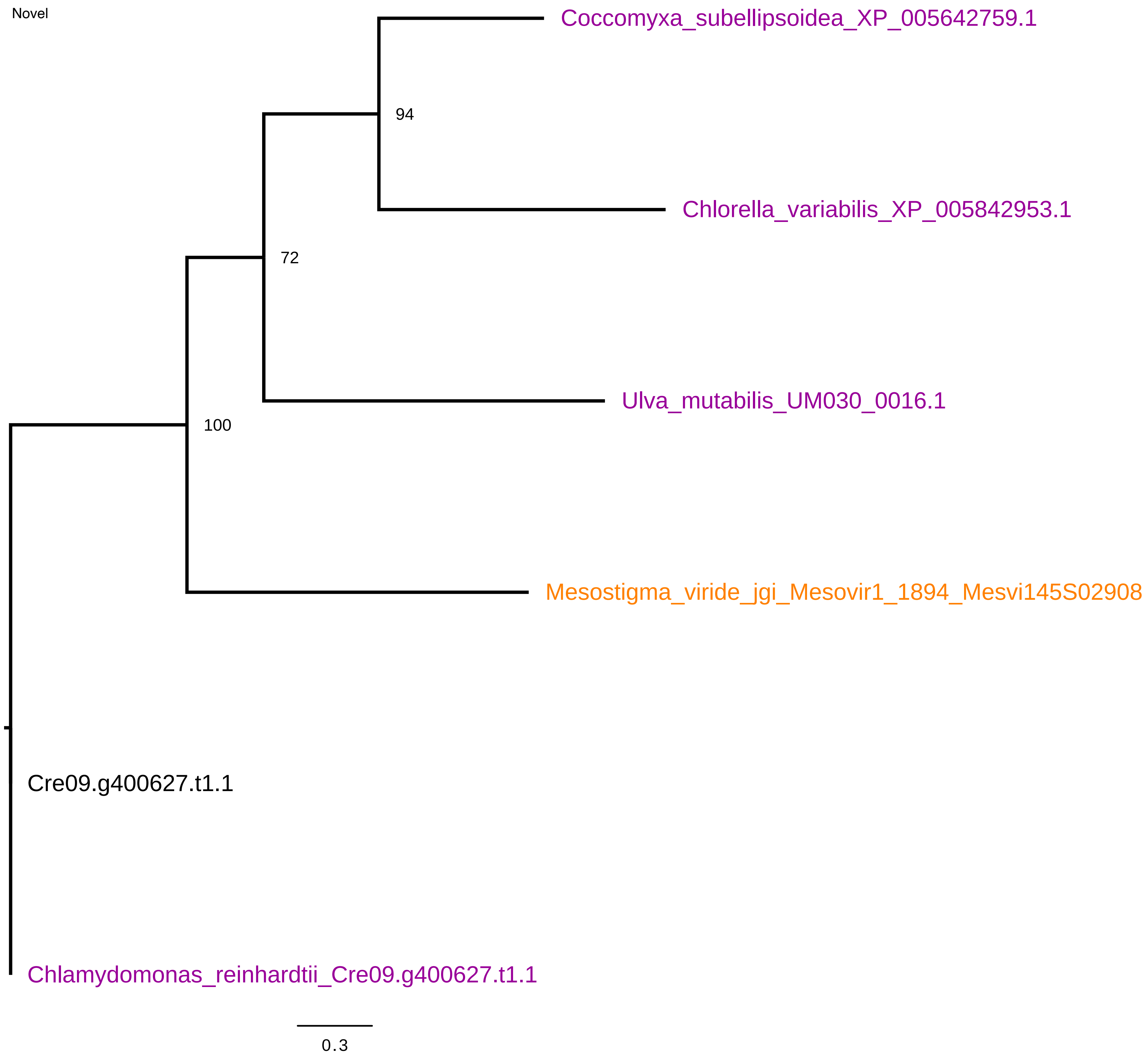

### Cre12.g497350.t1.1.hits.fa.mafft.treefile_color.tre.pdf

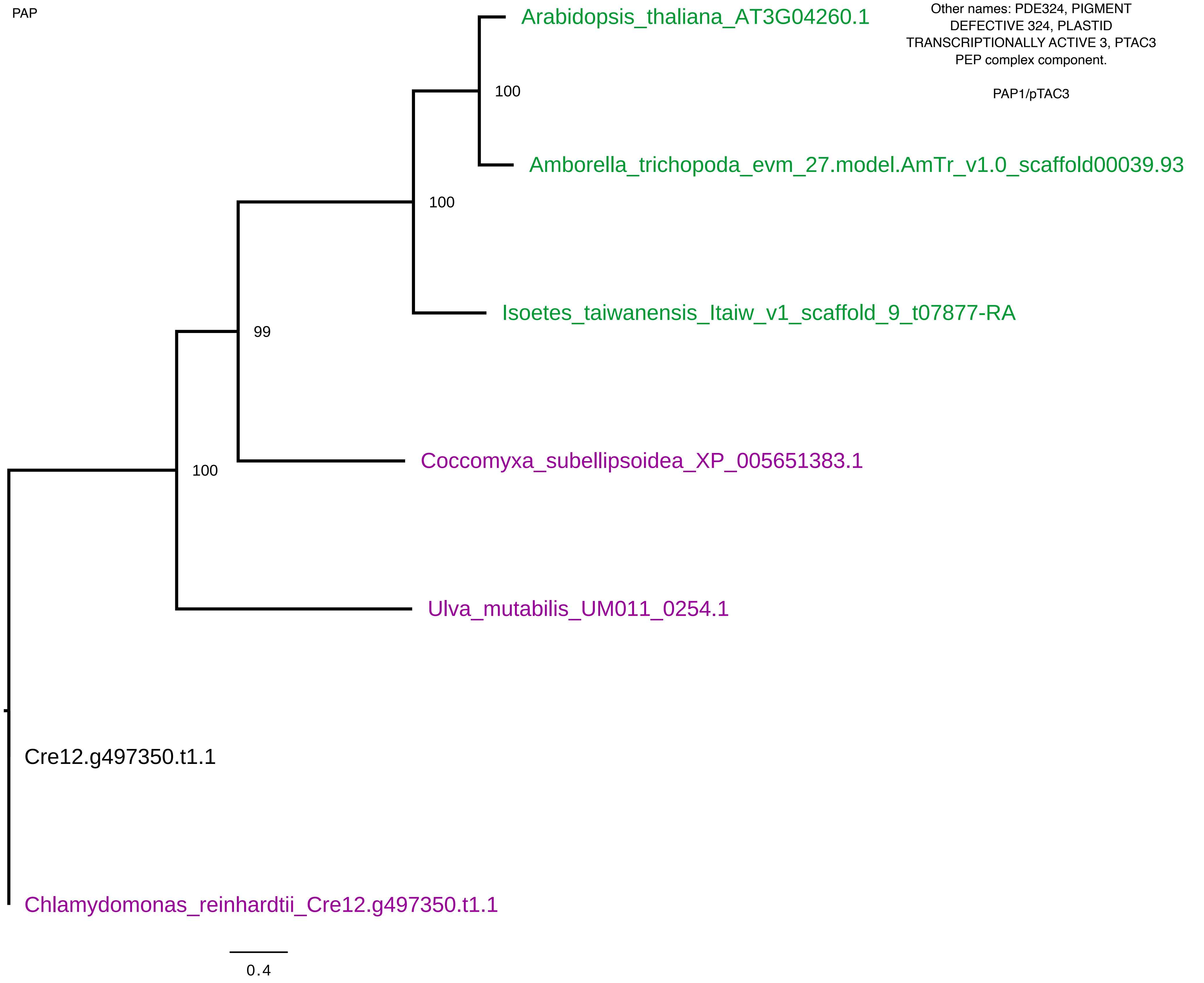

### Cre12.g519900.t1.2.hits.fa.mafft.treefile_color.tre.pdf

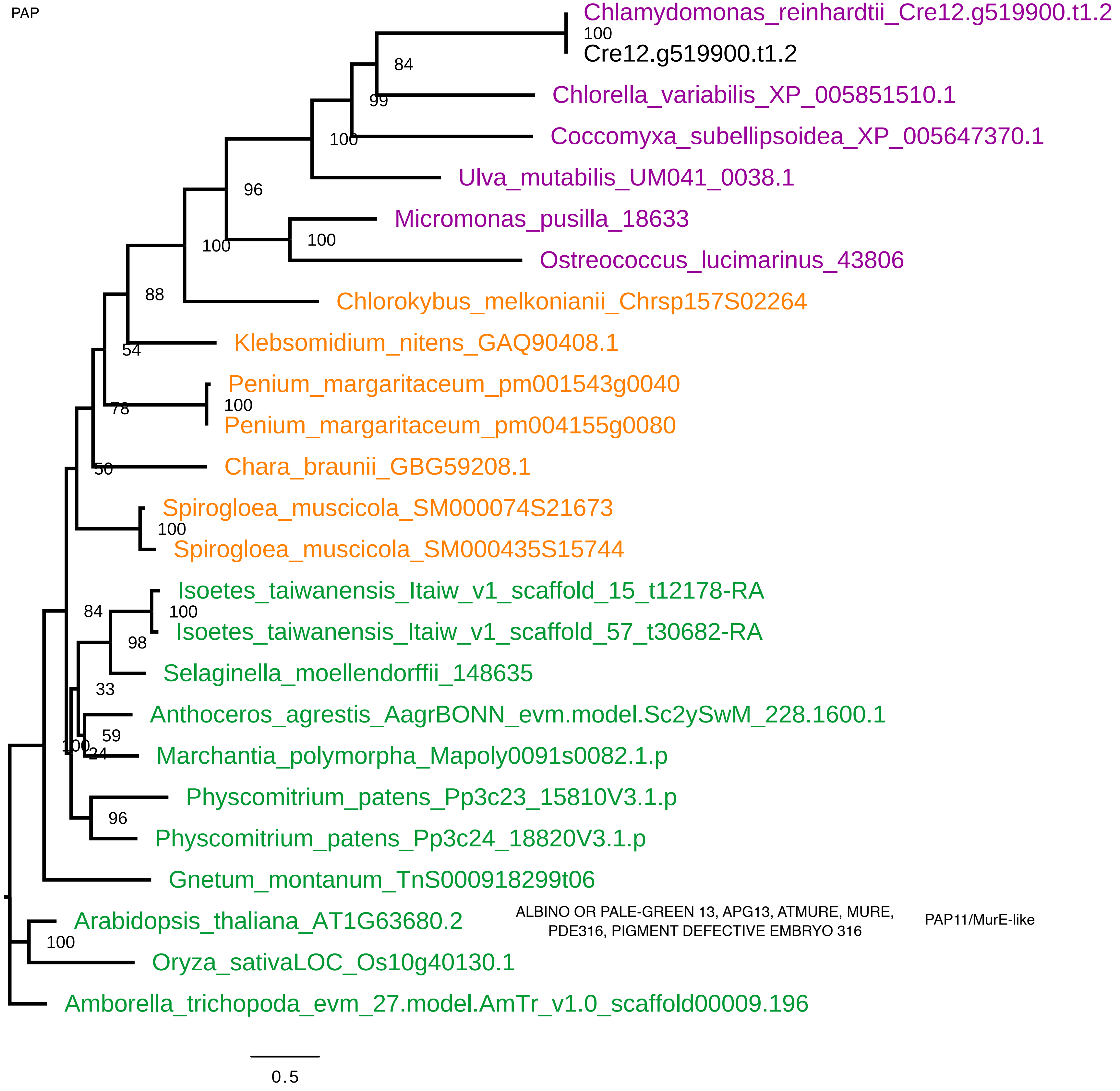

### Cre13.g576400.t1.2.hits.fa.mafft.treefile_color.tre.pdf

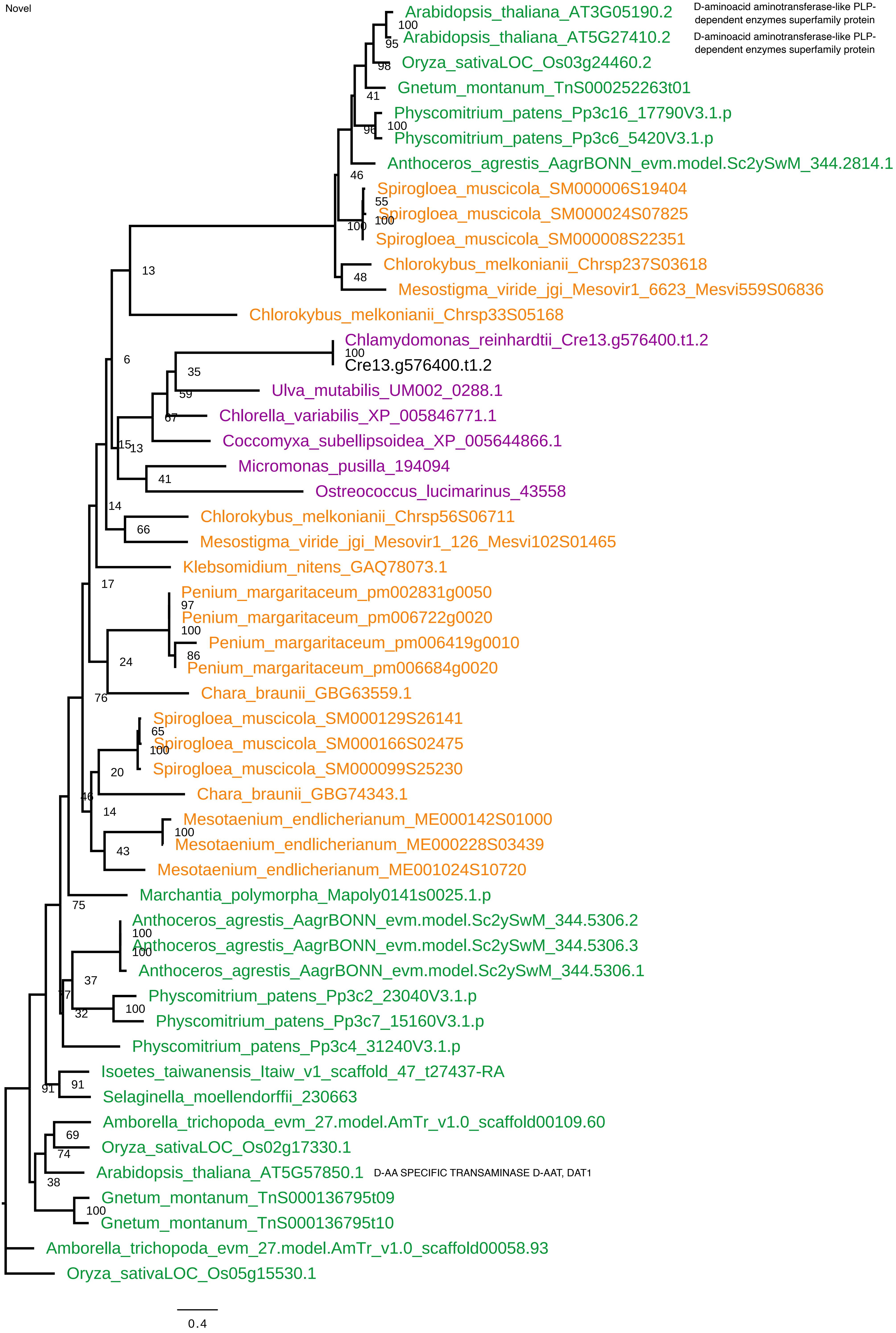

### Cre15.g635650.t1.2.hits.fa.mafft.treefile_color.tre.pdf

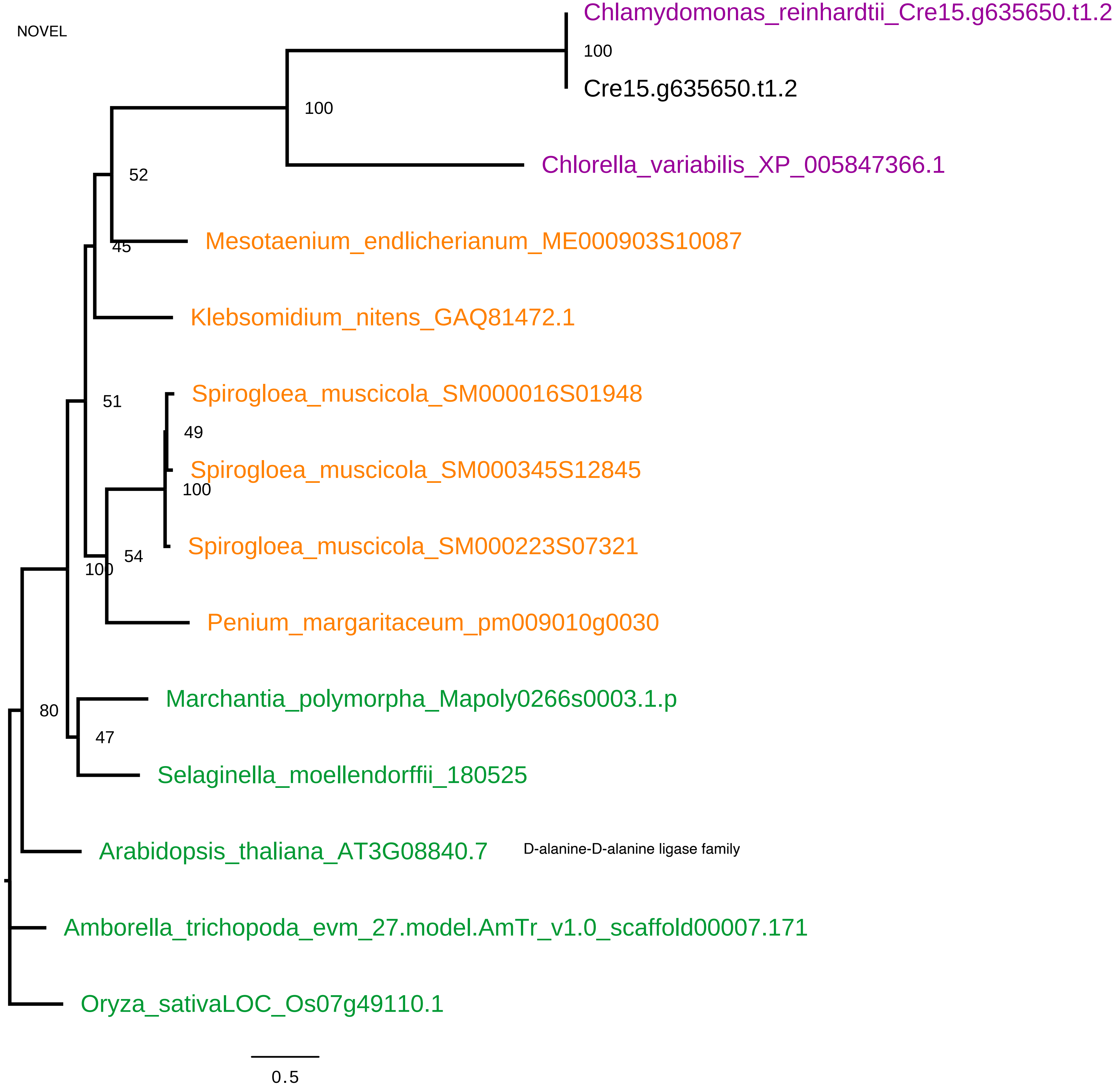

### Cre16.g653350.t1.2.hits.fa.mafft.treefile_color.tre.pdf

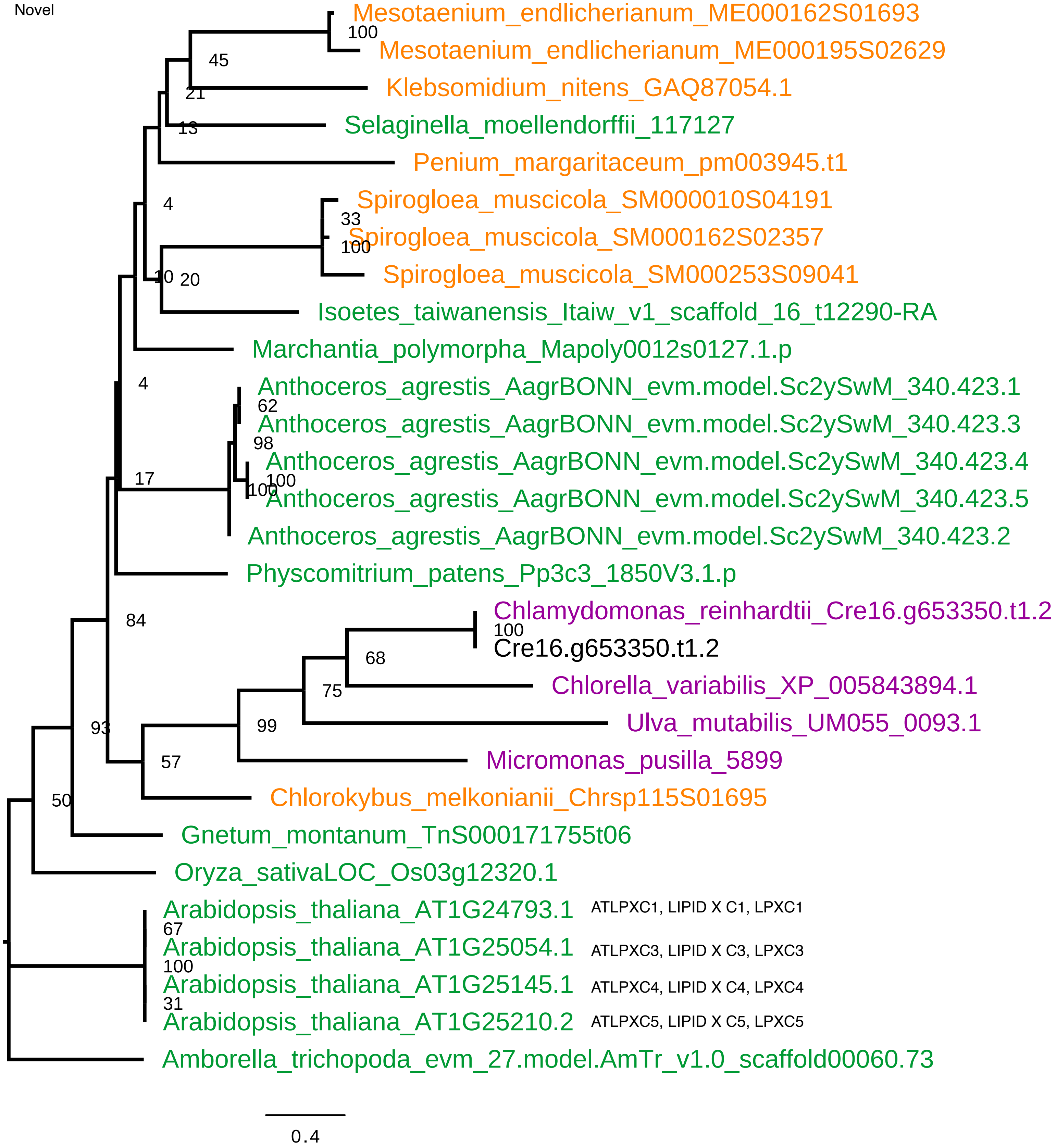
